# A JAK/STAT-Mediated Inflammatory Signaling Cascade Drives Oncogenesis In AF10-Rearranged AML

**DOI:** 10.1101/2020.08.31.273631

**Authors:** Bo-Rui Chen, Anagha Deshpande, Karina Barbosa, Maria Kleppe, Xue Li, Narayana Yeddula, Alexandre Rosa Campos, Robert J. Wechsler-Reya, Anindya Bagchi, Soheil Meshinchi, Connie Eaves, Irmela Jeremias, Torsten Haferlach, David A Frank, Ze’ev Ronai, Sumit Chanda, Scott A. Armstrong, Peter D. Adams, Ross L. Levine, Aniruddha J. Deshpande

**Affiliations:** Tumor Initiation and Maintenance Program, Sanford Burnham Prebys Medical Discovery Institute, La Jolla, CA, 92037, USA; Human Oncology and Pathogenesis Program, Memorial Sloan Kettering Cancer Center, New York, NY, 10065, USA; Immunity and Pathogenesis Program, Infectious and Inflammatory Disease Center, Sanford Burnham Prebys Medical Discovery Institute, La Jolla, CA, 92037, USA; Proteomics Facility, Sanford Burnham Prebys Medical Discovery Institute, La Jolla, CA, 92037, USA; Clinical Research Division, Fred Hutchinson Cancer Research Center, Seattle, Washington, USA; Terry Fox Laboratory, British Columbia Cancer Agency, Vancouver, British Columbia, Canada; MLL Munich Leukemia Laboratory, Munich, 81377, Germany; Research Unit Apoptosis in Hematopoietic Stem Cells, Helmholtz Center Munich, German Center for Environmental Health, Munich, Germany; Department of Medical Oncology, Dana Farber Cancer Institute, Boston, MA, 02215, USA; Department of Pediatric Oncology, Dana-Farber Cancer Institute, Boston, MA, 02215, USA

**Keywords:** PICALM-AF10, MLL-AF10, MLLT10, AML, leukemia, JAK1, STAT3, atovaquone, inflammatory signaling

## Abstract

Leukemias bearing fusions of the AF10/MLLT10 gene are associated with poor prognosis, and therapies targeting these fusion proteins are lacking. To understand mechanisms underlying AF10 fusion-mediated leukemogenesis, we generated inducible mouse models of AML driven by the most common AF10 fusion proteins, PICALM/CALM-AF10 and KMT2A/MLL-AF10, and performed comprehensive characterization of the disease using transcriptomic, epigenomic, proteomic, and functional genomic approaches. Our studies provide a comprehensive map of gene networks and protein interactors associated with key AF10 fusions involved in leukemia. Specifically, we report that AF10 fusions activate a cascade of JAK/STAT-mediated inflammatory signaling through direct recruitment of JAK1 kinase. Inhibition of the JAK/STAT signaling by genetic *Jak1* deletion or through pharmacological JAK/STAT inhibition elicited potent anti-oncogenic effects in mouse and human models of AF10 fusion AML. Collectively, our study identifies JAK1 as a tractable therapeutic target in AF10-rearranged leukemias.

**STATEMENT OF SIGNIFICANCE:** Gene fusions of AF10/MLLT10 are recurrent in acute myeloid and lymphoid leukemia and are associated with extremely poor survival outcomes. We show that the JAK1 kinase is required for activation of the AF10 fusion oncotranscriptome and for leukemogenesis. Since a number of JAK/STAT pathways inhibitors are in clinical development or approved for use, our studies may help develop a therapeutic strategy for AF10-rearranged leukemias.

## INTRODUCTION

A significant subsest of human malignancies display balanced chromosomal translocations or other genetic aberrations resulting in formation of chimeric fusion oncoproteins (1). Since some recurrent fusion proteins (FPs) found in cancer cells are key drivers of oncogenic signaling, they can be attractive candidates for therapeutic targeting. This is illustrated by the success of tyrosine kinase inhibitors (TKI) against BCR-ABL fusions in chronic myelogenous leukemia (CML) (2); ALK fusions in non-small cell lung carcinoma, T-cell anaplastic large-cell lymphoma and neuroblastoma (reviewed in (3)); TRK fusions in diverse malignancies (reviewed in (4)); and RET fusions (5). One reason the therapies cited above have been successful is because they target kinases—a class of proteins now pharmacologically tractable due to development of clinically-relevant small molecule inhibitors (6). Most other fusion oncoproteins, such as those involving transcription factors or chromatin regulators, have been difficult to target by traditional methods. One way of overcoming this hurdle is to identify molecular vulnerabilities arising from aberrant fusion oncoprotein expression in tumor cells. In this effort, defining tumor-promoting molecular pathways co-opted by fusion oncoproteins may help in the design of precise therapeutics against malignancies harboring those fusions.

The AF10/MLLT10 gene, located on chromosome 10, is recurrently involved in chromosomal translocations in T-cell and B-cell acute lymphoblastic leukemia (T-ALL and B-ALL) as well as acute myeloid leukemia (AML) (7-9). These translocations, which lead to the fusion of AF10 – a chromatin regulatory protein – to an N-terminal partner on another chromosome, are found in both children and adults (reviewed in (7-9)). AF10 fusions to the mixed lineage leukemia gene (MLL) comprise 17-33 % of all MLL-fusions in childhood AML, making it the second most common MLL-fusion event in pediatric AML. Apart from MLL, AF10 also recurrently fuses to the CALM/PICALM gene on chromosome 11. Notably, in pediatric T-ALL, CALM-AF10 fusions are the most frequent gene-fusion events (10). Novel translocations in which AF10 is fused to several other partner genes have been described in both AML as well as in T-ALL (11-14).

In AML, AF10 gene-fusions are associated with adverse outcomes (15-18). AML patients with MLL-AF10 fusions show increased early morbidity, including leukocytosis-related complications, extramedullary disease, and a high risk of relapse (19). A study of pediatric AML patients with MLL-AF10 fusions observed that these patients had a median event-free survival of only 8 months (20). In a large multi-center international study of pediatric AML patients, it was demonstrated MLL-AF10 fusions are independent predictors of unfavorable prognosis. Patients with MLL-AF10 fusions fare significantly worse than those with most other MLL-fusions and are therefore often treated using high-risk protocols (21). In pediatric T-ALL, CALM-AF10 fusions are associated with early relapse and poor survival outcomes (22, 23). Given that there are no clinically effective, precision-medicine based approaches available to treat these leukemias and that currently mandated therapies cause considerable, long-term toxicities, there is an urgent, clinically unmet need to identify specific, targeted therapies for leukemias bearing AF10 FPs. In this study, using a series of complementary approaches, we demonstrate the dependence of AF10 FPs on JAK/STAT-mediated inflammatory signaling for leukemogenesis and in doing so identify a clinically tractable pathway to treat these leukemias.

## RESULTS

### Inducible mouse models of AF10 fusion-driven AML

AF10 FPs are implicated in the activation of transcription of oncogenic networks in AF10 rearranged (AF10-R) leukemia (reviewed in (9)) although the underlying mechanisms driving leukemogenesis remain largely elusive. We generated tetracycline (Tet)- repressible (Tet-Off) mouse models of AML driven by the two most recurrent leukemia-associated AF10 FPs, namely CALM(PICALM)-AF10 and MLL(KMT2A)-AF10 (henceforth termed iCALM-AF10 and iMLL-AF10 AML respectively). In the absence of tetracycline, retroviral expression of iCALM-AF10 or iMLL-AF10 in lineage-depleted bone marrow (BM)-derived hematopoietic stem and progenitor cells (HSPCs) led to rapid serial replating capacity *in vitro* (see Methods and **Fig. 1A** for schematic). We reasoned that these Tet-Off models would be useful for discovery of potential AF10 fusion-regulated gene networks, since doxycycline (Dox) treatment completely abrogates AF10-fusion gene expression (**Fig. S1A**). Dox-treatment of iCALM-AF10- and iMLL-AF10-transformed BM cells led to a significant decrease in the percentage of blast-like colonies by day 7, based on colony forming unit (CFU) assays (**Fig. 1B** and data not shown). Injection of AF10 FP-transformed BM cells into sub-lethally irradiated syngenic mice gave rise to aggressive AML with a latency of 3-6 months (median of 16 weeks for iCALM-AF10 and 20 weeks for iMLL-AF10) in primary transplants and a latency of 2-3 weeks in secondary transplanted AML. Strikingly, despite potent proliferative and blast-colony forming capacity *in vitro* and leukemogenesis *in vivo*, AML cells derived from both primary and secondary AMLs retained their acute sensitivity to Dox-induced loss of AF10 FPs. This continued dependence on AF10 FP expression was most evident from the fact that administration of Dox-containing chow to terminally ill, secondary iCALM-AF10 AML mice rapidly reversed disease symptoms, significantly increasing disease latency (**Fig.1C**). We then performed RNA-seq analysis to investigate transcriptome-wide changes following Dox-induced abrogation of AF10FP (iCALM-AF10 or iMLL-AF10) expression. Similar experiments were also performed with the iMLL-AF9 fusion oncogene. Importantly, these RNAseq data on the three fusions can be used to distinguish targets of MLL-fusions (common to MLL-AF10 and MLL-AF9) from targets of AF10 fusions (common to CALM-AF10 and MLL-AF10) (see **Fig. 1D** and **1E**). Genes that are downregulated when the leukemia-associated FPs were switched off were termed candidate “target genes” (**Table S1**). Most of the common targets of the 3 leukemic FPs were developmental transcription factors that have previously been implicated in leukemogenesis including *Hoxa* cluster genes (*Hoxa5, Hoxa7, Hoxa9 and Hoxa10*), *Meis1* and *Six1*, as well as other genes not previously implicated in AML, including *Ssbp2, Cnksr3*, and *Nt5e* (**Fig. 1E, Fig. S1B** and **Table S1**).

**Fig.1.**
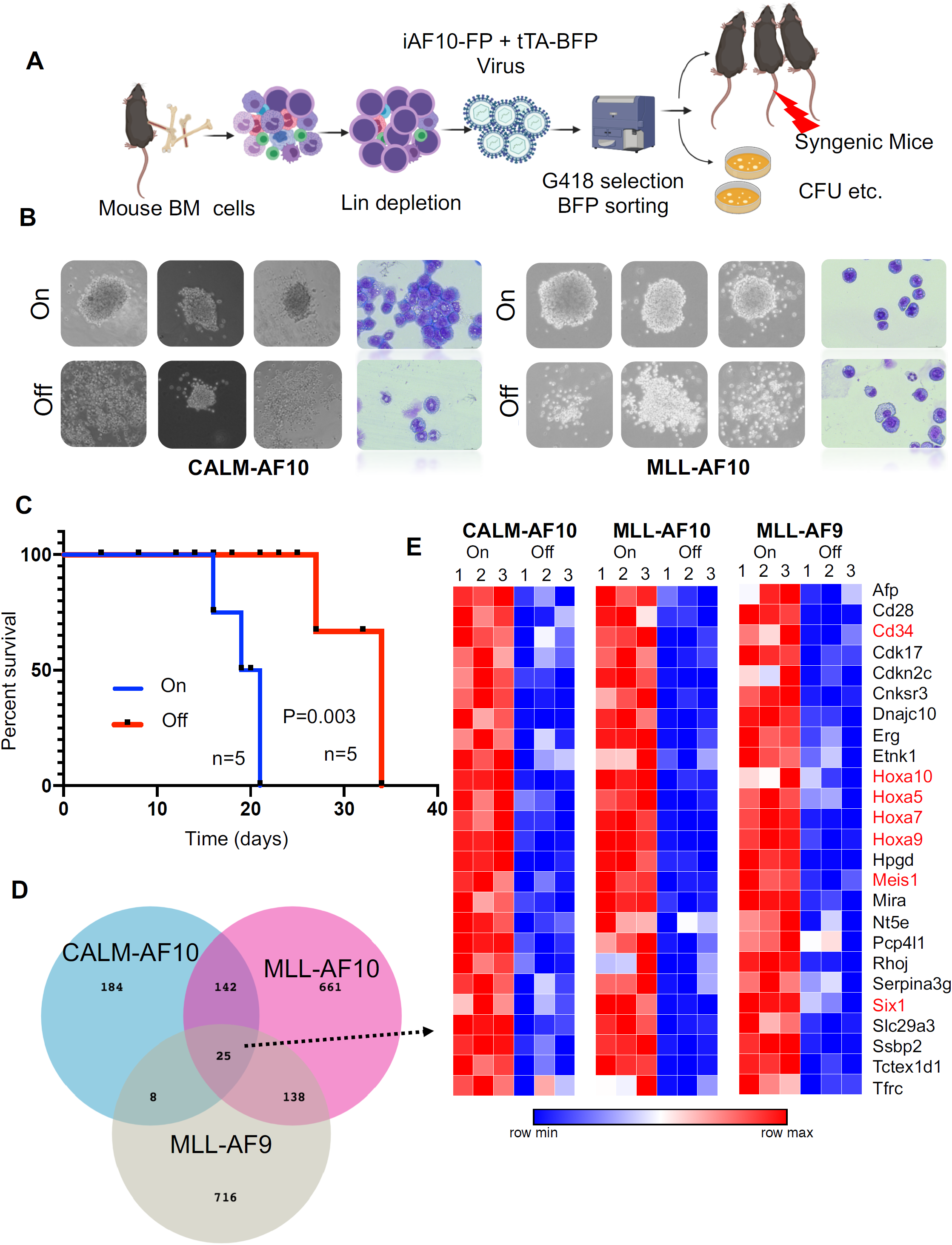
Inducible models of AF10 FP AML. **A)** Schematic representation of the bone marrow (BM) transplantation model (see Methods for detailed description). **B)** Representative images of colonies from iCALM-AF10 or iMLL-AF10 transformed BM cells with DMSO (AF10 fusion On) compared to doxycycline (Dox)-treated (AF10 fusion-Off) are shown. Also shown are Wright-Giemsa stained cytospins from the different indicated conditions. **C)** Kaplan-Meier survival curve for mice with the iCALM-AF10 fusion treated with vehicle (DMSO – CALM-AF10-On) compared to Dox (iCALM-AF10-Off) is shown. Number of mice in each group and P value of differences between two groups (Log rank Mantel-Cox test) is shown. **D)** Number of genes significantly downregulated (p value<0.05) when the respective fusion oncoproteins were switched-off using Dox within 48 hours are shown, with overlaps displayed in the Venn diagram. **E)** Heatmaps for the 25 genes downregulated by the all the 3 indicated AML-associated FPs are shown with gene names. Genes previously implicated in leukemia stem cell biology are marked in red.

### AF10 FPs drive potent activation of inflammatory signaling

An unbiased analysis of genes significantly associated with CALM-AF10 expression (**Table S1 –** CALM-AF10 target genes) revealed these genes to be strikingly enriched in signatures related to cytokine signaling, cytokine receptor activity, interferon signaling and innate immunity (**Fig. 2A** and **2B**). An Ingenuity Pathway Analysis (IPA) of CALM-AF10 target genes showed a highly significant enrichment for the JAK/STAT-mediated cytokine signaling pathway (P value: 1.48E-13) and revealed NFKB and STAT1/3 proteins as predicted upstream regulators of CALM-AF10-target genes (**Fig. S2A**). Consistent with these data, mapping of H3K27 acetylated super-enhancers (SEs) in CALM-AF10-On versus CALM-AF10-Off states (**Table S2**) demonstrated association of CALM-AF10 fusion expression with activation of a large number of SEs at genes associated with inflammatory signaling (-log10 p value= −12.6, **Fig. S2B**). Since we were most interested in AF10 FP target genes, we focused on the subset of genes commonly downregulated when CALM-AF10 or MLL-AF10 FPs (AF10 FP target genes **Table S1**), but not MLL-AF9, were turned off. These factors, which we term AF10 fusion target genes (**Table S1**), showed striking enrichment for inflammatory signaling pathways (**Fig. S2C**).

**Fig.2.**
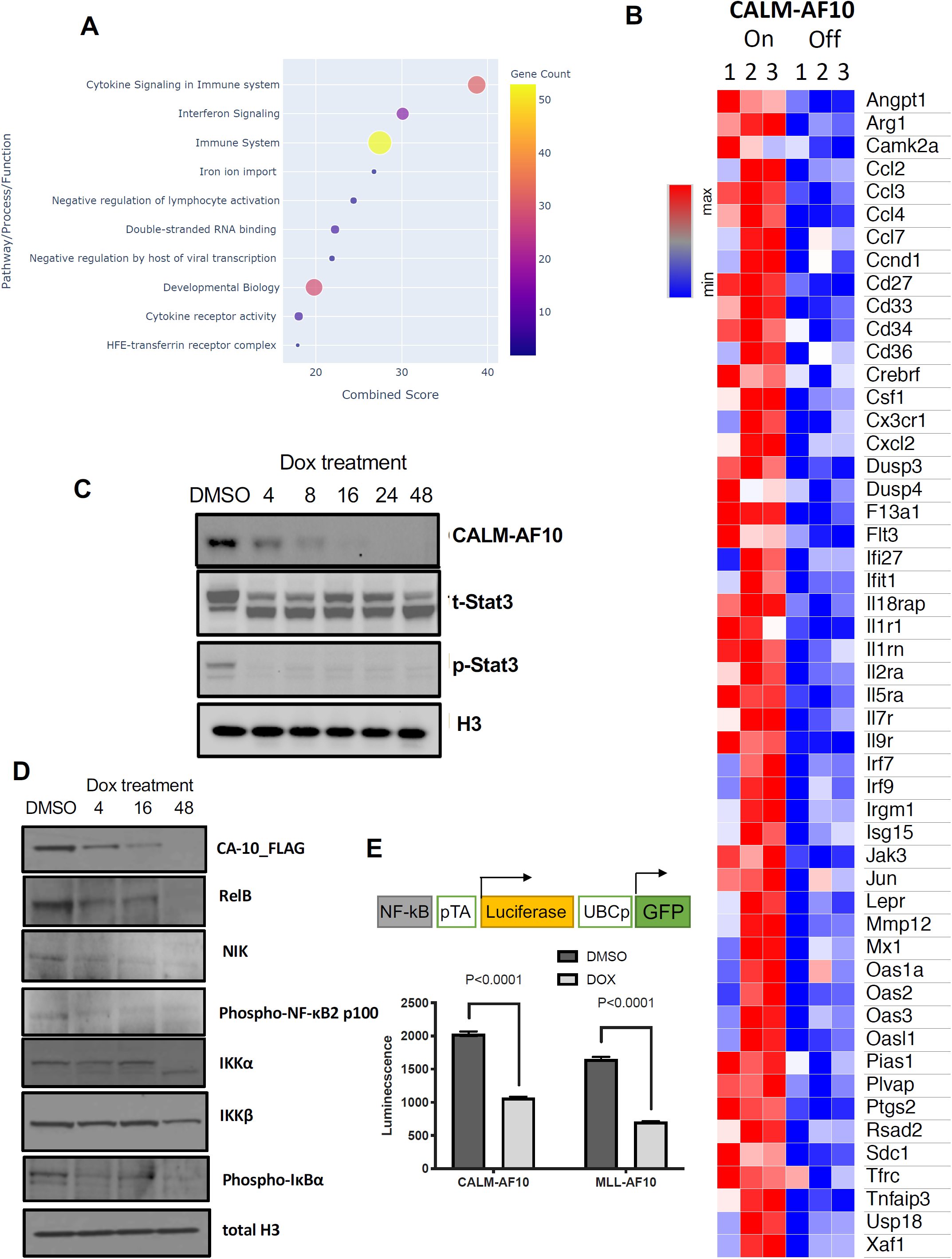
Activation of Inflammatory Signaling by AF10 FPs: **A)** Metascape pathway enrichment analysis of CALM-AF10 target genes is shown with the combined score plotted on the X-axis. Size and color of the cirlces are proportional to the gene counts. **B)** A representative heatmap showing genes involved in inflammatory signaling that are significantly elevated in the CALM-AF10-On compared to the CALM-AF10-Off state from RNA-seq data. **C)** Immunoblotting for Flag-CALM-AF10 (CA10), total STAT3 (t-STAT3) and phospho-STAT3 are shown at timepoints indicated above (in hours). Histone 3 (H3) is used as a loading control. **D)** Immunoblotting for NF-kB signaling pathway components at various indicated timepoints is shown with histone 3 (H3) as loading control. **E)** The relative luciferase activity (luminescence plotted as relative light units) of CALM-AF10-expressing (iCALM-AF10-On) compared to CALM-AF10-Off AML cells with an NF-kB luciferase reporter (schematic on top) is shown upon lipopolysaccharide (LPS) stimulation. Expression of the constitutive GFP reporter was used to normalize the signal.

To biochemically test the prediction that STAT1/3 and/or NF-kB are activated in CALM-AF10 AMLs (**Fig. S2A**), we monitored STAT1/3 phosphorylation and NFkB pathway activation using Western Blotting for key pathway components. We found that CALM-AF10 expression was strongly associated with phosphorylation of STAT3, a key downstream mediator of JAK/STAT signaling (**Fig. 2C**). This dramatic reduction in STAT3 phosphorylation, which occurred as early as 4 hours after CALM-AF10 inactivation, suggests that STAT3 activation is a direct or early/immediate consequence of CALM-AF10 protein expression in AML cells. Furthermore, CALM-AF10 activation was strongly associated with activation of the NF-kB signaling pathway, as assessed by immunoblotting for key effectors of the NF-kB pathway and confirmed using an NF-kB luciferase reporter assay (**Fig. 2D** and **2E**). Taken together, these studies indicate AF10 FPs—in particular the CALM-AF10 fusion protein—potently activate the transcriptional circuitry of inflammatory signaling networks.

### CALM-AF10 protein expression drives enhanced secretion of inflammatory cytokines

Our transcriptional data indicated that AF10-FP expression may lead to activation of several inflammatory cytokines and signaling molecules that in turn influence the proliferative activity or growth-factor sensitivity of AML cells. To test this possibility, we cultured murine BM cells transformed with CALM-AF10, MLL-AF10 or MLL-AF9 FPs in BM culture medium from which all supplemented cytokines were withdrawn (see Methods). Interestingly, a significantly higher proportion of murine BM cells transformed by either CALM-AF10 or MLL-AF10 remained viable in cytokine-depleted medium Vs. comparably treated MLL-AF9 transformed cells, which had close to 0% viability after 5 days of culture without cytokines (**Fig. 3A**). By contrast, co-culture of CALM-AF10 cells with MLL-AF9 cells in a contact-free trans-well culture significantly enhanced viability of MLL-AF9 transformed cells (**Fig. 3B**), indicating that cytokines and/or other growth factors produced and secreted by the CALM-AF10 AML cells may exert paracrine pro-survival effects on AML cells. To determine whether activation of inflammatory signaling associated with CALM-AF10 FP expression promotes secretion of inflammatory cytokines/chemokines, we quantitatively measured the relative abundance of cytokines, chemokines and growth factors in culture supernatants of CALM-AF10-On compared to CALM-AF10-Off AML cells using the Proteome Profiler cytokine antibody array (see Methods). Compared to CALM-AF10-Off cells, supernatants of CALM-AF10-expressing AML cells exhibited significantly greater abundance of cytokines implicated in AML cell growth and/or survival, including macrophage colony stimulating factor (M-CSF), granulocyte-macrophage colony stimulating factor (GM-CSF) and interleukin 3 (IL3) (**Figs. 3C** and **3D** and **Table S3**).

**Fig.3.**
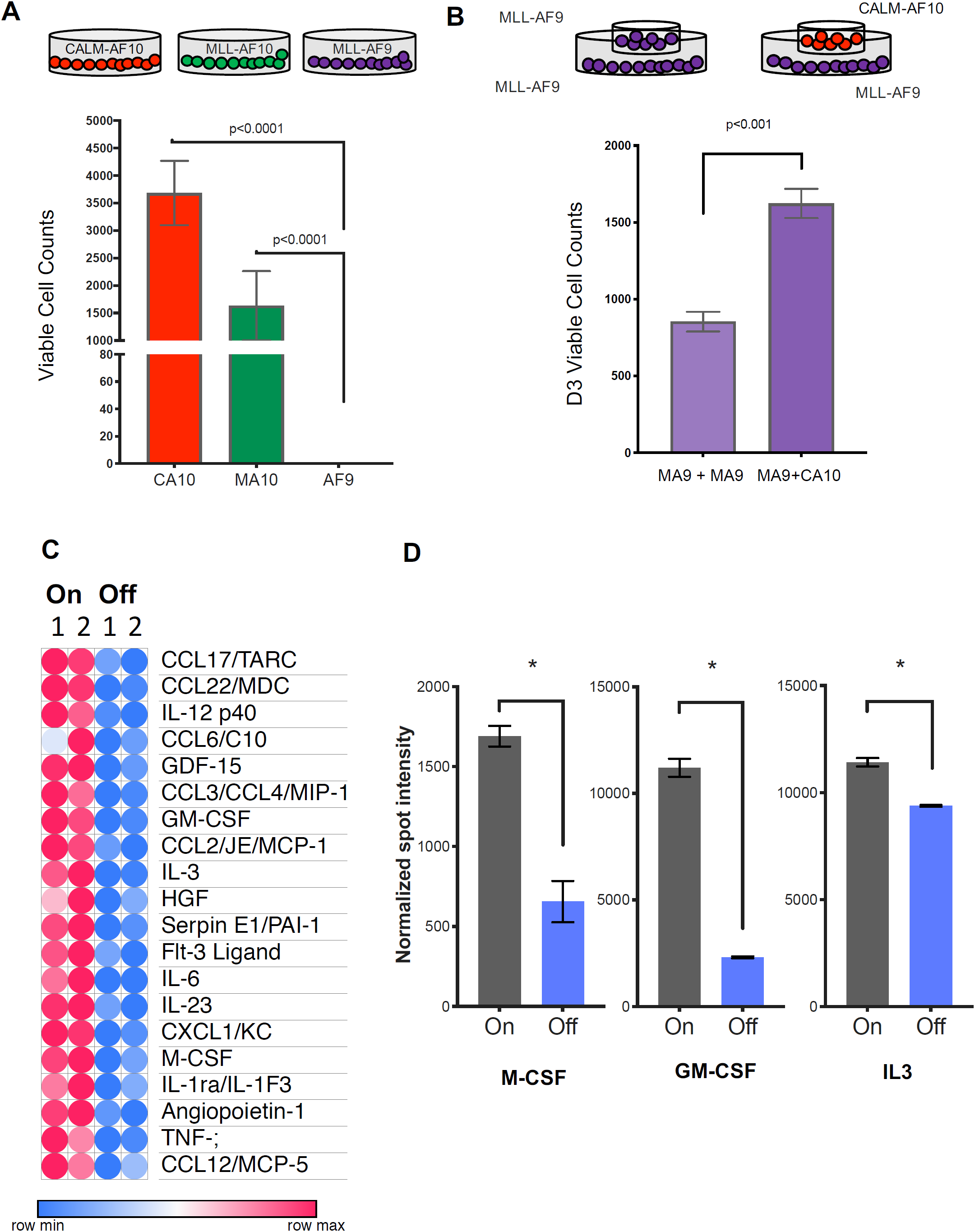
Cytokine sensitivity and expression in AF10 FP expressing AMLs: **A)** Number of viable cells per 10,000 initially seeded cells at days 5 after cytokine withdrawal are are plotted on the y-axis for the indicated AML fusion proteins. Legend: CA10: CALM-AF10, MA10: MLL-AF10, MA9: MLL-AF9. **B)** Schematic on top shows the format of the co-culture experiment in which a trans-well was inserted into a 6-well plate for co-culture. Number of viable cells per 10,000 cells at day 3 with MLL-AF9 AML cells co-cultured with the same cells on top (left panel) or overlaid with CALM-AF10 AML cells (right panel) are shown. **C)** Relative levels of cytokines, chemokines or growth factors measured as normalized spot-intesity in 2 replicates each of iCALM-AF10-On compared to iCALM-AF10-Off AML cells are shown. **D)** Heatmap shows relative levels of the top 20 most significantly altered cytokines/chemokines or growth factors in the iCALM-AF10-On and iCALM-AF10-Off conditions. **E)** Normalized spot intensity of indicated cytokines is shown in the bar graph with indicated P values.

### Interactome analysis reveals AF10 FP interaction with Jak1

In order to better understand how AF10 FPs may drive the expression of their target genes, we conducted a comprehensive and unbiased identification of the AF10 FPs interactome. We used affinity-purified epitope-tagged AF10 FPs and identified interacting proteins using Mass Spectrometry (MS) in murine AF10 fusion-driven AML cells (see **Fig. 4A** for schematic). We identified several novel CALM-AF10 interacting proteins including Stk38, serotransferrin (TF), Tmod3, Sun2 and thymidylate synthase (Tyms) (**Fig. 4B** and **Table S4**) and several of these interacting proteins (72 proteins) were also identified as MLL-AF10 fusion protein interactors (**Table S4**). Strikingly, the most abundant protein interactor of both the CALM-AF10 as well as the MLL-AF10 FPs was the Janus kinase protein Jak1. Of note, multiple fragments (30-32 unique peptides) of Jak1 were identified with high abundance in all 3 independent biological replicates of the CALM-AF10 dataset and none in IgG controls (**Fig. S3A** and **Table S4**). To confirm the AF10 FP-JAK1 interaction in AML cells, we used immunoprecipitation (IP) of CALM-AF10 or MLL-AF10 FPs using Flag antibodies (**Fig. 4C** and **Fig. S3B**). Reciprocal co-IP using antibodies targeting endogenous Jak1 further confirmed the interaction (**Fig. S3C**). We then asked how AF10 FP expression impacted Jak1 protein mechanistically. Expression of either CALM-AF10 or MLL-AF10 was associated with higher levels of Jak1 protein compared to AF10 FP-Off AML cells (**Fig. 4E**), although JAK1 mRNA levels were unchanged (**Fig. S3D**). This observation suggests that the presence of AF10 FPs stabilizes Jak1 and protects it from degradation. In support of this hypothesis, treatment with the proteasome inhibitor MG132 reversed the reduced Jak1 protein expression in AF10 FP-Off cells (**Fig. 4E**).

**Fig.4.**
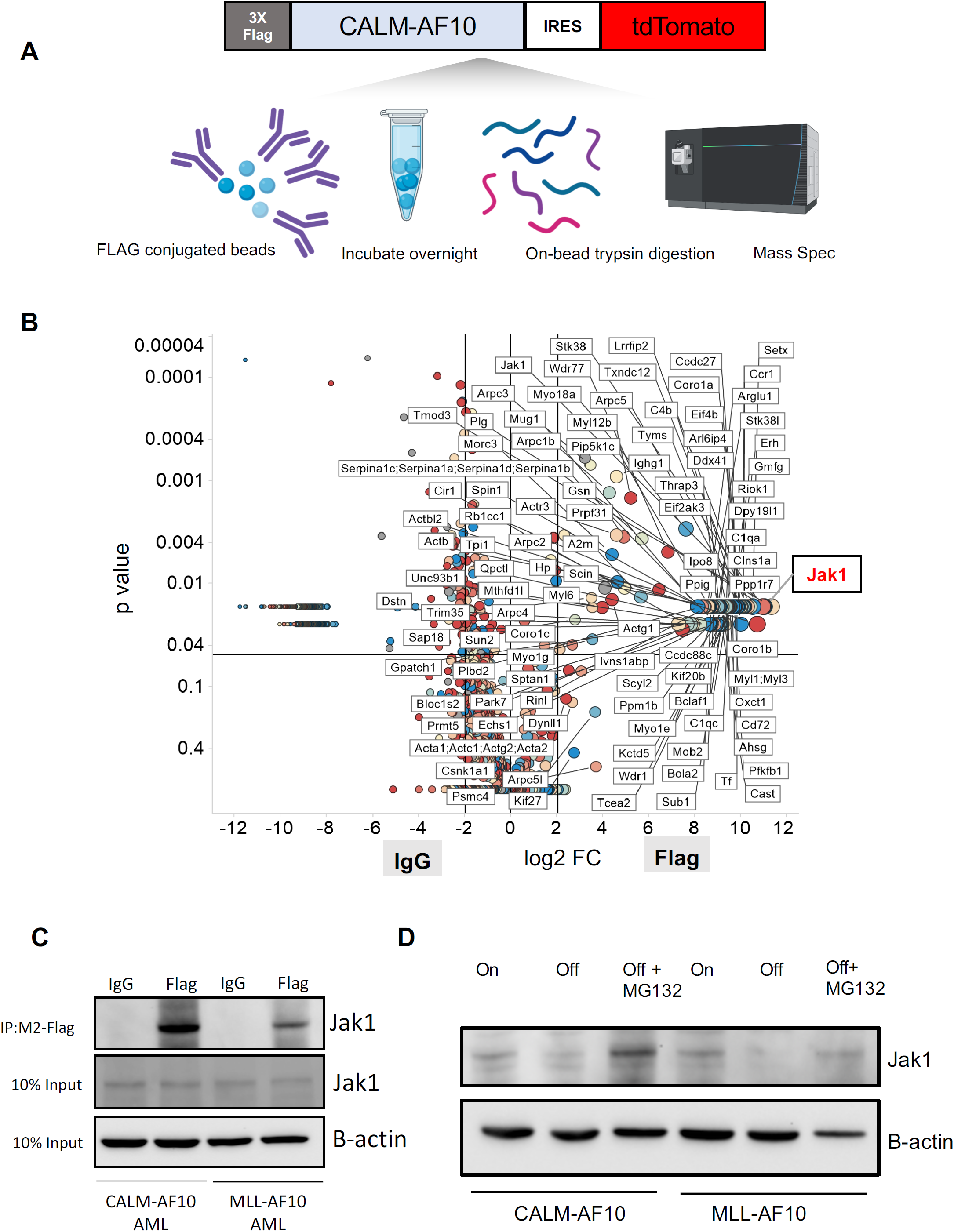
The interactome of CALM-AF10 fusion protein in AML A) **A)** Schematic representation of the CALM-AF10-flag pulldown and Mass Spectrometry (MS). **B)** Log2 fold-change (L2 FC – x-axis) and P-values (y-axis) of proteins differentially identified in the IgG control (left half) compared to 3xFlag-CALM-AF10 IP samples (right half) are plotted in the volcano plot. Murine CALM-AF10 AML cells were used for this study. **C)** IP using anti-Flag antibodies in CALM-AF10 or MLL-AF10 AML cells blotted with an antibody for endogenous Jak1. Beta-actin is used as a loading control. **D)** Immunoblotting for the endogenous Jak1 protein in murine iCALM-AF10 or iMLL-AF10 AML cells in the FP-On compared to FP-Off cells is shown together with Jak1 levels in MG132 treated AF10 FP-Off cells. Beta-actin is used as a loading control.

### CRISPR-screen of CALM-AF10 target genes identifies novel dependencies

Having identified several CALM-AF10 target genes and protein interactors, we sought to comprehensively determine whether these genes were required for the survival of CALM-AF10 AML cells *in vitro* and *in vivo*. For this analysis, we generated a pooled CRISPR library of sgRNAs against 420 CALM-AF10 transcriptional targets and protein interactors (**Table S5**) at a density of 10 sgRNAs/gene (See **Fig. 5A** for schematic and Methods). We then used this custom sgRNA library to conduct *in vitro* and *in vivo* CRISPR dropout screens in murine Cas9-expressing CALM-AF10 primary AML cells. Those analyses identified 28 statistically significantly depleted genes in the *in vitro* screen (**Fig. 5B**) and 19 in the *in vivo* screen (**Fig. 5C**), as evidenced by significant depletion of several independent sgRNAs relative to non-targeting controls (See Methods). Of note, 13 genes were significantly depleted *in vitro* and *in vivo* (**Fig. 5D**). Importantly, sgRNAs targeting *Jak1* were among the most significantly depleted guides *in vitro* and *in vivo* (**Fig. 5B** – **5D**), indicating that Jak1 is important for CALM-AF10 mediated leukemogenesis.

**Fig.5.**
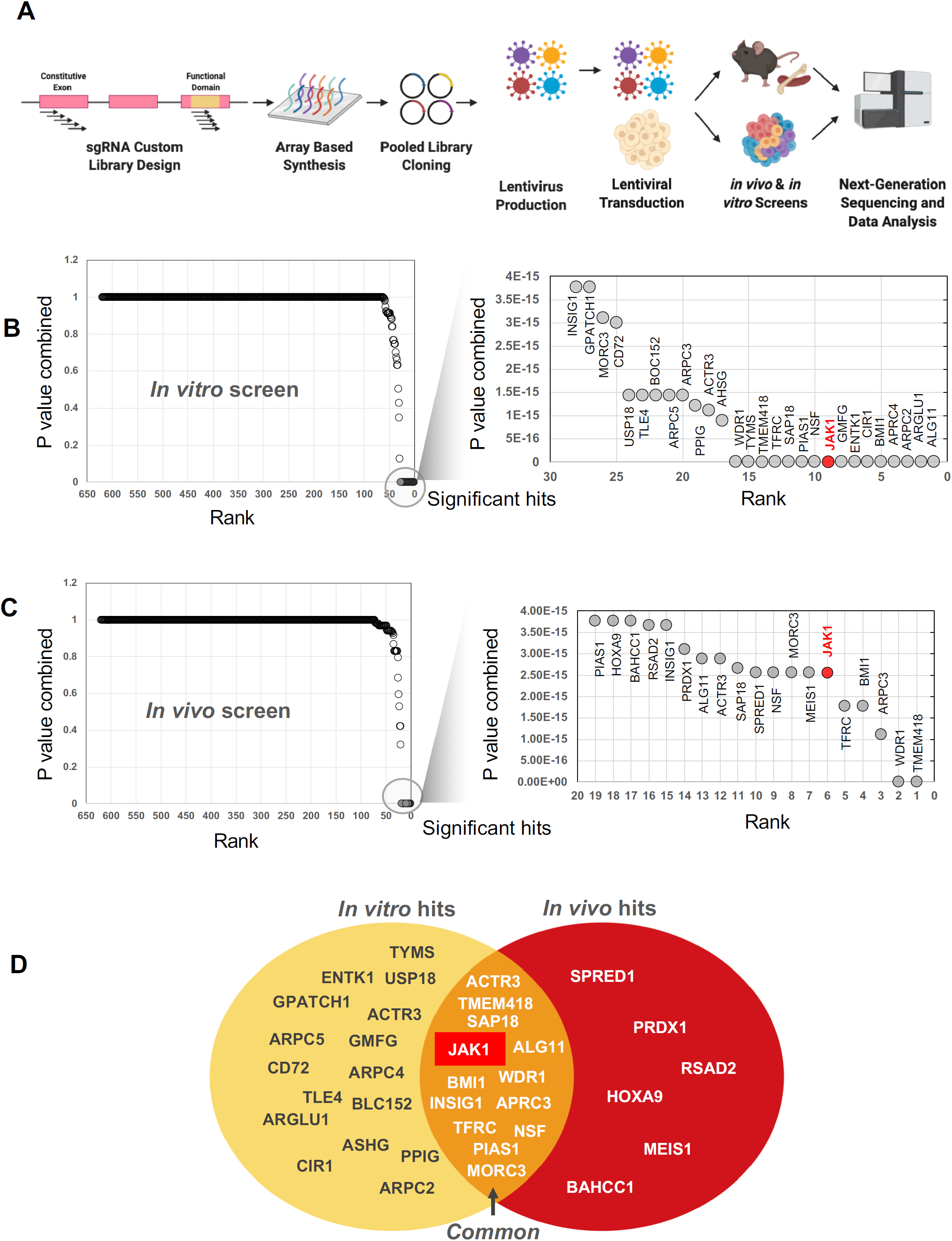
Genetic *Jak1* depletion inhibits CALM-AF10 mediated transformation: **A)**.Schematic of the CRISPR screen. A custom pooled library of sgRNAs targeting each of the 420 CALM-AF10 target genes and protein interactors was generated. 5 sgRNAs targeting the early constitutive exon and 5 targeting functional domains were combined for a total of 10 sgRNAs per gene. sgRNA library virus was transduced in CALM-AF10-Cas9 AML cells and sgRNA representation was determined after *in vitro* culture or from AML cells obtained from terminally ill mice. **B** and **C)** Plot showing the combined P values from 2 replicates (y-axis) of sgRNAs ranked in the order of depeletion (x-axis) in the T_0_ compared to the T_12_ (**5B**) or T_D_ (**5C**) timepoints. Statistically significant hits are zoomed in and displayed in a separate graph on the left of each figure. *JAK1*, ranked 9^th^ *in vitro* and 6^th^ in vivo is labelled in red. **D)** Venn diagram of *in vitro* hits (yellow oval) and in vivo hits (red oval) are shown with overlapping genes displayed in the union of the two sets.

### Genetic or pharmacologic Jak1 inhibition impairs CALM-AF10-driven transformation

Given that Jak1 was one of the top hits in our CRISPR library sceen, we addressed whether conditional *Jak1* deletion alters leukemic transformation by the CALM-AF10 oncoprotein. To do so, we transformed biallelic *Jak1* floxed BM cells with the CALM-AF10 fusion protein, introduced an estrogen-receptor-fused Cre (ER-Cre) recombinase into those cells, and then deleted *Jak1* floxed alleles by inducing with 4-hydroxytamoxifen (4-OHT) (**Fig. 6A)**. *Jak1* deletion led to a drastic reduction in the number of blast-like colonies in CFU assays and a concomitant, increase in the number of morphologically differentiated colonies relative to *Jak1* wildtype cells through 3 weeks of replating (**Fig. 6B** and **6C**). We also observed a significant increase in apoptosis in CALM-AF10 AML cells after Jak1 deletion (**Fig. S4A)**, while changes in cell cycling between genotypes were insignificant (data not shown). Furthermore, ectopic expression of dominant negative Stat3 (Stat3-DN) alone significantly decreased the number of second- and third-round blast-like colonies in CALM-AF10 AML cells, and together with *Jak1* deletion, completely abrogated colony formation in CALM-AF10 AML cells by 3 weeks of culture (**Fig. 6B**). This finding demonstrates that perturbing *Jak1* and/or its downstream effector *Stat3* impairs leukemic transformation of CALM-AF10 AML cells. Since we had comprehensively characterized CALM-AF10 target genes, we then asked how many of these genes were dependent on Jak1 for their expression. Strikingly, RNA-seq of *Jak1* deleted CALM-AF10 AML cells demonstrated that within 72 hours of 4-OHT-mediated *Jak1* deletion, a large proportion of the CALM-AF10 target genes were significantly downregulated as evidenced by gene-set enrichment analysis (GSEA) (**Fig. 6D**). CALM-AF10 target genes dependent on Jak1 included a high proportion of inflammatory signaling genes, including several inflammatory genes, cytokines and chemokines including *Ptgs2, Ccl3, Ccl4, Gdf15, Il2ra, Il1rn*, and *Csf1* **(Fig. 6E** and **Fig. S4B)** several of which were found in significantly higher abundance in the supernatant of CALM-AF10-On compared to Off leukemia cells **(Fig. 3C, Table S3** and **Fig. S6B)**. Taken together, these results demonstrated that Jak1 is critically required for the clonogenic capacity of CALM-AF10 transformed BM cells and for the transcriptional activation of its targets, including components of the inflammatory signaling cascade.

**Fig.6.**
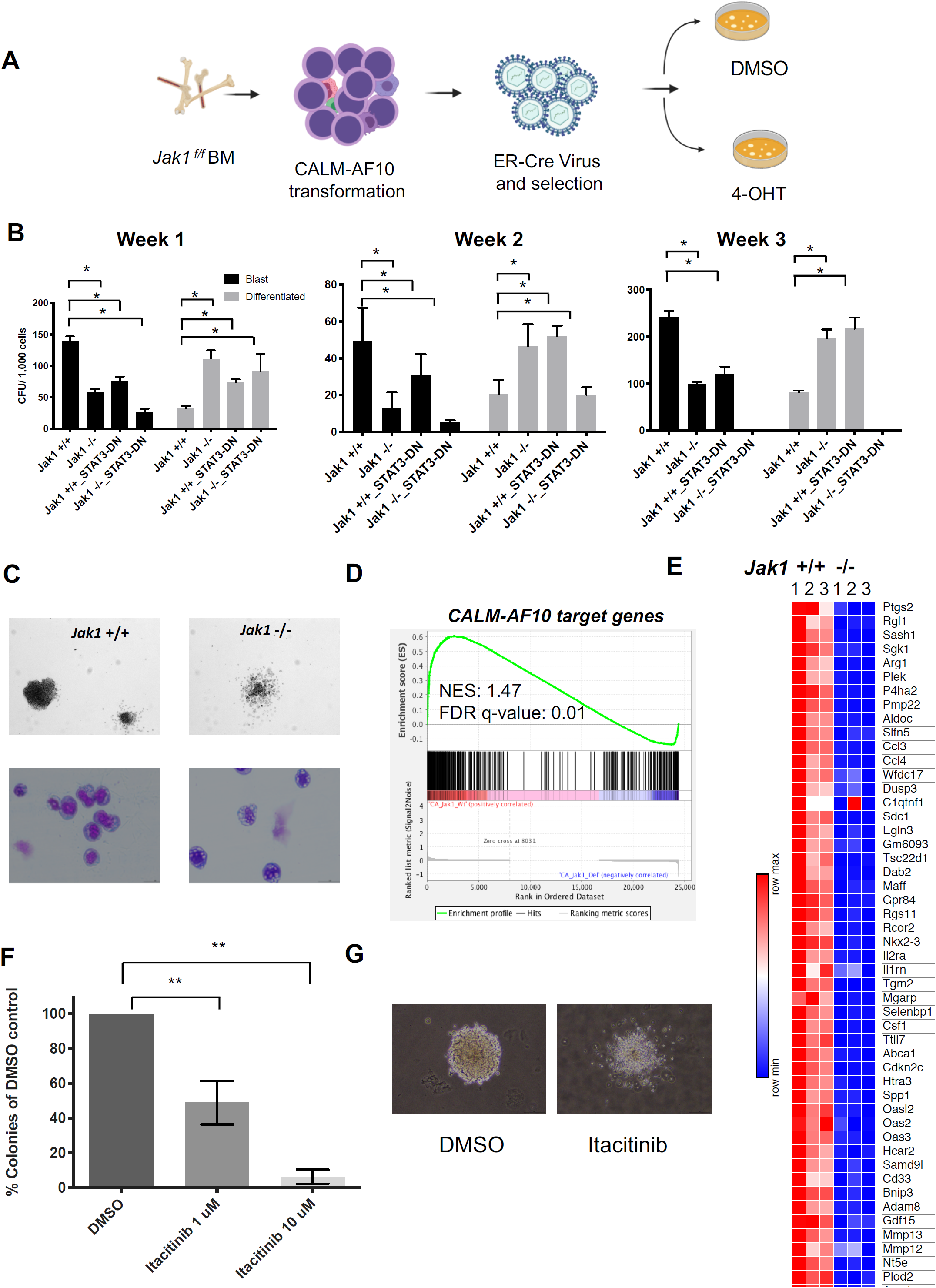
Jak1 inhibition impairs proliferation of AF10 FP AML cells: **A)** Schematic representation of the experiment – Bone marrow from *Jak1* floxed mice were used to transform lineage depleted cells with the CALM-AF10 fusion. Transformed cells were transduced with a Estrogen receptor-fused Cre recombinase (ER-Cre) and *Jak1* was deleted using 4-hydroxy-tamoxifen (4-OHT) treatment for CFU assays. **B)** Number of blast-like or differentiated colony forming units (CFUs) per 1,000 cells from CALM-AF10 transformed cells upon Jak1 deletion (or Stat3 dominant negative overexpression) is shown for weeks 1, 2 and 3. Legend: DN: dominant negative. **C)** A picture of a representative colony in *Jak1* wildtype compared to *Jak1* deleted CALM-AF10 transformed cells is shown (top) with a Wright-Giemsa stained cytospin of cells from the CFU assay (bottom). *= p<0.05, ** = P<0.01 **D)** GSEA showing the distribution of CALM-AF10 target genes in RNA-seq data from *Jak1* wild-type compared to *Jak1* deleted cells is shown with the Normalized Enrichment Score (NES) and FDR q-value indicated. **E)** A heatmap for the top 50 CALM-AF10 target genes with the highest GSEA rank metric score is plotted and shown for 3 *Jak1* +/+ and 3 *Jak1* -/- replicates. **F)** Total number of colonies from CALM-AF10 transformed cells treated with indicated concentrations of itacitinib are plotted as a percent of DMSO (vehicle)-treated cells. ** P<0.01. **G)** A representative picture of the most common type of colonies in DMSO compared to itacitinib-treated CALM-AF10 transformed cells is shown.

### Inhibitors of JAK/STAT signaling show potent anti-oncogenic activity in models of AF10 FP AML

Finally, we asked whether AF10 FP-positive AMLs respond to pharmacologic JAK1 inhibition by treating murine CALM-AF10 cells with the JAK1 inhibitor itacitinib. Similar to phenotypes seen in our gene-depletion studies, treatment of these cells with itacitinib significantly decreased their clonogenic capacity in a concentration-dependent manner (**Fig. 6F**) and almost completely eliminated blast-like CFUs, while the number of colonies exhibiting a differentiated morphology significantly increased (**Fig. 6G** and data not shown). These inhibitor effects were accompanied by significant induction of apoptosis (**Fig. S5A**). Next, we wanted to test whether atovaquone – a clinically approved anti-parasitic agent recently identified as a potent STAT3 inhibitor (25) – for activity against AF10-R leukemias. For this, we treated murine CALM-AF10 and MLL-AF10 AML cells with different concentrations of atovaquone. Our results showed that atovaquone treatment at clinically safe concentrations potently inhibited STAT3 phosphorylation (**Fig. 7A**) and significantly impaired the proliferation of both CALM-AF10 as well as MLL-AF10 AML cells (**Fig. 7B**) leading to a significant increase in apoptosis (**Fig. S5B**). Of note, AML cells derived from BM transformed with the CEBPA inhibitory protein Trib2 (26) were significantly less sensitive to atovaquone (**Fig. 7B**), demonstrating selectivity for AF10-R AML cells. Strikingly, atovaquone treatment almost completely abrogated the formation of blast colonies from murine CALM-AF10 and MLL-AF10 AML cells (**Fig. 7C** and **7D**). Consistent with murine AML cells, treatment with either itacitinib or atovaquone significantly reduced the clonogenic capacity of AML cells from a patient bearing the MLL-AF10 fusion in methylcellulose-based CFU assays supplemented with human cytokines (**Fig. 7E** and **7F**). Interestingly, treatment of normal human cord-blood derived CD34+ cells with itacitinib showed a striking reduction in their clonogenic capacity in the day 14 (d-14) CFU-assay. Atovaquone exposure on the other hand, had modest effects (**Fig. S5 C**). Finally, we treated non-obsese diabetic severe combined immunodeficiency (NOD/SCID) mice engrafted with the human CALM-AF10 positive U937 cell line with Mepron – a microparticle formulation of atovaquone used clinically in humans. Mepron administration in U937-engrafted mice led to a statistically significant increase in disease latency (**Fig. 7G**). Collectively, our data demonstrate that atovaquone treatment may show selectivity in terms of its anti-leukemic effects and also impair *in vitro* and *in vivo* leukemogenicity of AF10-R leukemia with modest effects on normal HSPCs – consistent with its demonstrated safety profile in human subjects.

**Fig.7.**
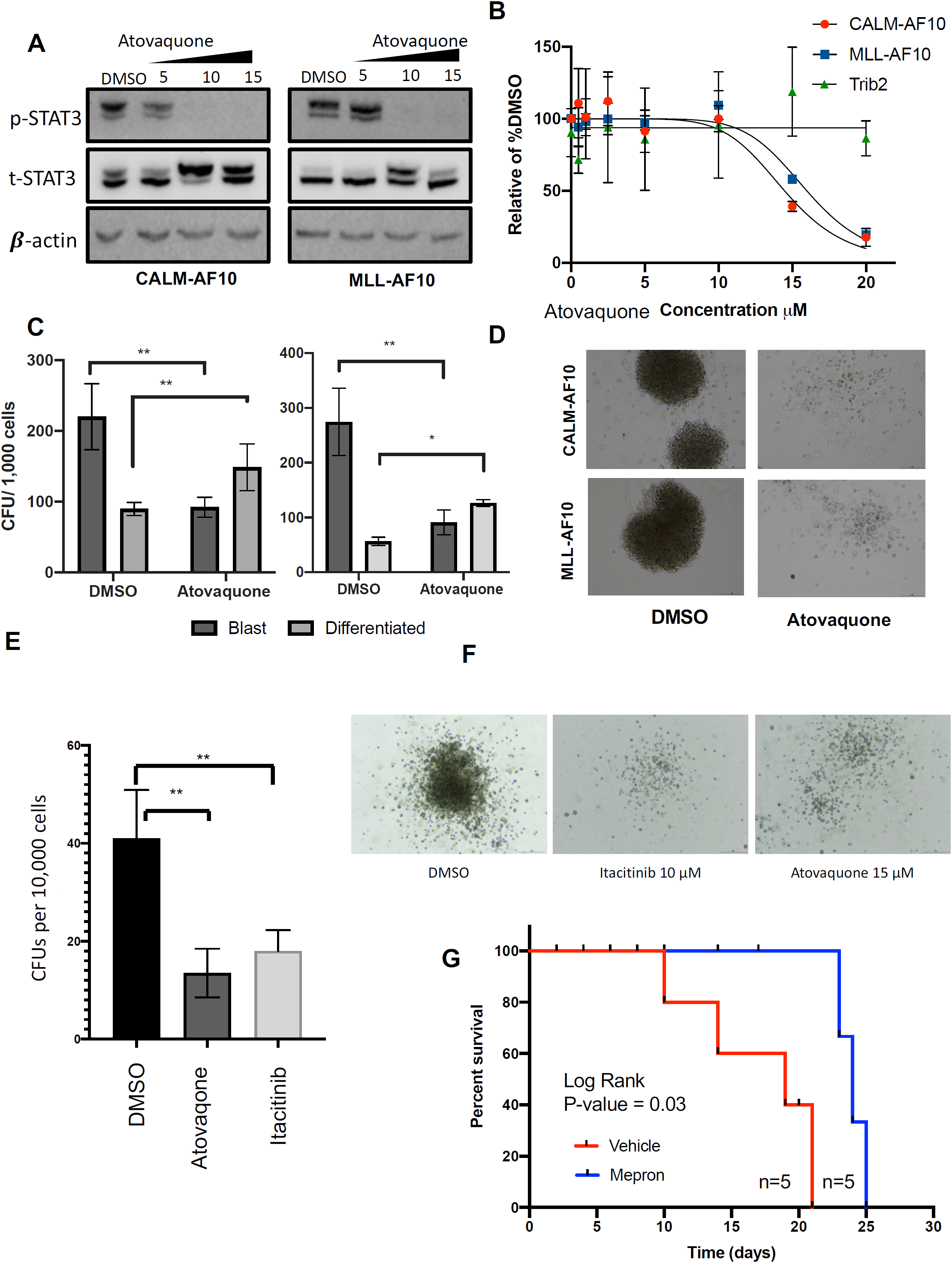
Atovaquone impairs AF10 FP-driven leukemogenesis *in vitro* and *in vivo*: **A)** Immunoblots showing phospho (p) or total (t) STAT3 with increasing indicated concetrations of atovaquone in micromolar concetrations are shown in CALM-AF10 or MLL-AF10 murine leukemia cells. B-actin is used as a loading control. **B)** Viable cells are plotted as a percentage of DMSO-treated cells (y-axis) with indicated concentrations of atovaquone (x-axis). Cells transformed with CALM-AF10, MLL-AF10 or Trib2 were used. **C)** Number of CFUs with a blast-like or differentiated morphology in DMSO or (10μM) atovaquone treated conditions is shown. *= P-value<0.05. **=P-value<0.01 **D)** Pictures of representative colonies from DMSO or atovaquone treated CALM-AF10 or MLL-AF10 AML cells are shown. **E)** Total number of day-14 CFUs per 10,000 cells from a human patient with MLL-AF10 fusion in methylcellulose supplemented with human cytokines are shown with DMSO, itacitinib or atovaquone treatment. **F)** Representative pitcture of a colony from patient cells treated with indicated drugs or DMSO in the d-14CFU assay are shown. **G)** Survival curves from mice injected with the CALM-AF10 positive P31 cell line administered vehicle (red) or Mepron – a clinically-used formulation of atovaquone, are shown. P-value calculated using Log-Rank T-test (Mantel-Cox) and the number of mice (n) in each group are indicated.

## DISCUSSION

AF10 gene fusions are generally associated with activation of overlapping molecular pathways and with poor survival outcomes (15, 22, 23, 27). A recent study of pediatric AML reported that AF10 fusions are significantly enriched in patients unresponsive to frontline therapies (the so-called “Induction Failures”) (28). These patients are therefore considered one of the highest risk-groups in pediatric AML, further reinforcing the urgent, unmet need to identify efficacious and highly-targeted therapies for this malignancy in the pediatric cohort.

While other targeted therapies are being explored as treatment for AF10-R AML, there is so far limited success in the clinic. It was previously demonstrated that the histone methyltransferase DOT1L is an attractive target in AF10-R AML, but the small molecule DOT1L inhibitor pinometostat has met with limited success against AML (29). Other notable actionable targets have been reported for CALM-AF10 AML, including the nuclear export protein XPO1 (30, 31), the Polycomb group protein BMI1 (32) and factors associated with transferrin-mediated iron uptake (33), but the translational potential of these targets remains to be determined in clinical studies. Patients with AF10 fusions continue to be treated with extremely cytotoxic therapies associated with induction failures and severe debilitating effects even when lasting cures are achieved.

Notably, studies performed in cell lines show that AML cells harboring AF10 fusions are highly sensitive to direct genetic targeting of the AF10 FP, evidence that these fusions are attractive candidates for targeted therapy for AF10-R leukemia. Specifically, siRNAs targeting the CALM-AF10 fusion transcript lead to potent loss of viability of CALM-AF10 AML cells, and sgRNAs targeting the fusion partners of CALM-AF10 are highly selected against in DepMap Screens (34, 35). Despite a strong rationale for targeting tumor-promoting fusion oncoproteins in cancer, most of these oncoproteins remain difficult drug targets. One way of circumventing this problem is by identifying downstream transcriptional or signaling networks activated by these FPs and targeting these networks instead. In this study we developed mouse models where the key AF10 fusion oncoproteins CALM-AF10 and MLL-AF10 can be rapidly switched off, enabling the systematic mapping of genetic and biochemical networks driven by AF10 FPs. In addition to these transcriptomic studies, our extensive characterization of the AF10 FP interactome helps expand the molecular network of AF10-R leukemia, which will help in understanding mechanisms of AF10-R leukemia that have so far remained elusive. More importantly, these datasets on AF10-R leukemia will help identify and prioritize therapeutically targetable candidates in the disease. Our finding that AF10 FPs directly recruit JAK1 and potently activate inflammatory signaling offer one such attractive avenue for therapeutic intervention.

The JAK/STAT pathway is an important node in regulating expression, secretion, and activity of cytokines, chemokines, and growth factors that play important roles in normal and malignant cells. Rapid strides have been made in the last few years in the development, characterization and clinical translation of inhibitors of the JAK family of kinases, given their roles in inflammatory diseases and cancer. The JAK1/2 inhibitor ruxolitinib was the first JAK inhibitor approved by the FDA for use in myeloid malignancies (36). Furthermore, more selective JAK1 inhibitors such as itacitinib and filgotinib are being tested in advanced stage clinical trials for inflammatory conditions (reviewed in (37)).

Excitingly, FDA-approved drugs used as antiparasitic agents have recently been shown to inhibit STAT3 activation in hematological malignancies and are therefore being repurposing for use in cancer (24). Genomic studies have revealed that the antiparasitic drug atovaquone inhibits STAT3 phosphorylation and has anti-leukemia effects (25). In fact, atovaquone is sometimes prescribed to leukemia patients, not for its recently recognized anti-leukemia effect, but to ward off parasitic infections following immunosuppressive treatment. Interestingly, the anti-leukemia effects of atovaquone were observed in human AML patients in retrospective analyses, although it remains unclear which leukemia patients might benefit most from these therapies (24, 25). As a result, a clinical trial is currently ongoing for atovaquone combined with chemotherapy in *de novo* AML. Since atovaquone has been used safely in humans for several decades, we asked whether these drugs are effective against AF10-R AML. Our studies showing potent effects of atovaquone in mouse and human AF10-R leukemia cells indicate a novel therapeutic approach in this disease in addition to currently used therapies.

## ACKNOWLEDGEMENTS

The authors gratefully acknowledge Dr. Brian James for valuable assistance with the project. We would like to thank Yoav Altman from the Flow Cytometry core for his excellent support with cell sorting and flow cytometry. We would like to thank Dr. Warren Pear for the MSCV-Trib2 plasmid. We also acknowledge the support of our funding sources: NIH R00 CA154880, NIH/NCI P30 CA030199, the Rally Foundation for Childhood Cancer Research and Luke Tatsu Johnson Foundation under Award Number 19YIN45, an Emerging Scientist Award from the Children’s Cancer Research Fund, and the V Foundation for Cancer Research (TVF) under Award Number DVP2019-015. We acknowledge the support of the Lady Tata foundation to A.D and from the MSKCC Support Grant NIH P30 CA008748, and grants to R.L.L. including National Cancer Institute R35 CA197594 and SCOR grants from the Leukemia and Lymphoma Society.

## CONFLICTS OF INTEREST

R.L.L. is on the supervisory board of QIAGEN and is a scientific advisor to Loxo (until Feb 2019), Syndax. Mission Bio, Imago, C4 Therapeutics, and Isoplexis. He receives research support from and consulted for Celgene and Roche and has consulted for Lilly, Jubilant, Janssen, Astellas, Morphosys, and Novartis. He has received honoraria from Roche, Lilly, and Amgen for invited lectures and from Celgene and Gilead for grant reviews.

## AUTHOR CONTRIBUTIONS

B.C., A.D., K.B., M.K., and A.R-C, conducted the experiments; X.L. provided bioinformatic support for the sequencing studies, N.Y., C.E., I.J., and T.H. provided key reagents and technical expertise. P.A., T.H., R.W.R, A.B., S.M., S.A.A, S.C., Z.R. D.F. and R.L. provided critical resources and helped analyze data and prepare a draft of the manuscript; and A.J.D conceptualized the studies, analyzed data and prepared the manuscript.

## Supplemental Figure Legends

**Fig. S1A: Loss of CALM-AF10 protein Expression in the iCALM-AF10 Leukemia Model:** Immunoblotting for the Flag-tagged CALM-AF10 fusion protein at various indicated time (in hours) after Dox-induced repression of CALM-AF10. Histone H3 is used as a loading control.

**Fig. S1B**: **Enrichment of Developmental Pathways in Common Target Genes:** Metascape pathway enrichment analysis of genes commonly downregulated (significance <0.05) in the iCALM-AF10, iMLL-AF10 and iMLL-AF9 AMLs upon switching off fusion gene expression at 48 hours is shown with the −Log10 p value plotted on the X-axis.

**Fig. S2A: Prediction of Putative Regulators of CALM-AF10 Target Genes:** Metascape analysis predicting the top upsteam regulators of CALM-AF10 fusion targets (Table S1) is shown with the −Log10 p value of each specified regulator plotted on the X-axis.

**Fig. S2B: Pathway enrichment in CALM-AF10-associated Super Enhancers:** Metascape analysis pathway enrichment analysis of super-enhancer-linked genes activated in the CALM-AF10-ON, but not the CALM-AF10-Off state (Table S2) is shown with the −Log10 p value plotted on the X-axis.

**Fig. S2C: Pathway Enrichment of AF10 Common Targets:** Metascape analysis pathway enrichment analysis of genes commonly downregulated (significance <0.05) in the iCALM-AF10 and iMLL-AF10 but not iMLL-AF9 AML cells – termed AF10 fusion-target genes (Table S1) is shown with the −Log10 p value of each specified regulator plotted on the X-axis.

**Fig. S3A: Relative abundance of Jak1-specific peptides in CALM-AF10 IP studies:** The relative log intensity of unique peptides mapping to Jak1 (y-axis) in the IgG control (left) compared to the Flag pulldown in 3 independent replicates is shown.

**Fig. S3B: Jak1 pulldown using AF10-fusion proteins:** An independent replicate experiment of Jak1 pulldown from Flag-IP of CALM-AF10 or MLL-AF10 fusions in murine leukemia cells is shown. M2-Flag beads and IgG controls are shown and 10% input control loaded at the bottom.

**Fig. S3C: Reciprocal IP of CALM-AF10:** Immunoprecipitation was performed with an endogenous Jak1 antibody in 293T cells transfected with the Flag-CALM-AF10 protein. Flag-CALM-AF10 immunoblotting in control IgG pr Jak1 antibody pulldowns is shown. Total H3 is used as a loading control.

**Fig. S3D: JAK1 transcript expression in CALM-AF10-On Vs Off cells:** The abundance of Jak1 mRNA (in normalized FPKM values) is plotted on the y-axis in CALM-AF10-On compared to CALM-AF10-Off leukemia cells. P values were calculated using Student’s T-test. N.S= not significant.

**Fig. S4A: Apoptosis induction upon Jak1 depletion in CALM-AF10 leukemia cells:** Percent apoptotic cells upon treatment of CALM-AF10 floxed mouse leukemia cells transduced with ER-Cre and treated with DMSO (Jak1 +/+) or with 4-OHT (Jak1 -/-) are shown as measured by Annexin V staining. P values were calculated using Student’s T-test. *< 0.01

**Fig. S4B: Jak1 depletion leads to transcriptional downregulation of CALM-AF10 regulated inflammatory signaling genes:** Upper panel shows the expression levels of several cytokines, chemokines and growth factors in the CALM-AF10-On (dark bars) compared to the CALM-AF10-Off leukemias. FPKM values are plotted on the y-axis. Transcription of the same genes in *Jak1* +/+ compared to *Jak1* -/- leukemia cells is shown in the bottom panel. P values were calculated using Student’s T-test. *P<0.05, **P<0.01. ***P<0.005

**Fig. S5A: Apoptosis in itacitinib-treated leukemia cells:** Percent apoptotic cells upon treatment of CALM-AF10 or MLL-AF10 to indicated concentrations of itacitinib are shown as measured by Annexin V staining. P values were calculated using Student’s T-test. *P<0.05, **P<0.01. ***P<0.005

**Fig. S5B: Apoptosis in atovaquone-treated leukemia cells:** Percent apoptotic cells upon treatment of CALM-AF10 or MLL-AF10 to indicated concentrations of atovaquone are shown as measured by Annexin V staining. P values were calculated using Student’s T-test. *P<0.05, **P<0.01. ***P<0.005

**Fig. S5C:** Cord blood-derived CD34+ve purified cells were treated with 10mM of atovaquone, DMSO or Itacinib and plated in methylcellulose medium supplemented with human cytokines. Colonies were scored after 14 days of culture. Average from 2 replicates are shown. Lower panel shows 3 representative colonies from each indicated treatment. Legend: CFU-GEMM-oligopotent colony forming units containing granulocyte, erythrocyte, monocyte, megakaryocyte cells. GM – colony containing granulocyte and/or macrophage cells, BFU-E: erythroid burst-forming unit colony.

## MATERIALS AND METHODS

### Cell culture

Human leukemia U937 and P31-Fujioka cells were cultured in RPMI-1640 medium supplemented with 2 mM L-glutamine and sodium pyruvate, 10% Fetal bovine serum and 50 U/ml Penicillin/Streptomycin (Thermo Fisher Scientific, Carlsbad, CA), and incubated in 5% CO_2_ at 37°C. Murine leukemia cells were cultured in DMEM medium supplemented with 2 mM L-glutamine, 15% FBS and 50U/ml Penicillin/Streptomycin, in the presence of following cytokines: 10 ng/ml murine Interleukin 6 (mIL-6), 6 ng/ml murine Interleukin 3 (mIL3) and 20 ng/ml murine stem cell factor (mSCF) (all from Peprotech, Rocky Hill, NJ), and incubated in 5% CO_2_ at 37°C. 293T cells were cultured in DMEM medium supplemented with 2 mM L-glutamine and sodium pyruvate, 10% Fetal bovine serum and 50 U/ml Penicillin/Streptomycin, and incubated in 5% CO_2_ at 37°C.

### Plasmids

The CALM-AF10 and MLL-AF10 fusions described previously were cloned into MSCV-based vectors to generate the 3XFlag-CALM-AF10-IRES-tdTomato, 3XFlag-MLL-AF10-IRES-tdTomato, 3XFlag-TRE-CALM-AF10-IRES-tdTomato (iCALM-AF10), or 3XFlag-TRE-MLL-AF10-IRES-tdTomato (iMLL-AF10) plasmids.

### Virus preparation

An ecotropic retroviral packaging plasmid was used with MSCV-based plasmids containing the gene of interest to make retrovirus. Packaging plasmids pMD2.G and psPAX2 were used with pLKO plasmids to make lentivirus carrying plenti-guide RNA vector and luciferase reporter plasmid. Briefly, co-transfection was performed with packaging plasmids in 293T cells by polyethylenimine (PEI)-based transfection and virus containing supernatant was harvested by filtering through a 0.45 μm filter after 48 and 72 hours. Virus was stored at −80 °C until use.

### Mouse bone marrow transformation and leukemia

Hematopoietic stem and progenitor cells (HSPCs) from wild-type or Jak1 floxed C57/Bl6 mice were isolated using EasySep mouse hematopoietic progenitor cell isolation kit (Stemcell technologies, Vancouver, Canada) which remove lineage marker-expressing cells with biotinylated antibodies directed against non-hematopoietic stem cells and non-progenitor cells (CD5, CD11b, CD19, CD45R/B220, Ly6G/C(Gr-1), TER119, 7-4). Isolated cells were cultured *in vitro* for 24 hours in DMEM medium supplemented with 2 mM L-glutamine, 15% FBS and 50 U/ml Penicillin/Streptomycin, in the presence of following cytokines: 10 ng/ml mIL-6, 6 ng/ml mIL3 and 20 ng/ml mSCF in 5% CO_2_ at 37°C. 24 hours later, these HSPCs cells were transduced with recombinant murine stem cell virus (MSCV) vectors containing 3XFlag-CALM-AF10-IRES-tdTomato (MIT-CALM-AF10), 3XFlag-MLL-AF10-IRES-tdTomato (MIT-MLL-AF10), 3XFlag-TRE-CALM-AF10-IRES-tdTomato (iCALM-AF10), or 3XFlag-TRE-MLL-AF10-IRES-tdTomato (iMLL-AF10) virus-containing medium. For CALM-AF10 or MLL-AF10 transfored BM cells, HSPCs were tranduced with virus containing supernatant from MIT-CALM-AF10 or MIT-MLL-AF10 plasmids and sorted for tdTomato positive cells after 5 days on a FACSAria II system (Becton Dickinson, La Jolla, CA). For generation of inducible leukemias, iCALM-AF10 or iMLL-AF10 cells were co-transduced with an MSCV plasmid containing the Tet-transactivator tTA linked to a 2a-BFP fusion. Cells co-transduced with iCALM-AF10 or iMLL-AF10 and MSCV-tTA-2a-BFP were sorted for BFP and selected for 2 weeks using G418. *In vitro* transformed cells were intravenously injected into sub-lethally irradiated mice (600 Gy) to generate leukemias.

### Genotyping for Jak1 floxed bone marrow cells

Genotyping for Jak1 conditional knockout was performed using a three-primer strategy as previously described [REF]. PCR was performed with the following primers:

JakL3 (forward): AGTGACAGGGACCTGTCCCTAGTC

Ex LoxP (forward): GGGGATCCTCTAGTGGCATAA

Exc R (reverse): CGCGTGGAAAACTGCTAGA

The *Jak1*^WT^ allele resulted in a 300 bp band and *Jak1*^fl^ allele in a 400 bp band.

### Immunoprecipitation of AF10 fusion complex and proteomic analysis

3XFlag-CALM-AF10 or 3XFlag-MLL-AF10 expressing cells were pelleted by centrifugation at 500 × g for 3 min at 4°C and washed with PBS twice. The cells were lysed in RIPA lysis buffer (0.5M Tris-HCl, pH 7.4, 1.5M NaCl, 2.5% deoxycholic acid, 10% NP-40, 10mM EDTA) containing protease inhibitor cocktail (Millipore Sigma, Billerica, MA) for 20 min on ice. Whole cell lysates were clarified by centrifugation at 13000 × g for 20 min at 4°C and stored at −80°C until further use. Whole cell lysate (200 µg/ml) in 5 ml RIPA buffer was mixed with 50µl of 50% slurry of anti-FLAG M2 agarose (Millipore Sigma, Billerica, MA) and protease inhibitor (Thermo Fisher, Carlsbad, CA) and incubated with gentle rocking overnight at 4°C. The anti-FLAG M2 beads were washed five times with 50 mM ammonium bicarbonate (pH7.4). The sample was suspended in 50 mM ammonium bicarbonate with a final concentration of 1–2 mg/mL. Then, the proteins were reduced by dithiothreitol (DTT), alkylated by iodoacetamide (IAM) and quenched by DTT with the same concentration as IAM. Proteolytic digestion was performed after diluting ten times the solution volume by 50 mM ammonium bicarbonate with trypsin (50:1, protein / trypsin, w/w). Protein samples were digested by 2 µg sequencing grade trypsin (Promega, Madison, WI) at 37°C with shaking overnight. The protein was desalted and recovered by C18 Ziptip (Thermo Scientific, Carlsbad, CA), dried out by Speed-Vac (Thermo Scientific, Carlsbad, CA) followed by liquid chromatography mass spectrometry analyses.

### Western blotting

Whole cells were lysed in cold RIPA buffer containing the protease and phosphatase inhibitor cocktail (Thermo Fisher, Carlsbad, CA), supernatant was collectedafter 12,000 rpm centrifugation and stored at −80°C until use. Protein concentrations from whole cell lysates were determined using a Bradford reagent (Thermo Fisher, Carlsbad, CA). Protein samples (50µg total protein) were resolved by SDS-PAGE using 4-12% Bis-Tris gel (Nupage, Invitrogen, Carlsbad, CA) and transferred to a nitrocellulose membrane (Bio-Rad, Hercules, CA). The membranes were blocked with 5% non-fat milk in Tris-buffered Saline containing 0.01% tween-20 (0.01% TBST). Immunoblotting for specific proteins was carried out using the following primary antibodies: Mouse monoclonal antibodies against anti-FLAG M2 (Millipore Sigma, Billerica, MA), Jak1 (CST-50996), and STAT3 (CST-9139). Rabbit polyclonal antibodies against phospho-STAT3 (CST-9145), Histone H3 (Abcam, ab18521), RelB (CST-4922), NF-kB-inducing kinase (NIK) (CST-4994), phospho-NF-κB2 p100 (CST-4810), IKKα (CST-61294), IKKβ (CST-8943), and Phospho-IκBα (CST-2697) (all from Cell Signaling Technology, Danvers, MA). The membranes were probed with species-specific donkey anti-rabbit or goat anti-mouse HRP-conjugated secondary antibody and detected with enhanced chemiluminescence (ECL) detection kit (Thermo Fisher, Carlsbad, CA).

### Colony forming assays

For CALM-AF10 or MLL-AF10 On/Off experiments, iCALM-AF10 or iMLL-AF10 primary AML cells were plated in MethoCult M3234 medium supplemented with 10 ng/ml mIL-6, 6 ng/ml mIL3 and 20 ng/ml mSCF (STEMCELL Technologies, Vancouver, Canada) at a concentration of 500 or 1,000 cells per ml. Colonies were scored after 7 days using the ZEISS Primovert inverted microscope (ZEISS, Germany). For mouse Jak1 depletion experiments, ER-Cre transduced *Jak1*^*f/f*^ CALM-AF10 and MLL-AF10 mouse hematopoietic progenitor cells were grown for 1.5 weeks to obtain a highly proliferative transformed culture. Colony forming cell (CFC) assays were performed by plating 10^3^ tdTomato positive cells/ml in the same conditions as described above. Cells were treated with either 500 nM 4-hydroxytamoxifen (Millipore Sigma, Billerica, MA) or DMSO. Every week for 3 weeks colonies were scored as above. Tightly clustered, hypercellular colonies were termed blast or blast-like and diffused colonies with dispered, larger and more granular cells indicative of M-type, GM-type or G-type colonies were termed differentiated. Colony assays with itacitinib or atovaquone with CALM-AF10 or MLL-AF10 AML cells were performed and scored similarly. Colony cells were then aggregated by washing off using PBS and pooled for replating for testing their secondary or tertiary replating potential at the same cell numbers.

For human AML cells, cells from a human AML patient confired as MLL-AF10 positive using diagnostic PCR were plated in Methocult (H4434) semi-solid medium supplemented with human cytokines. Total number of colonies were ennumerated after 14 days of culture after various drug treatments.

### Cytokine arrays

Tetracycline-inducible CALM-AF10 mouse hematopoietic progenitor cells were cultured in 15% FBS DMEM medium without IL-6 and mCSF and treated with either 4 µg/mL doxycycline (DOX) (Millipore Sigma, Billerica, MA) or DMSO for 3 Days. The relative level of cytokines within the culture supernatant was determined using the Proteome Profiler Mouse Cytokine Array (R&D System, Minneapolis), as per manufacturer’s protocol. Briefly, culture supernatants were incubated with nitrocellulose membranes which spotted with 40 different antibodies to mouse cytokines, chemokines, and acute phase proteins overnight at 4°C on a rocking platform shaker. The membranes were washed three times with wash buffer and incubated with 1:2000 diluted streptavidin HRP for 30 min at room temperature on a rocking platform shaker. The membranes were subjected to enhanced chemiluminescence (ECL)-based detection and cytokine signals were visualized using ChemiDoc touch imaging system (BioRad, California).

### Cytokine starvation and co-culture

The co-culture assay was performed in a 24-well transwell assay plate (0.4 µm pore size) (Corning, USA). BM cells transformed with CALM-AF10, MLL-AF10, or MLL-AF9 fusion proteins were cultured in DMEM medium containing 15% FBS without mIL-6, mIL3, and mCSF cytokines for 5 and 10 days. MLL-AF9 cells and CALM-AF10 cells were seeded at the basal and apical chambers of the transwell plate, respectively; at an initial seeding density of 10^3^ cell/ 1mL and maintained at 37 °C in a 5% CO2 incubator for 3 and 5 days. Live cells were analyzed by trypan blue exclusion assay and enumerated using a hemocytometer (Marienfeld, Germany).

### Construction of custom sgRNA library

The custom sgRNA library was designed based on the transcriptional targets and proteomic interactors (420 genes) of CALM-AF10, resulting from the RNA-Seq and proteomics-MS experiments (Table S5). Ten independent sgRNAs were designed for each gene using the Broad Institute Genetic Perturbation Platform (GPP) CRISPRko sgRNA design tool. All the sgRNAs were designed using principles reported previously (38) and the sgRNAs with the prediction of high off-target effect were excluded. Non-targeting negative control sgRNAs were included in the design and represented ∼5% of the total library size. Library sequences were synthesized on an array platform and then cloned into the lenti-guide Puro vector (Addgene: #52963, kind gift from Dr. Feng Zhang, MIT). The library plasmid was amplified using ElectroMAX™ Stbl4™ Competent Cells (Invitrogen, cat.11635018). The number of bacterial colonies was kept >500 times the number of sgRNA sequences in the library to maintain representation. To ensure the representation and identity of sgRNAs in the amplified pooled lentiviral plasmids, deep sequencing was performed on a HiSeq instrument (Illumina) and fastq files were analyzed using CRISPREssoCount (39).

### Pooled CRISPR Screening

Single-cell clones of Cas9-expressing CALM-AF10 cells were established by cloning in a Lenti-Cas9-2A-Blast vector (Addgene: #73310, kind gift from Dr. Jason Moffat), followed by blasticidin selection (10 μg/ml, 5 days), FACS-single sorting and outgrowth. Single-sorted clones were tested for Cas9 editing using an sgRNA targeting the Roasa26 locus followed by TIDE (40). Two single clones with high (>90%) Indel frequencies at the Rosa26 locus were selected for pooled screening. Pooled CRISPR library virus was produced using HEK293T cells transfected with PEI, pMD2.G and psPAX2 as previously described (41). Virus titer was measured by infection of cells with serially diluted virus. To ensure transduction of a single sgRNA per cell, multiplicity of infection (MOI) was set to 0.3 ∼ 0.4. Adequate representation of sgRNAs during the screen was ensured by keeping >1000x cells in culture relative to the library size. Cells ware harvested at initial (day 3 post-puromycin selection) and final (20 population doubling times after the initial) time points. Extraction of genomic DNA was performed using the Zymo Quick-DNA™ Miniprep Plus Kit (Zymo Research, Cat. D4068) according to manufacturer’s instructions. Two-step PCR-amplification and sequencing library preparation was conducted with TaKaRa Ex Taq™ Polymerase (TaKara, Cat. RR001) and custom primers to achieve adequate sequencing coverage in 1×75bp single-end reads, following published guidelines (39). Barcoded libraries were pooled and sequenced using Hiseq 500 (Illumina). The sequence data were de-multiplexed and analyzed using PinAPL-py (42).

### RNA-Seq

Murine in vitro transformed iCALM-AF10 cells were plated at a density of 2 million cells per 5 mL per well of a 6 well Non-Tissue culture treated plate. Doxycycline was added to a final concentration of 4ug/mL to the wells to turn off the CALM-AF10 fusion in cells. DMSO was added to control wells. Control and DOX treated cells were pelleted for RNA extraction after 48 hours. RNA was extracted using RNeasy Mini kit (Qiagen). RNA-seq Libraries were prepared using NEBNext Ultra II Directional RNA Library Prep Kit (New England Biolabs, Ipswich, MA) for Illumina as per the manufacturer’s protocol. 500 ng of total RNA was amplified in 10 cycles of PCR amplification. RNA-sequencing performed as single read 50 nucleotides in length on the Illumina NextSeq500 by the Genomics core at SBP Medical Discovery Institute, san Diego.

### ChIP-seq

*In vitro* transformed iCALM-AF10 cells were cultured with or without Doxycycline for 48 h before collecting the pellets for Chromatin immunoprecipitation. Immunoprecipitation was performed as described earlier (43). 2 million cells from each sample were fixed using 1% formaldehyde and chromatin was sheared using Diagenode Bioruptor for 15 min with 15 cycles (with each 30 sec on, 30 sec off cycle) setting at 4 °C. ChIP for H3K27Acetyl was performed using the Anti-Histone H3 (acetyl K27) antibody (Abcam, ab4729) specific to the H3K27Ac modification. Library preparation on eluted DNA was performed using NEBNext Ultra II DNA library prep kit for Illumina (E7645S and E7600S) as per the manufacturer’s protocol. Library prepped DNA was subjected to sequencing by NextSeq 500 (Illumina, La Jolla, CA) with a single read of 50 nucleotides in length at the Genomics core, SBP Medical Discovery Institute (La Jolla, CA).

### Animal studies

100 μL of 200,000 U937 human myeloid leukemia cells were mixed with 100 μL of cold matrigel (Corning, Oneonta, NY) and subcutaneously injected into the flanks of 8 week-old female NOD/SCID mice (The Jackson Laboratory, Bar Harbor, ME). After tumors reached a size of 200 mm^3^ mice were treated with vehicle (benzyl alcohol, poloxamer 188, purified water, flavor, xanthan gum, and saccharin sodium) or Mepron by oral gavage (200 mg/kg/day) for 21 days. Mice were sacrificed when the tumor volume reached 2000 mm^3^.

### NFκB luciferase reporter assay

To evaluate the NF-κB activity in the presence or absence of AF10 FPs iCALM-AF10 or iMLL-AF10 AML cells were transduced with lentivirus expressing NFkB-TA-Luc-UBC-GFP-W (Addgene plasmid 49343, kind gift of Dr. Darrell Kotton). GFP positive cells were selected by FACS sorting and cultured for 2 days in the presence of 4 μg/mL doxycycline. Subsequently, cells were treated with 100 μg lipopolysaccharides (LPS) for one hour and washed three times with PBS. Cellular luciferase expression was quantified using the DualGlo luciferase assay system (Promega, Madison, WI), and GFP was measured using flow-cytometry.

### Quantitation of mRNA

The mRNA levels were measured by real time quantitative PCR (RT-qPCR) using standard protocols. Briefly, RNA was extracted using the RNA extraction kit (Qiagen Biotech, Hilden, Germany). Quality of the extracted RNA was verified at A260/A280 (ratio>1.8) in a spectrometer (Nanodrop One, Thermo Fisher, Carlsbad, CA). cDNA was synthesized from RNA using qScript cDNA SuperMix (Quantsbio, Gaithersburg, MD). For PCR-based amplification, cDNA and primers were added to the SYBR Green PCR Master mix or by using Taq Man probe (Thermo Fisher, Carlsbad, CA). The primer oligo sequence (supplemental table) and relative quantity of mRNA was determined by the ΔΔCT method as previously described (43) using Glyceraldehyde 3-phosphate dehydrogenase (GAPDH) as the internal reference.

