## Supplemental Figures for "A JAK/STAT-Mediated Inflammatory Signaling Cascade Drives Oncogenesis In AF10-Rearranged AML"

**S1A**

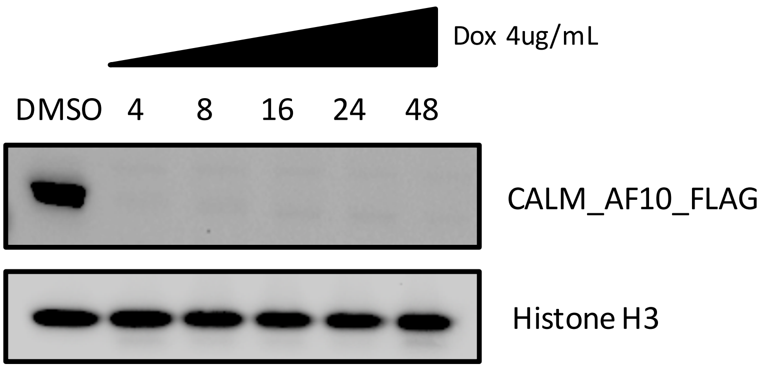


**Targets common to MLL-AF9, MLL-AF10 and CALM-AF10**

**S1B**

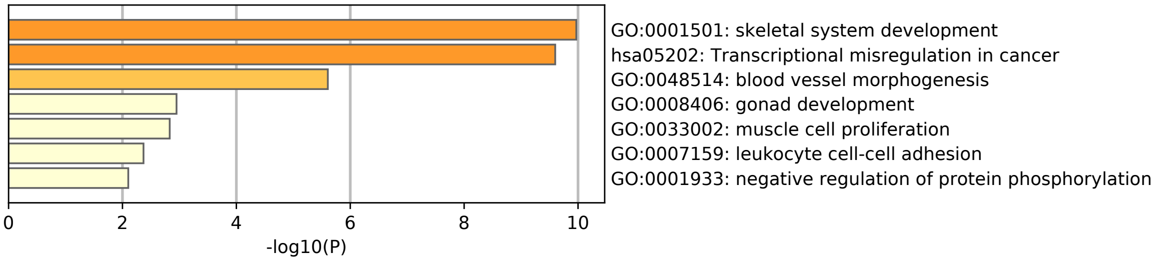


**Predicted Upstream Regulators of CALM-AF10 target genes**

**S2A**

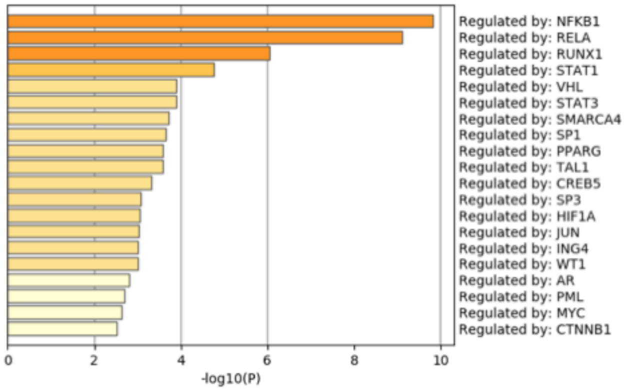


**S2B**

**Super-enhancer-linked genes in the CALM-AF10-ON leukemia**

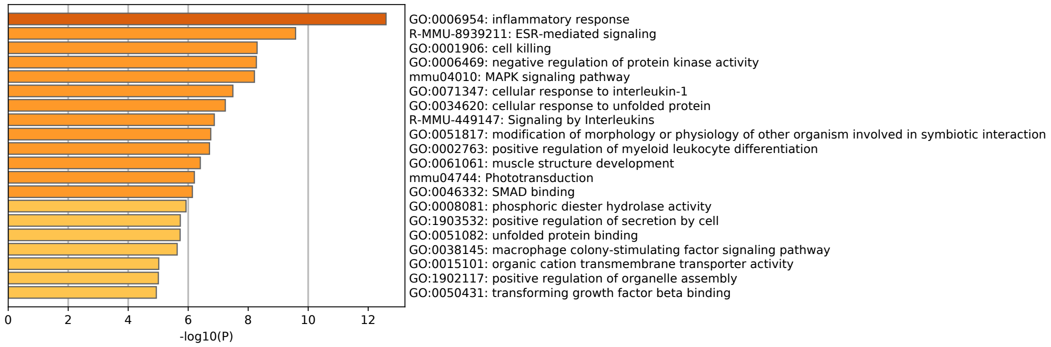

## S2C

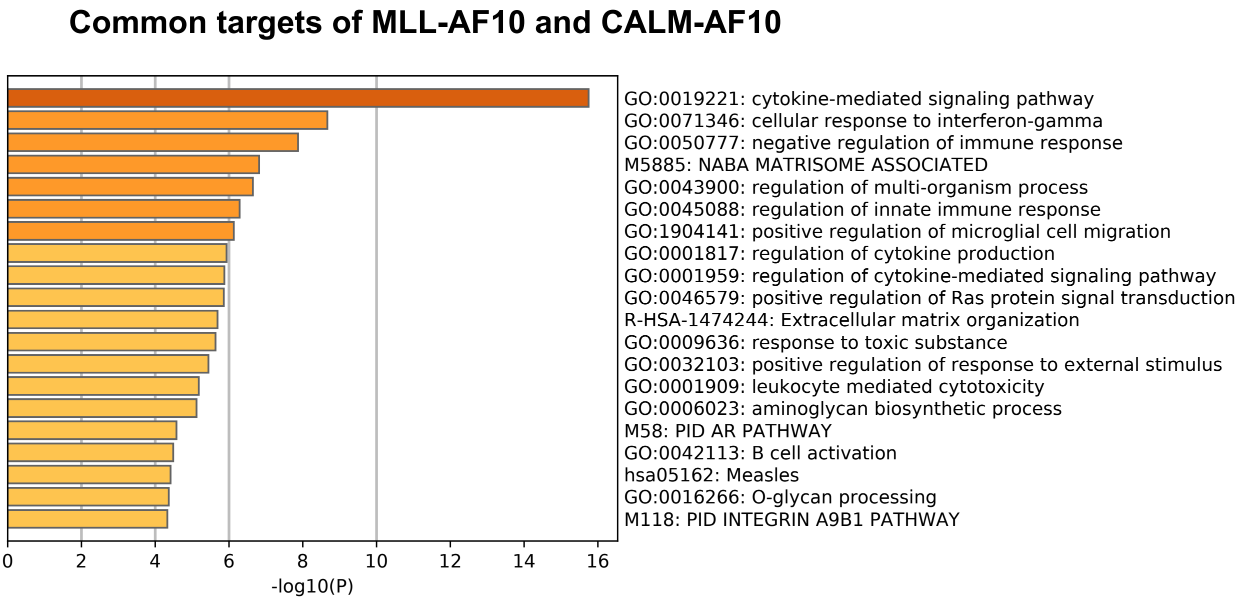


## S3A

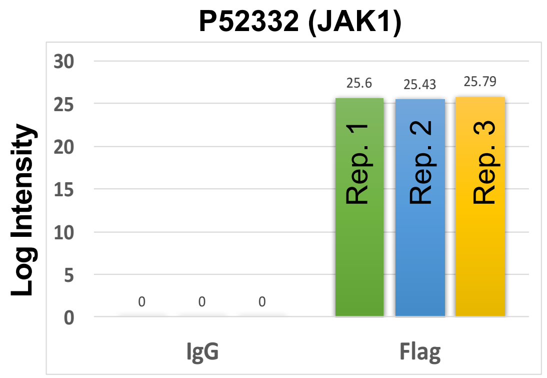


## S3B

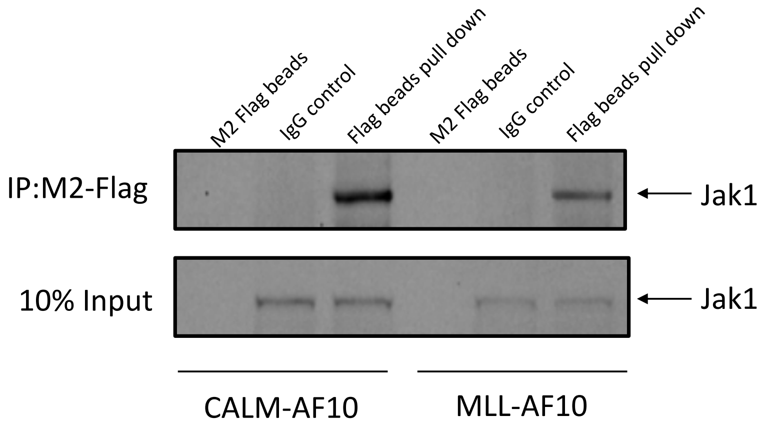


## S3C

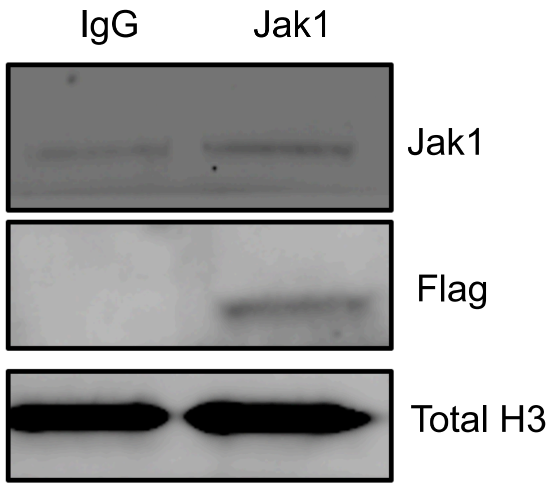


## S3D

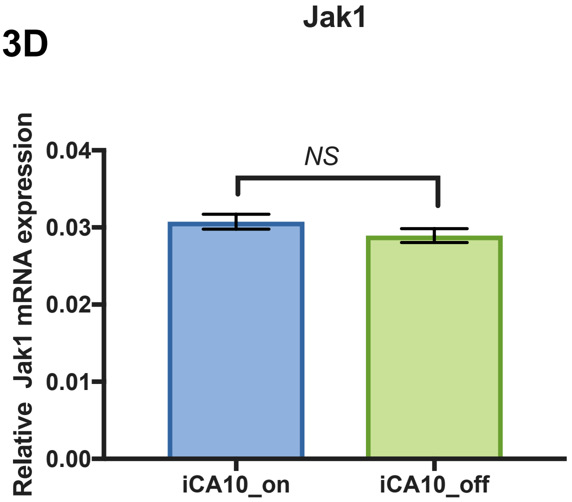

**S4A**

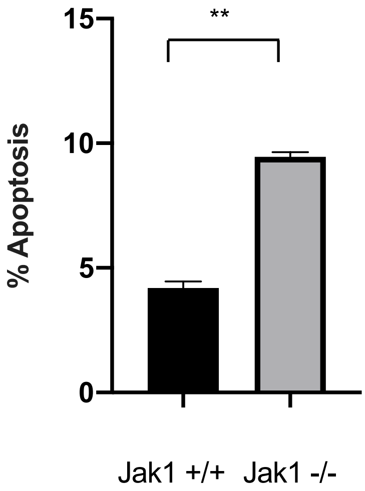

**
15
10
5
0
% Apoptosis
Jak1 +/+
Jak1 -/-


**S4B**

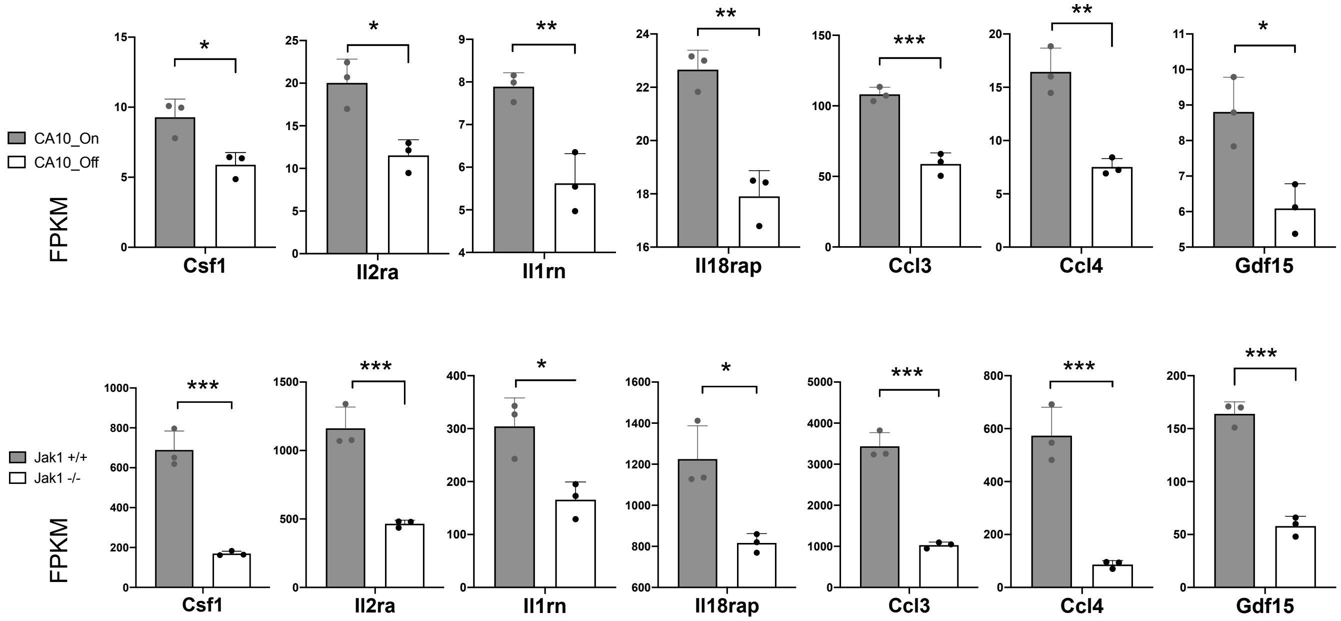

CA10_On
CA10_Off
FPKM
Csf1
Il2ra
Il1rn
Il18rap
Ccl3
Ccl4
Gdf15
Jak1 +/+
Jak1 -/-
FPKM
Csf1
Il2ra
Il1rn
Il18rap
Ccl3
Ccl4
Gdf15

**S5A**

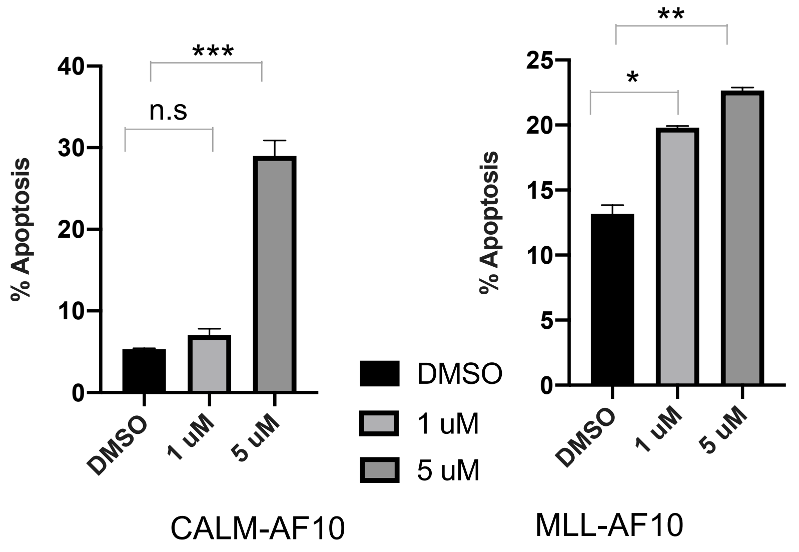
% Apoptosis
40
30
20
10
0
n.s
***
DMSO
1 uM
5 uM
CALM-AF10
DMSO
1 uM
5 uM
% Apoptosis
25
20
15
10
5
0
*
**
DMSO
1 uM
5 uM
MLL-AF10

**S5B**

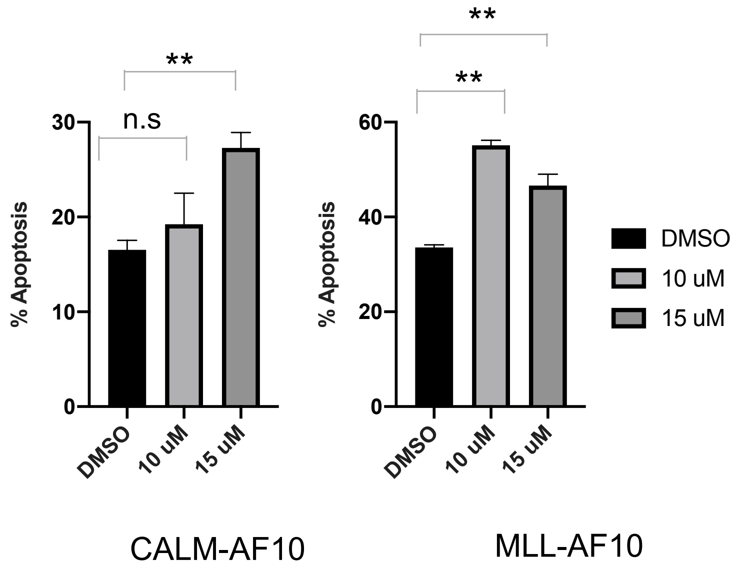
% Apoptosis
30
20
10
0
n.s
**
DMSO
10 uM
15 uM
CALM-AF10
% Apoptosis
60
40
20
0
**
**
DMSO
10 uM
15 uM
DMSO
10 uM
15 uM
MLL-AF10

S5C

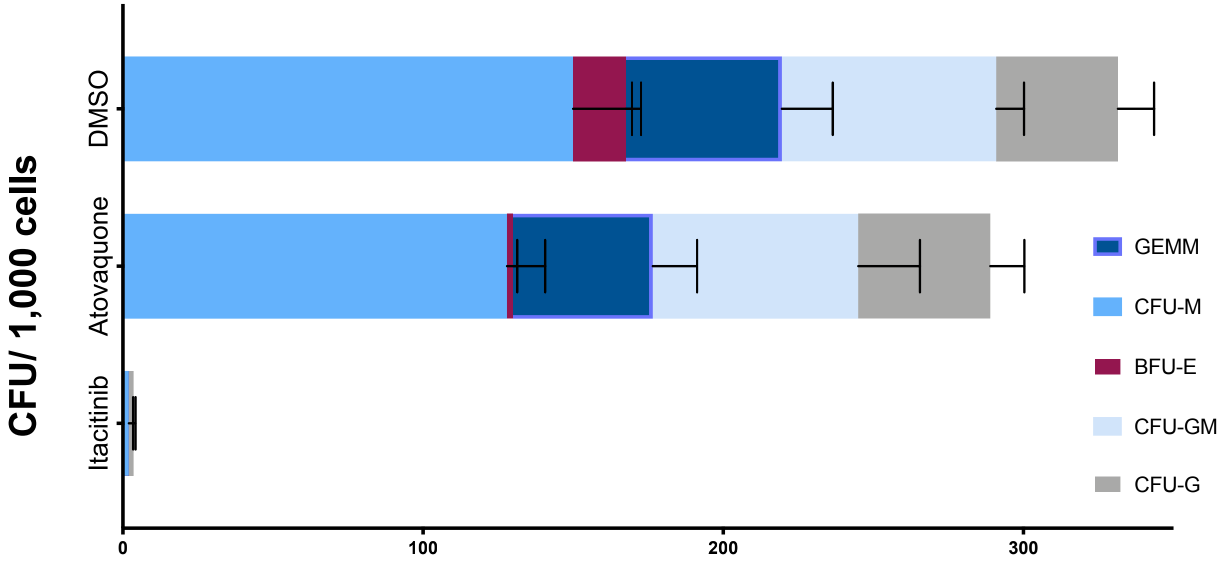


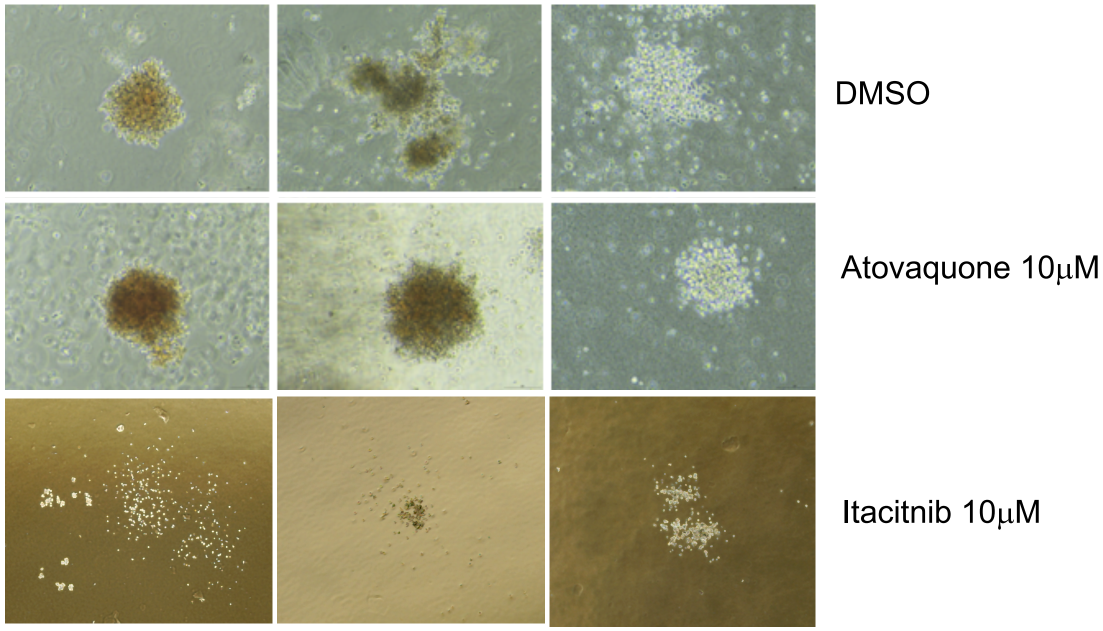


Cord blood CD34+ ve cells
