## Supplemental Tables for "A JAK/STAT-Mediated Inflammatory Signaling Cascade Drives Oncogenesis In AF10-Rearranged AML"

**TABLE S1**

| Down in CA10_Off (CALM-AF10 targets) | Down in MA10_Off (MLL-AF10-targets) | Down in MA9_Off (MLL-AF9-targets) | Common to _all 3 fusions (Common targets) | Common to AF10_fusions_only (AF10 fusion targets) |
|---|---|---|---|---|
| Il9r | Fxyd1 | Hpgd | Hpgd | Kcnh2 |
| Pmp22 | Pacsin1 | Cmklr1 | Cnksr3 | Rtp4 |
| Kcnh2 | Klhl30 | Meis1 | Slc29a3 | Arg1 |
| Oas2 | Skida1 | Ssbp2 | Hoxa10 | P4ha2 |
| Hpgd | Kcnh2 | Mira | Hoxa9 | Mmp12 |
| Oasl1 | Tpm2 | Hoxa5 | Mira | Ipcef1 |
| Rtp4 | Tmem178 | Sgip1 | Pcp4l1 | Mecom |
| Arg1 | Epha7 | Cnksr3 | Hoxa7 | Emp1 |
| P4ha2 | Siglech | Hoxa7 | Hoxa5 | Plod2 |
| Cdh2 | Afm | Hoxa9 | Nt5e | Il7r |
| Mmp13 | Hrc | Mef2c | Meis1 | Nkx2-3 |
| Apln | Gimap6 | Gstm5 | Ssbp2 | Tnip3 |
| Mmp12 | Flt3 | Car4 | Cdkn2c | Ccl4 |
| Ipcef1 | Slitrk2 | Znhit3 | Cd28 | Oas3 |
| Ifit1 | Col19a1 | Cdk6 | Six1 | Irf7 |
| Mecom | Msrb3 | Cdk17 | Etnk1 | Oasl2 |
| Ptgs2 | Gimap1 | Slc29a3 | Tfrc | Eya1 |
| Cnksr3 | Sema6d | Trim59 | Erg | Atp6v0d2 |
| Slc29a3 | Il31ra | Kmt2a | Rhoj | Spp1 |
| Adgrd1 | Pdzd4 | Pbx3 | Serpina3g | Ccnd1 |
| Scml4 | Mecom | Akip1 | Afp | Piwil2 |
| Hoxa2 | Ccr7 | Fnbp1l | Cdk17 | Scn3a |
| Emp1 | Paqr5 | Parp8 | Dnajc10 | Gcnt4 |
| Plod2 | Mrvi1 | Pus7l | Tctex1d1 | Ccl3 |
| Hoxa10 | Dpp4 | Gng12 | Cd34 | Adgrl4 |
| Hcar2 | Wnt10b | Tmem119 | | Gbp6 |
| Slc26a9 | Spib | Nrg4 | | Gm5111 |
| Pla2g4c | P4ha2 | Baz2b | | Xaf1 |
| Il7r | Jag2 | Lhfp | | Pim2 |
| Nkx2-3 | Plod2 | Mylk | | Il2ra |
| Hlf | Dlk1 | Runx2 | | Cd27 |
| Tnip3 | Rcor2 | Klhl5 | | Lhfpl2 |
| Ccl4 | Epb4.1l4b | Pdcd4 | | Irg1 |
| Ccl7 | Gbp11 | Dirc2 | | Mrvi1 |
| Hid1 | Cacna1b | Smim14 | | Rsad2 |

| | | | | |
|---|---|---|---|---|
| Oas3 | Il12a | Clec11a | | Sdc3 |
| Procr | Nkx2-3 | Rhoj | | Gimap6 |
| Gm6377 | Mgarp | Six1 | | Sdc1 |
| Irf7 | Camk2a | Myb | | Bhlha15 |
| Ccl2 | Hmces | Gria3 | | Calcrl |
| Oasl2 | 3425401B19Rik | Dnph1 | | Angpt1 |
| Eya1 | Pigz | Gbp3 | | Cd33 |
| Tbxa2r | Hpgd | Exosc2 | | Csf1 |
| Atp6v0d2 | Dennd5b | Rrp9 | | B3glct |
| Icos | Sox4 | Elf1 | | Gzmb |
| Il5ra | Tsc22d1 | Mpo | | Rasgrp3 |
| Ly6a | Ldoc1l | Ankrd49 | | Tox |
| Spp1 | Gcnt4 | D16Ertd472e | | Tsc22d1 |
| Usp18 | Tnfsf11 | Hspa2 | | Flt3 |
| Ccnd1 | Fgfbp3 | Tsc22d2 | | Fads3 |
| Cd36 | Pygm | Cth | | Lpar1 |
| Piwil2 | Mdm2 | Rev1 | | Fam84b |
| Myct1 | Angpt1 | Atm | | Samd9l |
| Gja5 | Colq | Slc4a5 | | Hmgn3 |
| Rasgef1b | Pcdh7 | Slc7a1 | | Mov10 |
| Scn3a | Atp1a2 | Mgat5 | | Muc13 |
| Gcnt4 | Kansl1l | Slc19a1 | | Armcx1 |
| Rasl11a | Cd34 | Ddx51 | | F2rl3 |
| Lepr | Scn3a | Recql | | Rcor2 |
| Slc35f2 | Sdc4 | Hist1h2an | | Pear1 |
| Ccl3 | Cdkn2c | Frat2 | | Sash1 |
| Hoxa9 | Tctex1d1 | Sgk3 | | Gm12250 |
| Adgrl4 | Cdkn1c | Zbtb1 | | Mllt10 |
| Endou | Il2ra | Cd28 | | Prdx4 |
| Gbp6 | Bcl11a | Ifrd2 | | Abcb1b |
| Slfn5 | Erg | Exo1 | | Tfr2 |
| Sox6 | Gm17745 | Hist1h2ad | | Helz2 |
| Gm5111 | Tubb2b | Prkrir | | Wfdc17 |
| Mira | Hoxa3 | Ramp1 | | Tns1 |
| Xaf1 | Ssbp2 | Xpo4 | | Bmi1 |
| Pim2 | Meis1 | 4933404O12Rik | | Cd93 |
| Lincred1 | Pear1 | Fmr1 | | F13a1 |
| Il2ra | A930004D18Rik | Gemin4 | | Tcf4 |
| Picalm | Slpi | Jmjd1c | | Samsn1 |

| | | | | |
|---|---|---|---|---|
| Abcb4 | Mylpf | Trps1 | | 4921529L05Rik |
| Pcp4l1 | Vangl2 | Chsy1 | | Abca1 |
| Wnt9b | Bcl2 | Otud4 | | Raph1 |
| Abca4 | Trim66 | Dtx4 | | Bend4 |
| Cd27 | Armcx4 | Mgat2 | | Bahcc1 |
| Hoxa7 | Tle2 | Npepl1 | | Mgarp |
| Trpc6 | H2-Ob | Tipin | | Jag2 |
| Lhfpl2 | Pim2 | Ankrd28 | | Gbp11 |
| Hrh2 | Trp53i11 | Flnb | | Plekha6 |
| Irg1 | Bhlha15 | Lars | | Cx3cr1 |
| Mrvi1 | Thsd1 | Myo18b | | Dock9 |
| Fam198b | Angptl2 | Ctsg | | Nlrp1a |
| Rsad2 | Plekha6 | Umps | | Cgnl1 |
| Csgalnact1 | Spp1 | Akap13 | | Prdx1 |
| Cd96 | Mef2c | Armcx4 | | Gm6093 |
| Clnk | Tcf4 | Hoxa10 | | Bnip3 |
| Dusp4 | Ap1m2 | Lmo2 | | Ptprcap |
| Hoxa5 | Nrgn | Tmem229b | | Pias1 |
| Sdc3 | Pdgfrb | Xpo5 | | Adgrg1 |
| Arhgap32 | Clec2f | Atg12 | | Skida1 |
| Gimap6 | Fads3 | Slpi | | Amot |
| Hbb-bt | Hmgn3 | Trmt6 | | Slc39a10 |
| Sdc1 | Ptprcap | Wdr12 | | Endod1 |
| Cysltr2 | Cgnl1 | Elk3 | | Il18rap |
| Mir155hg | Eya1 | Lsm6 | | Rgs11 |
| Bhlha15 | Hoxa9 | Slc16a10 | | Fsd1l |
| Calcrl | Muc13 | Baz1a | | C130046K22Rik |
| Angpt1 | Tnip3 | Nol9 | | 2900026A02Rik |
| Isg15 | Irg1 | Pecam1 | | Selenbp1 |
| Cd33 | Serpina3g | Sypl | | Fstl1 |
| Csf1 | Aff3 | Urb2 | | Mgat4a |
| Kbtbd11 | Gimap9 | Utp20 | | Chd3os |
| Rgs1 | Rgs11 | Ftsj3 | | Rgl1 |
| B3glct | D630045J12Rik | Nop58 | | Carns1 |
| Gzmb | Bahcc1 | Nubp2 | | Slc14a1 |
| Maff | Pdpn | Tshz1 | | B3gnt5 |
| Hmga2 | Piwil2 | Dcun1d1 | | Myo10 |
| Nt5e | Atp2b4 | Rpa2 | | Aldoc |
| Rasgrp3 | Gfi1b | Nat10 | | Nrgn |

| | | | | |
|---|---|---|---|---|
| Tox | Cobll1 | Nop16 | | Camk2a |
| Tsc22d1 | D5Ertd605e | Polr1a | | Adam8 |
| Flt3 | Itgax | Pwp2 | | C030034L19Rik |
| Fads3 | Satb1 | Bace1 | | Arhgef18 |
| Lpar1 | Srgap3 | Etnk1 | | Egln3 |
| Meis1 | Ak4 | Mbnl1 | | Gfi1b |
| Mmp8 | Il7r | Nop2 | | Hoxa3 |
| 6030419C18Rik | Sdc1 | Trmt2a | | A430078G23Rik |
| Fam84b | Tespa1 | Cad | | Zfp760 |
| Parp12 | Fam84b | Dhx37 | | D8Ertd82e |
| Pglyrp2 | Hoxa10 | Nt5dc2 | | Jak3 |
| Fam20a | Ppp1r14c | Ppip5k2 | | Igf2bp2 |
| Mast4 | Ryr1 | Sh3pxd2a | | Sgsh |
| Ssbp2 | Egln3 | 6030458C11Rik | | 1190002N15Rik |
| Lgals3bp | Heg1 | Eef1e1 | | Col4a2 |
| Samd9l | Zbtb16 | Fry | | Tspan13 |
| Slc30a4 | Zfp658 | Ddx18 | | Htra3 |
| Vldlr | Agpat9 | Lsg1 | | Slc4a8 |
| Cdkn2c | Gpr183 | Nek2 | | Pnrc1 |
| Hmgn3 | Lpar1 | Prmt3 | | Scin |
| Map9 | B3gnt5 | Rrp12 | | Tiparp |
| Mov10 | Hoxa5 | Tax1bp1 | | Gimap5 |
| Muc13 | Parp8 | Eftud2 | | Rftn1 |
| Nr1d2 | Ninl | Etv6 | | Cd2ap |
| Prkcq | Shisa2 | Hspa13 | | Lat2 |
| Gdf15 | Armcx2 | Mat2a | | Nanos1 |
| Armcx1 | Ccnd1 | Nol6 | | Capn5 |
| Ctla2b | Six4 | Nol8 | | St8sia4 |
| F2rl3 | Cx3cr1 | Srsf3 | | Klhl24 |
| Map7 | Mmp12 | Tyw1 | | |
| Cd28 | 4921529L05Rik | Gne | | |
| Lpl | Adgrg1 | Mthfd1 | | |
| Rcor2 | Chsy1 | Rsbn1 | | |
| Il1rn | Emp1 | Shmt2 | | |
| Msi2 | Neto2 | Smad5 | | |
| Pear1 | Slc43a1 | Wdhd1 | | |
| Sash1 | Calcrl | Eif1a | | |
| Cpne7 | Hoxa7 | Hist1h3i | | |
| Gm12250 | Myo10 | Morc2a | | |

| | | |
|---|---|---|
| Mllt10 | Samsn1 | Mybbp1a |
| Prdx4 | Six1 | Nmd3 |
| Shank3 | Slc29a3 | Polr1b |
| Abcb1b | Ttc41 | Rsl24d1 |
| Tfr2 | Ccdc102a | Akap11 |
| Mycn | Cd244 | Dnajc11 |
| Slc17a8 | Kcnip3 | Baz1b |
| Pea15a | Plekhg5 | Cyb5b |
| Ctla2a | Ptprk | Mdc1 |
| Helz2 | Spns2 | Mogs |
| Six1 | Amot | Nup107 |
| Irf9 | Nfil3 | Pole |
| Plek | Oaf | Rsl1d1 |
| Wfdc17 | Adgrl4 | Srm |
| Itih5 | Chd3os | Wdr43 |
| Dhx58 | Clec2i | Cbfb |
| Nsf | Ctr9 | Nolc1 |
| Tns1 | Mira | Ppapdc1b |
| Bmi1 | Cd28 | Psmd7 |
| Cd93 | Hes6 | Slc30a5 |
| Etnk1 | Rhobtb3 | Ssr1 |
| Sgk1 | Cacna1f | Adgrg3 |
| Ddx58 | Runx2 | Bclaf1 |
| F13a1 | 2010111I01Rik | Hist1h3g |
| Tcf4 | Cd33 | Ift57 |
| Slc18a2 | Dlg3 | Mcm10 |
| Samsn1 | Tspan13 | Hnrnpab |
| Lrrc32 | Ogdhl | Mthfd1l |
| 4921529L05Rik | Tns1 | Mtr |
| Mx1 | Atg4a | Plac8 |
| Tlr12 | Mylk | Srprb |
| Pdcd1lg2 | Sh2d5 | Syncrip |
| Plekha7 | Slc4a8 | Glipr1 |
| Sulf2 | Armcx1 | Hist2h2ac |
| C1qtnf1 | Dock9 | Ndc1 |
| Myh10 | Kif17 | Arid2 |
| Oas1c | Kmt2a | Atp11b |
| Car13 | B4galnt4 | Fen1 |
| Abca1 | BC064078 | Gemin5 |

| | | |
|---|---|---|
| Raph1 | Thbd | Iars |
| Bend4 | Tmem171 | Ide |
| Bahcc1 | Zfp820 | Psat1 |
| Tfrc | Fam212a | Smc4 |
| Mgarp | Gbp4 | Srgn |
| Ttll7 | St6gal1 | Supt16 |
| Jag2 | Tsc22d2 | Tfrc |
| Gbp11 | Vash1 | Cmtm7 |
| A630033H20Rik | Carns1 | Ctps |
| AA467197 | Ccl4 | Fus |
| Plekha6 | Ipcef1 | Gart |
| Cx3cr1 | Mterf2 | Heatr1 |
| Dock9 | Nanos1 | Ppm1f |
| Prrg4 | Cnksr3 | Sf3b3 |
| Slfn8 | F13a1 | Smg5 |
| Nlrp1a | Mex3b | Spn |
| Cgnl1 | Fam174b | Srsf10 |
| Tlr7 | Slc35e3 | Thrap3 |
| Tnfaip3 | Gpsm1 | Tomm20 |
| Jun | Kank2 | B4galnt1 |
| Prdx1 | Marcksl1 | Bcor |
| Dusp3 | Bnip3 | Dnajc10 |
| Insig1 | Zc3h12c | Larp4 |
| Crebrf | Adck3 | Mdn1 |
| Apbb2 | Gm6093 | Srsf2 |
| Oas1a | Phtf2 | Wdr5 |
| Gm6093 | Selenbp1 | Arid1b |
| Bnip3 | Xpc | Eef2k |
| Dok2 | Fabp4 | Pcdh7 |
| Ptprcap | Stau2 | Pes1 |
| Pias1 | Zfp608 | Rrs1 |
| Adgrg1 | Zscan2 | Thada |
| Gas2l3 | Pbx1 | Aars |
| Skida1 | Rev3l | Arhgap26 |
| Dcbld2 | 2900026A02Rik | Cd24a |
| Plk2 | Bmx | Eif4g2 |
| Amot | Cmpk2 | Mcm5 |
| Erg | Jmjd1c | Nars |
| Slc39a10 | 1190002N15Rik | Pdcd11 |

| | | |
|---|---|---|
| Endod1 | Lhfpl2 | Rpn1 |
| Il18rap | Pbx3 | Birc6 |
| Sel1l3 | Rftn1 | Hist1h2ab |
| Cfap57 | Mov10 | Plekha3 |
| Rgs11 | Afp | Rasgrp2 |
| Fsd1l | Pcp4l1 | Yars |
| Spred1 | Accs | Dgkg |
| Cxcl2 | Chd9 | Ptpn7 |
| C130046K22Rik | Dapp1 | Agpat6 |
| Creb5 | Kcnq5 | Hectd1 |
| Casp12 | Sesn1 | Ikzf1 |
| 2900026A02Rik | Elf1 | Mga |
| Selenbp1 | Frat2 | Abcd2 |
| Ggact | L1cam | App |
| Clstn3 | Zfp518b | Lmnb1 |
| Ifi47 | Abcb1b | Mcm3 |
| Pde11a | Arap2 | Manf |
| Car3 | Arfgef3 | Ran |
| Ehd3 | Bend4 | Sec24d |
| Fstl1 | Nutf2 | Znrf2 |
| Slc22a23 | Prss16 | Atp8b4 |
| Dab2 | Rnf220 | Ncl |
| Mgat4a | Fam60a | Sp9 |
| Fam26f | Jak3 | Cdkn2c |
| Chd3os | Osbpl5 | Tspan12 |
| Fnbp1l | Kdelc2 | Prmt6 |
| Rgl1 | Pecam1 | Kcnk12 |
| Slc22a3 | Acot1 | Pcp4l1 |
| Tgm2 | Ift57 | Msantd3 |
| Gpd1 | Med13l | E2f6 |
| Rhoj | Rab38 | Rcc1 |
| Neurl1b | Reps1 | Chchd4 |
| Carns1 | Slc39a14 | Bora |
| Serpina3g | Bex6 | Hirip3 |
| Crisp3 | Cd69 | Shcbp1 |
| Frmd5 | Elk3 | Ube2q2 |
| Me1 | Nrip1 | Cdc6 |
| Gm15915 | Smim14 | Hist1h2be |
| Slc14a1 | Zfp422 | Ints7 |

| | | |
|---|---|---|
| B3gnt5 | Ccl3 | Tsr1 |
| Eif2ak2 | Kdm5b | Knop1 |
| Myo10 | Rgs12 | Pfas |
| Trim30a | Stxbp4 | Naa40 |
| Hbegf | Tmem229b | Prkdc |
| Gpnmb | Dnajc10 | Stxbp5 |
| Aldoc | Hdac9 | Hn1l |
| Nrgn | P2rx3 | Tars |
| Gpr84 | Rabgap1l | Ifngr1 |
| Stxbp6 | Slc44a1 | Nt5e |
| Camk2a | Trim59 | Ppp5c |
| Spsb4 | Tshz1 | Psmc5 |
| Adam8 | Zfp760 | Sltm |
| Timp3 | Cdk6 | Cdv3 |
| Hsh2d | Fmr1 | Hspd1 |
| Tmem215 | Grap | Mcm6 |
| Tie1 | H2-Q4 | Zc3h4 |
| Ifi27 | Hmgn1 | Lincppara |
| Serpine2 | Hspa2 | Cmpk2 |
| C030034L19Rik | Pafah1b3 | Yars2 |
| Il1r1 | Tle6 | Ruvbl1 |
| Wdfy1 | Aldoc | Tada2a |
| Arhgef18 | Faf1 | Gm17296 |
| Egln3 | Frat1 | Tbl2 |
| Aldh3a1 | Lhfp | Gclc |
| Zfp882 | Mgat5 | Kat2a |
| Col4a1 | Milr1 | Imp4 |
| Akap2 | Endod1 | Hist1h3a |
| Gfi1b | Klhl22 | Srsf7 |
| Tle4 | Adam8 | Cct3 |
| Hoxa3 | Fam162a | Papss2 |
| Plvap | Matk | Sec61a1 |
| 4933430I17Rik | Rere | Abhd17b |
| Bhlhe41 | Rn45s | Anp32e |
| A430078G23Rik | Rnf144a | Serpina3g |
| Afp | Il18rap | Angptl4 |
| Btla | Lpxn | Hgh1 |
| Zfp760 | 5430427O19Rik | Nol10 |
| C530008M17Rik | Mllt10 | E130309D02Rik |

| | | |
|---|---|---|
| Sema7a | Arhgef18 | Ezh2 |
| Cd274 | Baz2b | Isg20l2 |
| Cdk17 | Cnn3 | Cebpz |
| Dnajc10 | Foxp1 | Hist4h4 |
| Phf11b | Slc27a1 | Hist1h4a |
| Kif5a | Spry2 | Set |
| D8Ertd82e | Tia1 | Snrnp200 |
| Enpp1 | Crebzf | Usp1 |
| Casc4 | Cul7 | Serpinb1a |
| Ifi203 | Fam189b | Ercc6l2 |
| Jak3 | Ikzf1 | Nop14 |
| Hivep3 | Mir17hg | Fam76b |
| Cdk14 | Pkib | Lsm8 |
| Igf2bp2 | Ablim1 | C2cd3 |
| Sgsh | Bcor | L3mbtl2 |
| 1190002N15Rik | E130012A19Rik | Tbl3 |
| Ccdc112 | Zfp800 | Nhp2 |
| Col4a2 | Bphl | Ppan |
| Vash2 | Dhrs4 | Mrpl18 |
| Ptgs2os2 | Dusp2 | Pitpnb |
| Luzp1 | Flnb | Sart3 |
| Tspan13 | Raph1 | Nsun2 |
| Tctex1d1 | Tpi1 | Aldh18a1 |
| Htra3 | Tapt1 | Rbbp4 |
| Slc13a2 | Tns4 | Srp68 |
| Slc4a8 | Trim65 | Dnaaf2 |
| C2cd4a | Zfp507 | BC027231 |
| Pnrc1 | Fstl1 | Chtf18 |
| St3gal1 | Jup | Clstn1 |
| Scin | Nlrc5 | 2010111I01Rik |
| Oas1b | Pias1 | Phf5a |
| Tiparp | Pou2f2 | Wdr75 |
| Scimp | Rbfox2 | Hells |
| Gimap5 | Thy1 | Ubr7 |
| Tgtp1 | Atp13a2 | Kbtbd7 |
| Rftn1 | Ccdc134 | Tap2 |
| Pls1 | Cd24a | Sergef |
| Tmem41b | Fam49a | Ahcy |
| Alg11 | Itgb3 | Mms22l |

| | | |
|---|---|---|
| Zbtb6 | Tfrc | Haus6 |
| Cd2ap | Bbx | Ss18 |
| Lat2 | Fut8 | Tatdn2 |
| Arhgef10l | Padi2 | Wars |
| Nanos1 | Tef | Keap1 |
| Fgf3 | Tpm4 | Calr |
| Capn5 | Ube2e3 | G3bp1 |
| Tlr13 | A430078G23Rik | H2afy |
| Cd34 | Arid1b | Fgfbp3 |
| Irgm1 | Fancm | Gng11 |
| St8sia4 | Lat2 | Bag2 |
| Klhl24 | Mier3 | 1700020L24Rik |
| | Rhoq | Gpatch4 |
| | Rnf219 | Hist1h2ag |
| | Sgk3 | Arglu1 |
| | Ap1s3 | Scap |
| | Bcl9l | Slc39a9 |
| | Dnmt3a | Tfdp1 |
| | Gbp3 | Dhx9 |
| | Gm7694 | Eif3b |
| | Kdm3a | Sh2d5 |
| | Kdm4b | Icam2 |
| | Klhl5 | Gle1 |
| | Pgm2 | Pctp |
| | Phf14 | Msh2 |
| | Runx3 | Bckdk |
| | Serpinb1a | Hist1h4k |
| | Zc3h7b | Nup155 |
| | D17H6S56E-5 | Tbc1d9b |
| | Hpse | Srsf11 |
| | Lmo2 | Cd34 |
| | Macf1 | Eif1ad |
| | Pgk1 | Polr3k |
| | Rgl1 | Orc2 |
| | Slc16a3 | Trim56 |
| | Tap2 | Utp15 |
| | Tbc1d4 | Mrpl12 |
| | Zmiz1 | Ern1 |
| | Fam133b | Fam102b |

| | | |
|---|---|---|
| | Il10rb | Pcyox1l |
| | Myb | Chd9 |
| | Pcsk9 | Tfec |
| | Tmem119 | Calm2 |
| | Adam17 | Tnpo1 |
| | Apobec3 | Tomm22 |
| | Arid4b | Shisa2 |
| | Pcyox1l | Ints4 |
| | Pfkl | Xpc |
| | Znrf1 | Kntc1 |
| | Galk1 | Dnajc3 |
| | Galnt6 | Mettl8 |
| | Pan3 | Zfp251 |
| | Pdrg1 | Cldn11 |
| | Xist | Col11a2 |
| | Akt3 | Hist1h3h |
| | Arid1a | Txndc11 |
| | Csnk1g3 | Espl1 |
| | Fam102b | Mcfd2 |
| | Fut7 | Zhx1 |
| | Gsn | Dpp7 |
| | Itga4 | Slc16a1 |
| | Lpgat1 | Eif4a3 |
| | Myef2 | Chtf8 |
| | Nedd4 | Nedd4 |
| | Slc44a2 | Pus7 |
| | Tax1bp1 | Dut |
| | AI504432 | Cep192 |
| | Cd93 | Cnbp |
| | Cdk17 | Pop1 |
| | Phip | Nup133 |
| | Pmaip1 | Ddx21 |
| | Socs2 | Gtf2h1 |
| | Zbtb18 | Nudc |
| | Cdk13 | Zmpste24 |
| | D930015E06Rik | Dhx15 |
| | Ero1l | Rrm1 |
| | Etv6 | Aatf |
| | Klhdc2 | Cdk19 |

| | | |
|---|---|---|
| | Spata13 | Mcm7 |
| | Tmem173 | Srsf1 |
| | Zfp609 | BC003965 |
| | Dcun1d1 | Orc6 |
| | Hnrnph1 | Cactin |
| | Ino80d | Mcph1 |
| | Nnt | Hist2h4 |
| | Smarcc1 | Ranbp1 |
| | Helz2 | Galnt7 |
| | Ube2a | Hist1h1d |
| | Wfdc17 | Hist2h2bb |
| | Atp1b3 | Sgms2 |
| | Slc25a51 | Nfyc |
| | Klhl24 | Nip7 |
| | Otud4 | Ddx24 |
| | Myog | Ltn1 |
| | Gimap5 | Btd |
| | Slc35d3 | Mb21d1 |
| | H2-Ab1 | Elac2 |
| | Cd276 | Elf2 |
| | Kif19a | Adss |
| | Mex3a | Lpgat1 |
| | Cbx2 | Lgalsl |
| | Zfp184 | Pmm2 |
| | Plcb4 | Hnrnph1 |
| | Gzmb | Nme1 |
| | B3glct | Tbccd1 |
| | Il21r | Prmt1 |
| | Repin1 | Ptges3 |
| | Mfge8 | Tpr |
| | Zfp280d | Psmc2 |
| | Arid5a | Mccc1 |
| | Rsbn1l | Erg |
| | Cbfa2t3 | Zbtb18 |
| | Ldha | Slc7a5 |
| | Brap | Cep76 |
| | Egln1 | Man1a |
| | Mdn1 | Nle1 |
| | Gimap7 | Ddx49 |

| | | |
|---|---|---|
| | Cd7 | Cops4 |
| | Sgip1 | Zbtb40 |
| | Serpinb6b | Def8 |
| | Tmprss7 | Hars |
| | Rsad2 | Etfb |
| | Zfp619 | Celf2 |
| | Hdac10 | Gnl3 |
| | Pdk1 | Cstf2 |
| | Kctd1 | Dph5 |
| | Tmem126a | Dkc1 |
| | Naa16 | Chaf1b |
| | Prdx4 | Cirh1a |
| | Fcho1 | Hist1h4b |
| | Ccng2 | Mcm4 |
| | Eef2k | Nup160 |
| | Itgb7 | Skap2 |
| | Pcm1 | Afp |
| | Arid2 | Faap24 |
| | Ube2r2 | Slc35a4 |
| | Serpina3f | Faf1 |
| | Myl4 | Serpinf1 |
| | Prelid2 | Naa30 |
| | Ppp3cc | Bop1 |
| | Arhgef28 | Ccrn4l |
| | Tox | Bckdha |
| | Clstn1 | Tor3a |
| | Nme4 | Rab3gap1 |
| | B230118H07Rik | Zfp91 |
| | Bend3 | Ipo11 |
| | Smim3 | D19Bwg1357e |
| | Wdr6 | Cep250 |
| | B4galt4 | Ddx42 |
| | Zbed4 | E130012A19Rik |
| | Rasa4 | Abhd14a |
| | Jmjd6 | 1810026J23Rik |
| | Dtnbp1 | Polr2a |
| | Uba7 | Abce1 |
| | Pnrc1 | Tmx4 |
| | Mbnl3 | Ssrp1 |

| | | |
|---|---|---|
| | Cxxc5 | Asns |
| | Ccm2l | Ccne2 |
| | Oasl2 | Spsb3 |
| | R3hcc1 | Rbm45 |
| | Tspyl3 | Ttc9c |
| | Rhoh | Zfp800 |
| | Ric8 | Rrm2 |
| | Lrba | Shq1 |
| | Ints7 | Tbp |
| | Gng12 | Otud6b |
| | Casp2 | Smpd4 |
| | Cp | Vps52 |
| | Hdac7 | Soga1 |
| | Rhobtb1 | Ranbp2 |
| | Fbxo21 | Ctu1 |
| | Scin | Pofut1 |
| | Smad5 | Sdad1 |
| | St7 | Ovca2 |
| | Trim28 | Ggcx |
| | Gnptab | Slc7a6 |
| | Etnk1 | Utp6 |
| | Tnrc18 | Smc6 |
| | Pten | Stt3b |
| | Rhoc | Sgsm1 |
| | F2rl2 | Pole4 |
| | Ranbp17 | Pin1 |
| | Eif5a2 | Ppil1 |
| | 2610035D17Rik | Hist1h2ae |
| | Ptpn3 | Abi2 |
| | Gtf2ird2 | Heatr3 |
| | Nsmaf | Hist1h3c |
| | Cnot6l | Btaf1 |
| | Elf2 | Aggf1 |
| | Bckdha | Rtel1 |
| | Ezh2 | Vrk1 |
| | Hhex | Slc44a2 |
| | Chd7 | Osbpl8 |
| | Rtp4 | Hist1h2bp |
| | Abhd6 | Mettl16 |

| | | |
|---|---|---|
| | Plekhh2 | Ndufa9 |
| | Tjp2 | Zmat2 |
| | Sash1 | Ap3s1 |
| | Casd1 | Elmo1 |
| | Ern1 | Farsa |
| | Baz1b | Mybl2 |
| | Galnt12 | Impdh2 |
| | Zfp873 | Psmd13 |
| | Zbtb1 | Rpl7l1 |
| | Zmym6 | Ncbp1 |
| | Bmi1 | Zmym1 |
| | Tiparp | Ccdc69 |
| | Stmn1 | Ntmt1 |
| | Fam65a | Spc24 |
| | C030034L19Rik | Scly |
| | Arg1 | Eaf1 |
| | Zfp773 | Cmtr2 |
| | Xaf1 | Fam133b |
| | Lrrc1 | Cluh |
| | Col4a2 | Gmfb |
| | Slc39a6 | Fasn |
| | Zfp36l2 | Bag5 |
| | Basp1 | Abcb8 |
| | Ckb | Ppp1r8 |
| | Msh6 | Pycr2 |
| | Ndrg1 | Atp2a2 |
| | Akap9 | Derl1 |
| | C2cd3 | Serbp1 |
| | Dido1 | Tctex1d1 |
| | Mif | C1qtnf6 |
| | Zc3h4 | Usp10 |
| | Gm5111 | Elp3 |
| | Atp6v0d2 | Nudt5 |
| | Smagp | Grpel2 |
| | F2rl3 | 3830406C13Rik |
| | Dusp18 | Tk1 |
| | Pdxp | Rrp7a |
| | Fam73a | Xrcc6 |
| | Zfp292 | Chtop |

| | | |
|---|---|---|
| | Tmem14a | Emilin1 |
| | Fbxw4 | Timm17a |
| | Abca1 | 2410016O06Rik |
| | Dirc2 | Mak16 |
| | St8sia4 | Mrpl47 |
| | Anks1 | Ddx39 |
| | Bclaf1 | Coq2 |
| | Ifitm1 | Kcnab2 |
| | Best1 | Luc7l3 |
| | Ly86 | Ddost |
| | Sgsh | Nudt19 |
| | Dennd1c | Zfp422 |
| | Il33 | Crlf3 |
| | Hilpda | Gorasp2 |
| | Nek8 | Casp8ap2 |
| | Fzd7 | Rpa1 |
| | Slco4a1 | Ppid |
| | Traf1 | AI662270 |
| | Ankrd55 | Kcnip3 |
| | Smco4 | Tex10 |
| | Oas3 | Mir703 |
| | Btd | Pmaip1 |
| | Dock10 | Lhpp |
| | Alpi | Slfn9 |
| | Fam214a | Atic |
| | Pwwp2a | Nup214 |
| | Il6st | Tardbp |
| | Sh3d19 | Mns1 |
| | 5530601H04Rik | Polr3e |
| | Map4k3 | Ptdss1 |
| | Ank | Abl2 |
| | Csf1 | Esf1 |
| | Bag2 | Timeless |
| | Smarcad1 | Dek |
| | Il17ra | Ssr3 |
| | Stxbp5 | 2810006K23Rik |
| | Gm16386 | Tmtc3 |
| | Cd320 | Idh3a |
| | L3mbtl3 | Brca1 |

| | | |
|---|---|---|
| | Srek1 | Med1 |
| | Bmyc | Eif3g |
| | Tfr2 | Rpa3 |
| | Lpar2 | Sco1 |
| | Morc2a | Nup85 |
| | Mga | Dhx29 |
| | Trem1 | Zfp638 |
| | Sccpdh | Psmg3 |
| | Cdc42bpg | Emc3 |
| | Pknox1 | Myo19 |
| | Pole2 | Aldh1b1 |
| | Commd7 | Klhdc10 |
| | Abhd17b | Ufd1l |
| | Cxcr4 | Mrpl20 |
| | Mc5r | Rapgef2 |
| | Cpa2 | Alg5 |
| | Armcx6 | Suz12 |
| | B630019K06Rik | Alyref |
| | Slc9a9 | Phtf2 |
| | Zkscan1 | Arhgef10l |
| | Bptf | Sec23ip |
| | Ppcs | Topbp1 |
| | Pla2g6 | Pcgf3 |
| | Lrrc20 | Rbm19 |
| | Fam208a | Ttk |
| | Zbtb40 | Ccbl2 |
| | Tbc1d10c | Tarsl2 |
| | Auh | Ift80 |
| | Ptpn4 | D430020J02Rik |
| | Snx9 | Mios |
| | Eaf1 | Igf1r |
| | Crlf3 | 2610001J05Rik |
| | Tnfsf9 | Gpt2 |
| | Glipr1 | Nudcd2 |
| | Srcap | Gstm2 |
| | A630072M18Rik | Plaa |
| | Gyltl1b | BC031181 |
| | Prdm11 | Zcchc17 |
| | Bambi | Cse1l |

| | | |
|---|---|---|
| | Cntnap1 | Hnrnpa0 |
| | Slc39a10 | Sppl2b |
| | Abl2 | Tubb4b |
| | Mlf1 | Mrpl50 |
| | Usp11 | Spata5 |
| | Cd2ap | Tmem97 |
| | Tes | Pola1 |
| | Fubp1 | Setdb2 |
| | C030039L03Rik | Clp1 |
| | BC147527 | U2af1 |
| | D8Ertd82e | Ipo8 |
| | Rccd1 | Ciapin1 |
| | Fndc3a | Nop56 |
| | 3110052M02Rik | Toe1 |
| | Glg1 | Traf2 |
| | Itgb5 | Atad3a |
| | Mgea5 | Abcf1 |
| | Clybl | Adar |
| | Nlgn2 | Cacybp |
| | Cdyl2 | Idi1 |
| | Ncoa5 | Wdr74 |
| | Psmb8 | Smyd2 |
| | Tifab | Tbrg4 |
| | Taf9b | Nup50 |
| | Igf2r | Il21r |
| | Cntln | Mtrr |
| | Ccdc180 | Sprtn |
| | Gm12250 | Vcp |
| | Otos | Anapc1 |
| | Paqr8 | Ncoa5 |
| | Ctnnal1 | Scnn1a |
| | Kmt2d | Fastkd5 |
| | Tcf3 | Etv4 |
| | 3300005D01Rik | Atr |
| | A230050P20Rik | Hyou1 |
| | Rhoj | Fam219b |
| | Cblb | Lyar |
| | Brd1 | Paf1 |
| | Eif4g2 | Exosc6 |

| | | |
|---|---|---|
| | B3gnt7 | Slc9a8 |
| | Sema4c | Ptgr2 |
| | Pogz | Scamp3 |
| | Chst1 | Ogfod1 |
| | Ms4a4b | Fam107b |
| | Zfp113 | Rad51ap1 |
| | Ociad2 | Spsb4 |
| | Msl3l2 | Tmem161b |
| | Dsp | Arf4 |
| | Atxn7 | Retsat |
| | Runx1 | Smarcad1 |
| | Gna15 | Rps6kb1 |
| | Tbc1d5 | Cks1b |
| | Gpr18 | Zpr1 |
| | Gbp6 | Rnf219 |
| | Grasp | Ttc7 |
| | Ccdc149 | Gars |
| | Cmtm7 | Hmces |
| | Ddah2 | Helz |
| | Ift81 | Rbsn |
| | BC005561 | Ogfr |
| | Mier2 | Irgm1 |
| | Samd9l | Nom1 |
| | Myl10 | Mrs2 |
| | Nol9 | Srf |
| | Ttc27 | Cmtm8 |
| | Rasal1 | Fbxo30 |
| | Kcng2 | Atad5 |
| | Ccbl2 | Ncoa4 |
| | Zfp266 | Pdap1 |
| | Elmo1 | Eef1d |
| | Tmem86b | Metap2 |
| | Cnp | Xpot |
| | Fis1 | Rbck1 |
| | Pole | Alg14 |
| | Mir8102 | Lonrf3 |
| | Poll | Chd1l |
| | Zfp518a | Hccs |
| | Igf2bp2 | Zdhhc6 |

| | | |
|---|---|---|
| | Net1 | Trak2 |
| | Mbnl1 | Gmpr2 |
| | Ints1 | Car9 |
| | Kcna3 | Josd2 |
| | Mafk | Tmem39a |
| | Capn5 | Tmem248 |
| | Chd3 | Plekhh2 |
| | Rmdn2 | Urb1 |
| | Thg1l | Mtfr1 |
| | Mbtd1 | Gmps |
| | Pcdhgb6 | Cbx3 |
| | Pdcd4 | Tcof1 |
| | Wtip | Trip13 |
| | 5830416P10Rik | Agap2 |
| | Frmd4b | Pea15a |
| | Rcbtb1 | Prpf3 |
| | Taf8 | Galnt1 |
| | Golm1 | Fut11 |
| | Cops4 | Mlec |
| | Adgrg3 | Alkbh1 |
| | Irf7 | Tmem186 |
| | Afap1l1 | Telo2 |
| | Tspan2 | Ssna1 |
| | Ino80c | Park7 |
| | Cdk4 | Inip |
| | Efna4 | Hist1h1e |
| | Glra3 | Apmap |
| | Slc14a1 | Slc25a10 |
| | Ip6k2 | Cpsf6 |
| | Dcakd | Nsf |
| | Msh2 | Hdac8 |
| | 2610008E11Rik | Fignl1 |
| | Ppp1r3b | Uhrf1 |
| | 4931428F04Rik | Kpnb1 |
| | Gpr171 | Mettl2 |
| | Nudt1 | Glcci1 |
| | Zxda | Cpeb2 |
| | Sdc3 | Wdfy1 |
| | Tmem181b-ps | Ano6 |

| | | |
|---|---|---|
| | Brpf3 | Txlna |
| | Hnrnpm | Rcc2 |
| | Ccdc157 | Trpm7 |
| | Ciart | Nxt1 |
| | Txlna | Champ1 |
| | Myadml2 | Pcna |
| | C130046K22Rik | Tsen2 |
| | Suv420h1 | Yipf6 |
| | Zc3hav1l | Dhx33 |
| | Gtf3c6 | Prpf19 |
| | Ramp1 | D030028A08Rik |
| | Tcf12 | Cwf19l1 |
| | Ptpra | Pa2g4 |
| | Prdx1 | Zranb2 |
| | Nt5e | Slfn3 |
| | Sfpq | Adsl |
| | Rfx7 | Harbi1 |
| | Btnl9 | Rad1 |
| | 4931414P19Rik | Phldb3 |
| | Lin37 | Nol11 |
| | Pigh | Nup153 |
| | 2810417H13Rik | Pcnt |
| | Ppfia4 | Pole2 |
| | Vwf | Ubtf |
| | Kansl2 | Cxcl10 |
| | Eldr | Pusl1 |
| | B4galnt2 | Pom121 |
| | Ccdc80 | Hist1h1b |
| | Dnph1 | Zfp180 |
| | Abracl | Fem1a |
| | Cdkn1b | Arhgap11a |
| | E130307A14Rik | Psmc4 |
| | Isoc2b | Wdr4 |
| | Nxpe3 | Dcaf8 |
| | Tubb4a | Pik3r1 |
| | Gpc2 | Ilf3 |
| | Wipi1 | Ythdf2 |
| | Agbl3 | Prmt7 |
| | Tnfsf8 | Lrrc40 |

| | | |
|---|---|---|
| | Fsd1l | Manba |
| | Cstf2 | Zbtb25 |
| | Lrif1 | Tmem263 |
| | Prmt5 | Ubn1 |
| | Erdr1 | Polr2b |
| | Commd9 | Rpp40 |
| | Fam65b | Dnase2a |
| | Kcne3 | Slc46a1 |
| | 3110057O12Rik | Wdr8 |
| | Agap2 | Vprbp |
| | Pum1 | Slc16a6 |
| | Bmp8a | Psph |
| | D3Ertd751e | Rad51 |
| | Gtf3c5 | 2700097O09Rik |
| | Fbp1 | Erbb3 |
| | Phldb1 | Cnot11 |
| | Sh3pxd2a | Zfp410 |
| | Ralgps1 | Pex2 |
| | Katnal1 | E2f1 |
| | Pde3b | Apex1 |
| | 5830416I19Rik | Slc4a2 |
| | Hltf | Galk1 |
| | Lrp10 | Ncapg |
| | Cers4 | Tsen15 |
| | N4bp2 | Anapc16 |
| | Snn | Nup205 |
| | AI314180 | Mmp8 |
| | Numa1 | Fubp1 |
| | Sms | 0610007P14Rik |
| | Haus4 | Ncbp2 |
| | Mgat4a | Rps27l |
| | Ccdc94 | Adrm1 |
| | Wiz | Abracl |
| | Grhpr | Slc39a7 |
| | 2810408A11Rik | Smc3 |
| | C920025E04Rik | Prpf31 |
| | Etl4 | Gtpbp3 |
| | Coprs | Mcm3ap |
| | Cenpv | Prpf4 |

| | | |
|---|---|---|
| | D16Ertd472e | Dis3 |
| | Siae | Slc35b2 |
| | Ulk1 | Mrpl33 |
| | AI429214 | Ncapg2 |
| | Acin1 | 9130011E15Rik |
| | Tmprss3 | Edem1 |
| | Zfp963 | Slc38a7 |
| | Zfp808 | Vimp |
| | H2-Oa | Snhg5 |
| | Nrde2 | Rangap1 |
| | Polr2a | Pus1 |
| | Rps27 | Slc39a6 |
| | Zfp455 | Srrt |
| | Dock6 | Ddx27 |
| | Pced1b | Hnrnpm |
| | Gria3 | Cog7 |
| | 1810006J02Rik | Hist3h2a |
| | A630001G21Rik | Slc33a1 |
| | Tdrkh | Akap1 |
| | Klf2 | Actr6 |
| | Mb21d1 | Tmed9 |
| | Smg6 | Uspl1 |
| | Cd22 | Tcerg1 |
| | H2-T10 | Stmn1 |
| | D11Wsu47e | Mina |
| | Hp1bp3 | C1qbp |
| | Xlr4b | Ung |
| | Cd27 | Dhrs4 |
| | Rab11fip4 | Zfp346 |
| | Dis3l | Ddx3x |
| | Bmpr2 | Casp2 |
| | E2f7 | Psme3 |
| | Ung | Zfp954 |
| | Nt5c2 | |
| | Zfp644 | |
| | Pomgnt2 | |
| | Abl1 | |
| | H2afy | |
| | Fmnl3 | |

| | |
|---|---|
| | Tnrc6a |
| | Wdr54 |
| | Parp11 |
| | Spats2 |
| | Cnot8 |
| | Fbxl20 |
| | Colec12 |
| | Tap1 |
| | Phlpp2 |
| | Ly9 |
| | Hirip3 |
| | Mef2d |
| | AW146154 |
| | Spag4 |
| | Pcx |
| | Akap13 |
| | Chst7 |
| | Pgam1 |
| | 1110002L01Rik |
| | Zhx2 |
| | Klra1 |
| | Ppp6r2 |
| | Iqsec2 |
| | Icam2 |
| | Rnf150 |
| | Azin2 |
| | Car15 |
| | Plec |
| | Sptbn1 |
| | Zfp937 |
| | Ms4a6b |
| | Enpp4 |
| | Tango6 |
| | Jmjd4 |
| | 4931406P16Rik |
| | Crtc1 |
| | Hesx1 |
| | Nlrp1a |
| | Ddx17 |

| | |
|---|---|
| | Zfp105 |
| | Pik3r1 |
| | 2810410L24Rik |
| | Jmy |
| | B3gnt2 |
| | Rasgrp3 |
| | Rbmx |
| | Taf1 |
| | Pdlim7 |
| | Fryl |
| | Sema4d |
| | Erbb2ip |
| | Pik3cg |
| | Epc1 |
| | Ubfd1 |
| | Guca1b |
| | Hmgxb4 |
| | 2010320M18Rik |
| | Clspn |
| | Wdr13 |
| | Rfwd3 |
| | Top2b |
| | Cela1 |
| | Rnf217 |
| | Syne4 |
| | Glyctk |
| | 5830415F09Rik |
| | Gdf11 |
| | Fam213a |
| | Dnajb5 |
| | Pgm2l1 |
| | Htra3 |
| | Cyth4 |
| | Gstm5 |

**TABLE S2**

| SE Linked Genes in CALM-AF10 On | SE Linked Genes in CALM-AF10 Off |
|---|---|
| Capn11 | Prkrip1 |
| Il10 | Ptrh2 |
| Eif2d | Ddc |
| Gadd45g | 1700042O10Rik |
| Cdc16 | D630003M21Rik |
| Cacna2d4 | Bpi |
| Mir1965 | 2010009K17Rik |
| Phactr1 | Gm16548 |
| Prlh | F5 |
| Mir466b-2 | Zbtb17 |
| Myl3 | Gm694 |
| Spns3 | Dhrs3 |
| Gng12 | Fnbp1 |
| E230016M11Rik | D330023K18Rik |
| Gadd45a | BC005624 |
| Kbtbd7 | Usp20 |
| Zbtbd6 | Gpr107 |
| Dars | P2ry6 |
| Mcm6 | Calr3 |
| Lct | 1700030K09Rik |
| Gm17296 | Slc22a16 |
| Abhd6 | Adam30 |
| Sirt5 | Ephx2 |
| Atg16l1 | Chrna2 |
| Aph1b | Tmem39b |
| Actg1 | Kpna6 |
| 0610009L18Rik | Rpn1 |
| Fscn2 | Rab7 |
| Map2k4 | Gm5577 |
| Mir744 | H1fx |
| Zkscan6 | Acsf3 |
| Tatdn1 | Pcdhgc4 |
| Ndufb9 | Pcdhgc3 |
| Slc22a5 | Arap3 |
| 1700025N23Rik | Pcdhgc5 |
| Gm13704 | Fam83a |

| | |
|---|---|
| Chrna1 | Heatr5a |
| Sun1 | Pdss1 |
| Mlh3 | Cfl2 |
| Elf2 | 1600020E01Rik |
| Ccrn4l | Pcbp1 |
| 4930577N17Rik | 2310040G24Rik |
| Dcaf7 | Mxd1 |
| Icmt | Asprv1 |
| Hes3 | Ifnar2 |
| AB124611 | Il10rb |
| Kptn | Cd247 |
| Snord23 | 1700066B19Rik |
| Gltscr2 | Tmem173 |
| Wisp1 | Ecscr |
| Gal | Ell2 |
| Ppapdc1b | 9130017N09Rik |
| Numa1 | Dusp7 |
| Rnf121 | Zzef1 |
| Il18bp | Srrm1 |
| Trpc2 | Ncmap |
| Lrrc51 | Kdm2a |
| Tomt | Spop |
| Lamtor1 | Mtmr3 |
| Anapc15 | Ascc2 |
| Rptor | Gm11961 |
| Tmem60 | Mcmdc2 |
| Phtf2 | Snord87 |
| Mau2 | Sav1 |
| Sugp1 | 4930430J02Rik |
| Pex11c | Mgat4b |
| Zfp358 | Pak1ip1 |
| Mcoln1 | Tead3 |
| 1700019B03Rik | Tsen2 |
| Crebrf | Slc25a12 |
| Slc45a1 | Dync1i2 |
| Mir582 | Hat1 |
| Zeb2 | Sowahb |
| Gm13476 | Irak2 |
| Mir5129 | Brk1 |

| | |
|---|---|
| Irak3 | Vhl |
| Tmbim4 | Ghrl |
| Helb | Tatdn2 |
| Llph | Fancd2os |
| Cnksr3 | Snora23 |
| Igfbp7 | Rplp0 |
| Noa1 | Pxn |
| Polr2b | Sirt4 |
| Lrrc33 | Pla2g1b |
| Bex6 | Gcn1l1 |
| H1foo | Luc7l2 |
| Plxnd1 | 1110001J03Rik |
| Rho | Klrg2 |
| 4931406H21Rik | Grip2 |
| Mecom | Chchd1 |
| Nadk | Fut11 |
| Gnb1 | 6230400D17Rik |
| Tmem52 | Zswim8 |
| Slc35e2 | Anxa9 |
| Id2 | Fam63a |
| Rgmb | Prune |
| Anp32b | Cers2 |
| Hemgn | Mllt11 |
| Gm17745 | Cdc42se1 |
| Elk3 | Bnipl |
| Tmem150b | Gm128 |
| Wdfy4 | Acox3 |
| Arhgap22 | Htra3 |
| Mtfr1l | Sh3tc1 |
| 4930431P03Rik | Trmt44 |
| 2010109A12Rik | 4931431C16Rik |
| Ccni | Eif2ak4 |
| Pi16 | Blm |
| Dip2b | Furin |
| Atf1 | Fam120aos |
| Tmprss12 | Fam120a |
| Mettl7a1 | Phf2 |
| Mir3473 | Mthfd2 |
| Tmem66 | Mob1a |

| | |
|---|---|
| Mboat4 | Elovl6 |
| Leprotl1 | Ptger1 |
| Dctn6 | Fam103a1 |
| Pin4 | Map3k6 |
| Ptpn2 | Hp1bp3 |
| Cep76 | Wfdc5 |
| Psmg2 | Kcns1 |
| Seh1l | Wfdc12 |
| Spink4 | Ide |
| Gpr68 | 9030617O03Rik |
| Frmd4b | BC005537 |
| Oscar | Tdp2 |
| Ndufa3 | D130043K22Rik |
| Tfpt | Acot13 |
| Prpf31 | Mgat4a |
| Leng1 | 4930594C11Rik |
| Tmc4 | Unc50 |
| Cnot3 | Coa5 |
| Mir3572 | Psmb7 |
| Metrnl | Fnbp4 |
| B3gntl1 | Nup160 |
| Ropn1l | Agbl2 |
| Jund | Atxn7l1 |
| Lsm4 | Efcab10 |
| Gm3336 | Twistnb |
| Pde4c | Zfand2a |
| Gm12504 | 4930500L23Rik |
| Dusp4 | Traf3 |
| Nudc | Amn |
| Nr0b2 | Ddx19a |
| A330035P11Rik | Aars |
| Tm9sf2 | Mir3473d |
| Txnl4a | Sf3b3 |
| Rbfa | Ddx19b |
| Hsbp1l1 | Cog4 |
| Pqlc1 | Arhgef2 |
| Adnp2 | Ssr2 |
| Rab20 | 2810403A07Rik |
| Bcl2l11 | Ubqln4 |

| | |
|---|---|
| Zfr | Rxfp4 |
| Mtmr12 | Rab25 |
| Mir1898 | Lamtor2 |
| Tfeb | Mir1905 |
| Mdfi | Pqlc3 |
| Pgc | Sltm |
| Gm14873 | Rnf111 |
| Msx3 | Cul3 |
| Selplg | 1700016L21Rik |
| Tmem119 | Usp48 |
| Sart3 | Ldlrad2 |
| Iscu | Rap1gap |
| Jun | Gata2 |
| Pik3ip1 | Dnajb8 |
| Limk2 | Tbc1d23 |
| Gng7 | Nit2 |
| Gadd45b | Tomm70a |
| Lmnb2 | Tmem30c |
| Diras1 | 4933411E08Rik |
| Tmprss9 | Tmem184b |
| Slc39a3 | Csnk1e |
| Timm13 | Mir1943 |
| Enah | Kcnj4 |
| Slc2a1 | Maff |
| Zfp691 | Plekhf2 |
| Lrrc32 | Ndufaf6 |
| Sumo3 | Erbb2ip |
| Tbcc | Hsd11b1 |
| Prph2 | Traf3ip3 |
| Gltscr1l | A130010J15Rik |
| Ptpra | G0s2 |
| Pced1a | Irf6 |
| 4930473A02Rik | Lamb3 |
| Mrps26 | Diexf |
| Vps16 | Tac4 |
| Hmg20b | Cbl |
| Rbm39 | Mcam |
| Phf20 | Nlrx1 |
| Romo1 | Rnf26 |

| | |
|---|---|
| Nfs1 | Ccdc153 |
| Unc13d | Pdzd3 |
| Wbp2 | Arid5b |
| Unk | 4930545H06Rik |
| H3f3b | Tmem71 |
| Trim65 | Phf20l1 |
| Mrpl38 | Lrrc6 |
| Trim47 | Mettl21a |
| Galk1 | Sfi1 |
| Cd33 | Pisd-ps1 |
| Iglon5 | Trem1 |
| Zfp658 | Mir706 |
| 4931406B18Rik | Wnk1 |
| Zfp719 | Rhd |
| Herpud1 | Sh3bp1 |
| Slc12a3 | Gga1 |
| Mir138-2 | Lgals1 |
| 9330175E14Rik | Cdc42ep1 |
| 4930529L06Rik | Nol12 |
| Dnaja1 | Lgals2 |
| Aptx | Triobp |
| Mir207 | Pdxp |
| Gm6297 | Timm22 |
| Sema3b | Ap3b1 |
| Gnai2 | Gm9776 |
| Slc38a3 | Ubap2 |
| Rassf1 | Ube2r2 |
| Ifrd2 | Aqp3 |
| Tusc2 | Nol6 |
| Nat6 | Raver2 |
| Hyal1 | Cacng8 |
| Gnat1 | Cacng6 |
| Hyal2 | Tarm1 |
| Hyal3 | Eif4enif1 |
| Armc9 | Ggt1 |
| Gimap6 | Arl11 |
| Gimap9 | 1700109G14Rik |
| Gimap4 | Ebpl |
| Gimap8 | Sgcg |

| | |
|---|---|
| Gimap5 | Lmnb1 |
| AI854703 | Ints7 |
| Gimap7 | Dtl |
| Gimap1 | Phf17 |
| Bin2 | Rap1a |
| Cela1 | Fam212b |
| Galnt6 | Gm5547 |
| Smagp | Ddx20 |
| C330013E15Rik | Hmgcr |
| Dazap2 | Csf1 |
| Tob2 | Tpd52l2 |
| Tef | Dnajc5 |
| Aco2 | Zbtb46 |
| Phf5a | Gm16119 |
| Tnpo2 | Abhd16b |
| Fos | Prpf6 |
| Sp2 | Samd10 |
| D030028A08Rik | Znf512b |
| Sp6 | Uckl1 |
| Pnpo | Prune2 |
| Cdk5rap3 | Slc35c2 |
| Prr15l | Elmo2 |
| Gcnt4 | Cdh22 |
| 1700029F12Rik | Zfp663 |
| Cxcl11 | Gm1587 |
| Art3 | Dnajc15 |
| Sdad1 | Epsti1 |
| Cxcl10 | Nedd4 |
| Cxcl9 | Ablim1 |
| Arpp19 | Matr3 |
| Myo5a | Paip2 |
| 1700113A16Rik | Snhg4 |
| Mef2d | Slc23a1 |
| Mir9-1 | Snora74a |
| Mir3093 | Mir1949 |
| Cnp | Mzb1 |
| Dnajc7 | Spata24 |
| Nkiras2 | Prob1 |
| Zfp385c | Psmd6 |

| | |
|---|---|
| Acly | Atxn7 |
| Ttc25 | Krt83 |
| Dnmbp | Krt7 |
| Cpn1 | Csnk1g2 |
| Sdc1 | Prdm2 |
| Rprd2 | 5031414D18Rik |
| Ecm1 | Gm4278 |
| Tars2 | Lrch1 |
| Clmp | Arf6 |
| Hspa8 | Nemf |
| Ndufb7 | Klhdc2 |
| Tecr | Xpo6 |
| Dnajb1 | Gsg1l |
| Olfr370 | Hs2st1 |
| Gipc1 | 15-Sep |
| Astl | Fhod3 |
| Stard7 | Nup210 |
| Dusp2 | Hdac11 |
| Adra2b | Esr1 |
| Tmem127 | Rps15a |
| Ciao1 | Smg1 |
| Rbl1 | Arl6ip1 |
| Samhd1 | Stau1 |
| Tldc2 | Cse1l |
| 9830001H06Rik | Ddx27 |
| Ldha | Cd84 |
| Gtf2h1 | Lrrc43 |
| Saa2 | Nop10 |
| Hps5 | Slc12a6 |
| Ldhc | Lpcat4 |
| Yy1 | Nutm1 |
| Mir345 | Gmfg |
| Slc25a47 | Polr2e |
| Slc25a29 | Gpx4 |
| 5430416N02Rik | Hmha1 |
| Lin54 | Dot1l |
| Sec31a | 2010016I18Rik |
| Atp1b3 | Dennd2d |
| Gm5148 | Cept1 |

| | |
|---|---|
| Rassf6 | Dram2 |
| Abcg1 | BC051070 |
| Tff2 | Ppp1r15b |
| Tff3 | Defb38 |
| Rufy1 | Igsf6 |
| Hnrnph1 | Mettl9 |
| Mir804 | Otoa |
| Canx | Pds5a |
| Arhgap18 | Dhx9 |
| St7 | Shcbp1l |
| Ttc33 | Npl |
| Prkaa1 | E330020D12Rik |
| Ptger4 | Apobec1 |
| Rpl37 | Aicda |
| Card6 | Mfap5 |
| Snord72 | Dppa3 |
| H2afv | Gdf3 |
| Zmiz2 | Tmem154 |
| Ppia | Tigd4 |
| Purb | Wdfy1 |
| Cdk12 | Mrpl44 |
| Med1 | Mcph1 |
| Fbxl20 | Agpat5 |
| Mir5119 | Angpt2 |
| Nfyb | Pum1 |
| 1700028I16Rik | Sdc3 |
| Txnrd1 | Snord85 |
| Hcfc2 | 2310050B05Rik |
| 2210018M11Rik | Coq2 |
| Prkrir | Hpse |
| Gm9958 | Nisch |
| Ankrd17 | Sema3g |
| Sdccag8 | Bap1 |
| Hmga2-ps1 | Stab1 |
| Cep170 | Smim4 |
| Mir350 | Nt5dc2 |
| Parp6 | Tnnc1 |
| Pkm | Phf7 |
| Gramd2 | Usp47 |

| | |
|---|---|
| Rilp | C87436 |
| Itgb4 | Rnf115 |
| Fam129b | Polr3c |
| Stxbp1 | Nudt17 |
| Snora65 | Ankrd35 |
| Lrsam1 | Pias3 |
| Rpl12 | Cd160 |
| Actn4 | Bach1 |
| Lgals7 | Bid |
| Eif3k | Bcl2l13 |
| Map4k1 | Atp6v1e1 |
| Capn12 | Slc16a10 |
| Vdac1 | Dhrs13 |
| 9530068E07Rik | Phf12 |
| Tcf7 | Flot2 |
| Klhl18 | Ccr7 |
| Ptpn23 | Tns4 |
| Kif9 | Gpr114 |
| Ngp | Ccdc102a |
| Slc38a10 | Vwa8 |
| 1810043H04Rik | Lsm12 |
| Enthd2 | G6pc3 |
| Slc44a2 | Hdac5 |
| Kri1 | Nags |
| Atg4d | BC030867 |
| Ilf3 | Asb16 |
| Cdkn2d | Tmem101 |
| Ap1m2 | Tmem232 |
| Scand1 | Hpcal1 |
| Cnbd2 | Myg1 |
| Tgfbr2 | Iqgap2 |
| Ccl9 | Foxd2 |
| E230016K23Rik | 9130206I24Rik |
| Ccl5 | Foxe3 |
| Gm11435 | Defb36 |
| Ccl6 | Tmem63a |
| S1pr3 | Lefty1 |
| Ptgir | Ephx1 |
| Calm3 | P4hb |

| | |
|---|---|
| Pnmal1 | Gcgr |
| Dact3 | Ppp1r27 |
| Pnmal2 | Fam195b |
| Gng8 | Serbp1 |
| Cep85l | Il12rb2 |
| Fam20a | Atp8a1 |
| Prkar1a | Aatk |
| Fhl3 | Cpsf2 |
| Utp11l | Trip11 |
| Mir698 | Atxn3 |
| Inpp5b | Ralbp1 |
| Sf3a3 | Elavl1 |
| Pou3f1 | Ccl25 |
| Mir697 | Timm44 |
| Senp6 | Map2k7 |
| 3110062M04Rik | Gm14378 |
| Tmem140 | Ctxn1 |
| Stra8 | Snapc2 |
| Wdr91 | Parp9 |
| Kif2a | Dtx3l |
| 3830408C21Rik | Kpna1 |
| Dimt1 | Parp14 |
| 4933426D04Rik | 6330418K02Rik |
| Smad7 | Lamtor4 |
| Hecw2 | BC037034 |
| Stk17b | Gal3st4 |
| Afap1l1 | Gpc2 |
| 1500015A07Rik | Stag3 |
| Pcyox1l | Nxpe5 |
| Grpel2 | Sidt2 |
| Wdr26 | Tagln |
| Ell | Pafah1b2 |
| 2810428I15Rik | Sik3 |
| Fkbp8 | Vps37c |
| Kxd1 | 4930524O05Rik |
| Uba52 | A430093F15Rik |
| Prkcq | Pga5 |
| Mns1 | Cd5 |
| Tex9 | Slc29a1 |

| | |
|---|---|
| 4930509E16Rik | Nfkbie |
| Rfx7 | Slc35b2 |
| Rnf169 | Hsp90ab1 |
| Chrdl2 | Aars2 |
| Eps8 | Gm7325 |
| Ugt1a2 | Tcte1 |
| Dnajb3 | Tmem151b |
| Ugt1a1 | Zmat5 |
| Prr14 | Uqcr10 |
| Fbrs | Gpr179 |
| Zfp688 | Socs7 |
| 1700008J07Rik | Arhgap23 |
| Zfp689 | Hif1a |
| Snora30 | Snapc1 |
| Trappc6a | Slc20a2 |
| Bloc1s3 | Smim19 |
| Nkpd1 | Zdhhc3 |
| A930016O22Rik | Exosc7 |
| Mark4 | Tgm4 |
| Csf1r | Clec3b |
| Hmgxb3 | Nxph3 |
| Slc26a2 | Dgke |
| Pde6a | Trim25 |
| Klhl25 | Gm525 |
| A930015D03Rik | Scpep1 |
| H2-T24 | Gm15698 |
| Gm6034 | 2210409E12Rik |
| Prr3 | Coil |
| Gnl1 | 1810013L24Rik |
| Gm11127 | Tmem40 |
| H2-T9 | Snora7a |
| Mir877 | Rpl32 |
| Abcf1 | Cand2 |
| H2-T23 | Tia1 |
| H2-T22 | Fam136a |
| Pip4k2b | Snrpg |
| Psmb3 | Pcyox1 |
| Pcgf2 | Kit |
| Mllt6 | Mast3 |

| | |
|---|---|
| Cisd3 | Il12rb1 |
| Fam160b1 | Pik3r2 |
| B230217O12Rik | Rab3a |
| Tmem176b | Ifi30 |
| Gimap3 | Arrdc2 |
| Tmem176a | Mpv17l2 |
| Kcmf1 | Kcnn1 |
| Lif | 2010320M18Rik |
| Bcar3 | Gm15850 |
| Mir760 | Kif21b |
| Tnfaip3 | 0610039K10Rik |
| Arid3b | Serinc3 |
| 4933406C10Rik | Pkig |
| Sypl | Ttpal |
| Gng2 | Fam155a |
| Clic1 | Prkcd |
| Msh5 | Mir3076 |
| Sapcd1 | Rft1 |
| Ly6g6d | Anln |
| Vars | Mir680-2 |
| Vwa7 | 9530077C05Rik |
| G6b | Gm6607 |
| AU023871 | Zfp810 |
| Abhd16a | Edc3 |
| Ddah2 | Cyp1a1 |
| Ly6g6f | Cyp1a2 |
| Ly6g6c | Pla2g15 |
| Ly6g6e | Slc7a6 |
| Gpcpd1 | Esrp2 |
| AU019990 | Slc7a6os |
| Stap1 | 1810019D21Rik |
| Cenpc1 | Prmt7 |
| Gm5464 | Smim6 |
| Dpysl2 | Recql5 |
| Pnma2 | Smim5 |
| Cdc42se2 | Sap30bp |
| Gm12228 | Stt3b |
| Lrrc27 | 4930428G15Rik |
| Stk32c | Stmn1-rs1 |

| | |
|---|---|
| Pwwp2b | Il15ra |
| Matn4 | Anxa2 |
| Rbpjl | Mir3109 |
| Slpi | Narg2 |
| Svs5 | Ppp2r4 |
| Svs6 | Dolpp1 |
| Snx29 | Crat |
| | Ier5l |
| | Fam73b |
| | Sh3glb2 |
| | Nt5e |
| | Snx14 |
| | Mrpl48 |
| | Plekhb1 |
| | Rab6a |
| | Rnf149 |
| | Creg2 |
| | Frmd4a |
| | Jam2 |
| | Atp5j |
| | Gabpa |
| | Mrpl39 |
| | Rprd1b |
| | Tti1 |
| | Pak6 |
| | Plcb2 |
| | Ankrd63 |
| | Phgr1 |
| | 5430417L22Rik |
| | A430105I19Rik |
| | Disp2 |
| | Rab10 |
| | 1700012B15Rik |
| | Ptp4a3 |
| | Gpr20 |
| | Suco |
| | Hist1h2be |
| | Hist1h4d |
| | Hist1h4h |

| | |
|---|---|
| | Hist1h3f |
| | Hist1h2ae |
| | Hist1h2af |
| | Hist1h2ad |
| | Hist1h3g |
| | Hist1h2bg |
| | Hist1h2bh |
| | Hist1h3d |
| | Hist1h1d |
| | Hist1h4f |
| | Hist1h2bf |
| | Hist1h3e |
| | Hist1h1e |
| | Calcrl |
| | Tfpi |
| | 2810006K23Rik |
| | Cdk2ap1 |
| | Mphosph9 |
| | Mthfd1 |
| | Tex21 |
| | Slc7a8 |
| | Cebpe |
| | Homez |
| | Ccdc138 |
| | Ak2 |
| | Adc |
| | Rnf19b |
| | Lpo |
| | Mks1 |
| | Mlec |
| | Acads |
| | Cabp1 |
| | Gm13826 |
| | Unc119b |
| | Klhl6 |

### TABLE S3

| Cytokine | CALM_AF10_On_1 | CALM-AF10_On_2 | CALM_AF10_Off_1 | CALM-AF10_Off_2 | On_Avg | Off_Avg |
|---|---|---|---|---|---|---|
| **M-CSF** | 1642.96875 | 1734.25 | 564.484375 | 746.796875 | 1688.60938 | 655.640625 |
| **GM-CSF** | 3016.4375 | 2781.67188 | 452.140625 | 583.078125 | 2899.05469 | 517.609375 |
| **IL-3** | 11279.6094 | 11558.9844 | 9349.29688 | 9402.26563 | 11419.2969 | 9375.78125 |
| **IL-6** | 6429.07813 | 6663.15625 | 5407.42188 | 5406.09375 | 6546.11719 | 5406.75781 |
| **CCL17/TARC** | 27540.8438 | 27056.3438 | 20757.125 | 19550.875 | 27298.5938 | 20154 |
| **CCL22/MDC** | 21533.1563 | 21283.4375 | 16209.7813 | 16471.7813 | 21408.2969 | 16340.7813 |
| **IL-12 p40** | 7747.46875 | 7143.23438 | 3785.29688 | 3506.59375 | 7445.35156 | 3645.94531 |
| **CCL6/C10** | 22142.3125 | 25314.8125 | 19785.9375 | 20521.0625 | 23728.5625 | 20153.5 |
| **GDF-15** | 7747.75 | 7844.46875 | 4753.90625 | 5122.4375 | 7796.10938 | 4938.17188 |
| **CCL3/CCL4/MIP-1α/β** | 8807.60938 | 8302.98438 | 5892.71875 | 6028.96875 | 8555.29688 | 5960.84375 |
| **GM-CSF** | 3016.4375 | 2781.67188 | 452.140625 | 583.078125 | 2899.05469 | 517.609375 |
| **CCL2/JE/MCP-1** | 5829.8125 | 5609.5625 | 3716.07813 | 3405.71875 | 5719.6875 | 3560.89844 |
| **IL-3** | 11279.6094 | 11558.9844 | 9349.29688 | 9402.26563 | 11419.2969 | 9375.78125 |
| **HGF** | 7582.09375 | 8442.64063 | 6004.59375 | 6469.23438 | 8012.36719 | 6236.91406 |
| **Serpin E1/PAI-1** | 2830.09375 | 3016.6875 | 1100.84375 | 1262.35938 | 2923.39063 | 1181.60156 |
| **Flt-3 Ligand** | 10172.3125 | 10341.5938 | 9108.14063 | 8864.98438 | 10256.9531 | 8986.5625 |
| **IL-6** | 6429.07813 | 6663.15625 | 5407.42188 | 5406.09375 | 6546.11719 | 5406.75781 |
| **IL-23** | 3469.73438 | 3513.54688 | 2491.25 | 2313.23438 | 3491.64063 | 2402.24219 |
| **CXCL1/KC** | 1800.54688 | 1751.20313 | 650.46875 | 742.25 | 1775.875 | 696.359375 |
| **M-CSF** | 1642.96875 | 1734.25 | 564.484375 | 746.796875 | 1688.60938 | 655.640625 |
| **IL-1ra/IL-1F3** | 1875.5 | 2109.76563 | 977.65625 | 1196.26563 | 1992.63281 | 1086.96094 |
| **Angiopoietin-1** | 1059.14063 | 1119.59375 | 297.9375 | 200.96875 | 1089.36719 | 249.453125 |
| **TNF-α** | 6070.28125 | 5855.17188 | 5158.40625 | 5206.15625 | 5962.72656 | 5182.28125 |
| **CCL12/MCP-5** | 2654.46875 | 2466.46875 | 1731.125 | 1970.625 | 2560.46875 | 1850.875 |

**TABLE S4**

| CALM-AF10 | | | MLL-AF10 | | |
|---|---|---|---|---|---|
| Majority protein IDs | Gene names | Unique peptides | Majority protein IDs | Gene names | Unique peptides |
| P52332 | Jak1 | 52 | P52332 | Jak1 | 49 |
| P20918 | Plg | 37 | Q9JHJ0 | Tmod3 | 14 |
| Q9JHJ0 | Tmod3 | 21 | Q91VJ4 | Stk38 | 24 |
| P13020 | Gsn | 32 | Q9WTI7 | Myo1c | 40 |
| Q91VJ4 | Stk38 | 24 | P20918 | Plg | 33 |
| Q91WK0 | Lrrfip2 | 17 | Q99104 | Myo5a | 36 |
| Q920Q8 | Ivns1abp | 19 | Q8BJS4 | Sun2 | 26 |
| Q3THE2 | Myl12b | 10 | Q8CCF0 | Prpf31 | 20 |
| Q922P9 | Glyr1 | 13 | P61161 | Actr2 | 18 |
| Q9CVB6 | Arpc2 | 21 | Q920Q8 | Ivns1abp | 25 |
| P61161 | Actr2 | 16 | P09103 | P4hb | 22 |
| A2AKX3 | Setx | 13 | Q9WUM4 | Coro1c | 16 |
| F7BJB9 | Morc3 | 17 | Q922P9 | Glyr1 | 15 |
| Q61838 | A2m | 21 | Q6VGS5 | Ccdc88c | 23 |
| P07607 | Tyms | 11 | Q9CVB6 | Arpc2 | 15 |
| Q9JM76 | Arpc3 | 10 | P07607 | Tyms | 14 |
| P60710 | Actb | 1 | P16381;Q62 | D1Pas1;Ddx | 3 |
| Q91VN6 | Ddx41 | 10 | F7BJB9 | Morc3 | 14 |
| Q9CQU0 | Txndc12 | 8 | Q9JJ28 | Flii | 14 |
| P68134;P68 | Acta1;Actc1; | 4 | Q9JM76 | Arpc3 | 10 |
| Q8BGD9 | Eif4b | 6 | Q61838 | A2m | 18 |
| P47963 | Rpl13 | 7 | O89053 | Coro1a | 14 |
| Q61233 | Lcp1 | 12 | Q61233 | Lcp1 | 18 |
| Q61189 | Clns1a | 4 | Q8CFE4 | Scyl2 | 15 |
| Q922Q2 | Riok1 | 7 | Q9D0K2 | Oxct1 | 10 |
| Q9ERL7 | Gmfg | 7 | Q921I1 | Tf | 10 |
| Q3THS6 | Mat2a | 7 | Q80WE4 | Kif20b | 14 |
| Q9D154 | Serpinb1a | 14 | P46471 | Psmc2 | 12 |
| P59999 | Arpc4 | 9 | P63268;P68 | Actg2;Acta1 | 3 |
| P14106 | C1qb | 6 | P18760 | Cfl1 | 7 |
| P84089 | Erh | 4 | P59999 | Arpc4 | 10 |
| P21855 | Cd72 | 6 | P62918 | Rpl8 | 9 |
| Q8VC57 | Kctd5 | 6 | P17426 | Ap2a1 | 8 |
| Q921I1 | Tf | 13 | Q6P5D3 | Dhx57 | 11 |
| P14115 | Rpl27a | 4 | Q9ERL7 | Gmfg | 10 |
| Q91X72 | Hpx | 11 | P36993 | Ppm1b | 11 |
| P84104 | Srsf3 | 6 | O88559 | Men1 | 10 |
| P11440 | Cdk1 | 11 | O88685 | Psmc3 | 10 |

| | | | | | |
|---|---|---|---|---|---|
| P18760 | Cfl1 | 5 | Q91VN6 | Ddx41 | 12 |
| P01029 | C4b | 7 | P35564 | Canx | 8 |
| Q6VGS5 | Ccdc88c | 10 | Q3TRM8 | Hk3 | 10 |
| Q9CPW4 | Arpc5 | 5 | Q9Z2B5 | Eif2ak3 | 11 |
| P70266 | Pfkfb1 | 1 | P47757 | Capzb | 12 |
| Q8BTI8 | Srrm2 | 6 | P17751 | Tpi1 | 11 |
| P84091 | Ap2m1 | 7 | Q810B6 | Ankfy1 | 11 |
| P21107;CON | Tpm3;Tpm | 3 | P35278 | Rab5c | 5 |
| Q569Z6 | Thrap3 | 4 | Q8BG32 | Psmd11 | 11 |
| Q8C2Q3 | Rbm14 | 6 | P80317 | Cct6a | 7 |
| Q8CDN6 | Txnl1 | 6 | Q9ESK9 | Rb1cc1 | 11 |
| Q9D0K2 | Oxct1 | 5 | Q8CGY8 | Ogt | 12 |
| Q60749 | Khdrbs1 | 5 | P50516 | Atp6v1a | 10 |
| Q9Z2B5 | Eif2ak3 | 9 | Q9CQU0 | Txndc12 | 5 |
| P42227 | Stat3 | 8 | Q9CPW4 | Arpc5 | 7 |
| Q02105 | C1qc | 3 | P97496 | Smarcc1 | 7 |
| P84099 | Rpl19 | 7 | O54734 | Ddost | 8 |
| O70161 | Pip5k1c | 6 | P19096 | Fasn | 10 |
| Q7M6Y3 | Picalm | 3 | Q8VC57 | Kctd5 | 7 |
| Q9JMA1 | Usp14 | 3 | Q8BHD7 | Ptbp3 | 5 |
| Q8C0P5 | Coro2a | 6 | P11440 | Cdk1 | 7 |
| Q9JM93 | Arl6ip4 | 6 | Q8C006 | Trim35 | 6 |
| Q9D1R9 | Rpl34 | 5 | Q9DBG3 | Ap2b1 | 7 |
| Q8CFE4 | Scyl2 | 12 | Q9D8W5 | Psmd12 | 10 |
| P51675 | Ccr1 | 4 | P52480 | Pkm | 8 |
| Q80WE4 | Kif20b | 7 | P58252 | Eef2 | 11 |
| P43277 | Hist1h1d | 3 | Q9QXS1 | Plec | 9 |
| Q8CH18 | Ccar1 | 7 | P80316 | Cct5 | 10 |
| P35980 | Rpl18 | 6 | P01029 | C4b | 6 |
| P29699 | Ahsg | 7 | P62204;Q9 | Calm1;Calm | 6 |
| P67984 | Rpl22 | 4 | CON__P02769 | | 6 |
| Q8K019 | Bclaf1 | 6 | Q3U9G9 | Lbr | 7 |
| P01027 | C3 | 8 | P61979 | Hnrnpk | 7 |
| Q8K1I7 | Wipf1 | 7 | Q99JI4 | Psmd6 | 9 |
| P35550 | Fbl | 4 | P98086 | C1qa | 4 |
| Q7TSE6 | Stk38l | 4 | Q9D7G0 | Prps1 | 5 |
| Q8VI63 | Mob2 | 2 | Q9R190 | Mta2 | 8 |
| P17751 | Tpi1 | 4 | Q9D0M1 | Prpsap1 | 4 |
| O54879 | Hmgb3 | 4 | Q60932 | Vdac1 | 8 |
| P29416 | Hexa | 5 | Q8VIJ6 | Sfpq | 9 |
| P28665 | Mug1 | 3 | O35593 | Psmd14 | 4 |
| P62267 | Rps23 | 4 | P62880 | Gnb2 | 5 |
| Q99JR8 | Smarcd2 | 5 | P11835 | Itgb2 | 8 |
| P35700 | Prdx1 | 5 | Q9QZE5 | Copg1 | 8 |
| Q8VEM8 | Slc25a3 | 3 | P26516 | Psmd7 | 6 |

| | | | | | |
|---|---|---|---|---|---|
| P62717 | Rpl18a | 6 | O88587 | Comt | 5 |
| Q8R326 | Pspc1 | 6 | P47738 | Aldh2 | 6 |
| Q8C006 | Trim35 | 4 | P12970 | Rpl7a | 7 |
| Q8C1B7 | 11-Sep | 5 | P43277 | Hist1h1d | 3 |
| Q99104 | Myo5a | 28 | P21107;CON | Tpm3;Tpm | 6 |
| P36993 | Ppm1b | 5 | P99024 | Tubb5 | 2 |
| P06151 | Ldha | 6 | P14685 | Psmd3 | 8 |
| P05977;P09 | Myl1;Myl3 | 1 | O88342 | Wdr1 | 7 |
| P98086 | C1qa | 5 | P35700 | Prdx1 | 8 |
| P01872 | Ighm | 9 | Q9R1C7 | Prpf40a | 9 |
| O35226 | Psmd4 | 5 | Q60972 | Rbbp4 | 3 |
| Q52KI8 | Srrm1 | 3 | O88351 | Ikbkb | 7 |
| Q7TMY7 | Ipo8 | 4 | Q9DB77 | Uqcrc2 | 7 |
| Q61142 | Spin1 | 2 | O88398 | Avil | 9 |
| Q61646 | Hp | 4 | P97855 | G3bp1 | 6 |
| P08551 | Nefl | 1 | Q99LB2 | Dhrs4 | 7 |
| Q9CYI4 | Luc7l | 4 | Q9R233 | Tapbp | 8 |
| Q8BGS2 | Bola2 | 2 | P26041 | Msn | 7 |
| Q91YR7 | Prpf6 | 6 | Q9WUA3 | Pfkp | 8 |
| Q9QXS1 | Plec | 15 | P80314 | Cct2 | 9 |
| Q9D7S7 | Rpl22l1 | 2 | Q99JB2 | Stoml2 | 5 |
| P61358 | Rpl27 | 2 | Q9JKR6 | Hyou1 | 6 |
| A2AR02 | Ppig | 4 | P14115 | Rpl27a | 4 |
| Q8BL97 | Srsf7 | 4 | Q6DTY7 | Pfkfb4 | 5 |
| Q80WJ7 | Mtdh | 6 | A6X919 | Dpy19l1 | 6 |
| P18528 | | 1 | Q8BGD9 | Eif4b | 4 |
| Q3TKT4 | Smarca4 | 8 | A2AR02 | Ppig | 4 |
| P62900 | Rpl31 | 1 | Q9R0P5 | Dstn | 5 |
| Q9CQS8 | Sec61b | 2 | Q921F2 | Tardbp | 7 |
| Q62425 | Ndufa4 | 5 | P84104 | Srsf3 | 5 |
| Q9D883 | U2af1 | 3 | P47754 | Capza2 | 5 |
| Q9JMG1 | Edf1 | 2 | Q8BGS2 | Bola2 | 6 |
| P14069 | S100a6 | 1 | P60335 | Pcbp1 | 5 |
| Q9JJI8 | Rpl38 | 1 | Q8JZQ9 | Eif3b | 7 |
| Q99JF8 | Psip1 | 4 | Q91X72 | Hpx | 6 |
| Q8K310 | Matr3 | 4 | Q9QZD9 | Eif3i | 6 |
| Q99M87 | Dnaja3 | 6 | P28665 | Mug1 | 6 |
| P62307 | Snrpf | 2 | Q8BH95 | Echs1 | 4 |
| E9Q3L2 | Pi4ka | 6 | F6ZDS4 | Tpr | 9 |
| Q9CWG9 | Bloc1s2 | 3 | P54775 | Psmc4 | 5 |
| Q8BH43 | Wasf2 | 3 | P62196 | Psmc5 | 6 |
| O55128 | Sap18 | 1 | Q91YR7 | Prpf6 | 10 |
| P97430 | Slpi | 4 | P63037 | Dnaja1 | 6 |
| Q61581 | Igfbp7 | 4 | P70168 | Kpnb1 | 7 |
| Q63870 | Col7a1 | 2 | P70335 | Rock1 | 7 |

| | | | | | |
|---|---|---|---|---|---|
| Q9D0M1 | Prpsap1 | 5 | Q3TCN2 | Plbd2 | 6 |
| Q60668 | Hnrnpd | 7 | Q8BH73 | Qpctl | 5 |
| Q8R574 | Prpsap2 | 3 | Q5XJY5 | Arcn1 | 6 |
| P70335 | Rock1 | 2 | P24547 | Impdh2 | 6 |
| Q80UG5 | 9-Sep | 4 | Q8BFZ9 | Erlin2 | 4 |
| P42208 | 2-Sep | 4 | E9Q3L2 | Pi4ka | 7 |
| Q9CRB9 | Chchd3 | 2 | Q9QXX4 | Slc25a13 | 6 |
| Q9JKB3 | Ybx3 | 3 | Q6URW6 | Myh14 | 2 |
| Q8VH51 | Rbm39 | 3 | Q8VI63 | Mob2 | 4 |
| O89053 | Coro1a | 12 | Q9JM93 | Arl6ip4 | 4 |
| Q9WUM3 | Coro1b | 7 | P62751 | Rpl23a | 5 |
| A6X919 | Dpy19l1 | 4 | P43275 | Hist1h1a | 4 |
| Q9D898 | Arpc5l | 4 | Q61142 | Spin1 | 5 |
| P43274 | Hist1h1e | 4 | Q8BTI8 | Srrm2 | 6 |
| CON__P02769 | | 3 | P42227 | Stat3 | 7 |
| Q9Z183 | Padi4 | 11 | Q6ZWV7 | Rpl35 | 4 |
| Q04750 | Top1 | 12 | Q8CI43 | Myl6b | 4 |
| Q7TSG5 | Sh3d21 | 1 | Q9CQQ7 | Atp5f1 | 4 |
| O35691 | Pnn | 4 | P21855 | Cd72 | 6 |
| Q80UW3 | Rinl | 3 | Q9CY27 | Tecr | 4 |
| Q3UM45 | Ppp1r7 | 6 | Q9Z0H4 | Celf2 | 6 |
| P61255 | Rpl26 | 3 | Q8VCC1 | Hpgd | 6 |
| Q3TCN2 | Plbd2 | 3 | P62301 | Rps13 | 5 |
| Q8JZU2 | Slc25a1 | 6 | Q9CQE8 | | 5 |
| O35737;P70 | Hnrnph1;H | 4 | O89079 | Cope | 3 |
| P63087 | Ppp1cc | 2 | P80315 | Cct4 | 8 |
| Q9D3D9 | Atp5d | 3 | Q61646 | Hp | 5 |
| P01636 | | 5 | P80318 | Cct3 | 7 |
| P07091 | S100a4 | 2 | P16045 | Lgals1 | 4 |
| P47753 | Capza1 | 4 | P05555 | Itgam | 8 |
| P47757 | Capzb | 5 | P42932 | Cct8 | 5 |
| Q9JKV1 | Adrm1 | 4 | P62334 | Psmc6 | 5 |
| Q9ESK9 | Rb1cc1 | 2 | P62858 | Rps28 | 5 |
| P62996 | Tra2b | 4 | Q8R326 | Pspc1 | 5 |
| Q922V4 | Plrg1 | 4 | Q61990;P57 | Pcbp2;Pcbp | 3 |
| Q7M6Z4 | Kif27 | 1 | Q9WUM5 | Suclg1 | 4 |
| P30681 | Hmgb2 | 5 | P01027 | C3 | 5 |
| Q63955 | Pou4f3 | 1 | Q78ZA7 | Nap1l4 | 3 |
| Q9CX56 | Psmd8 | 4 | Q9R0E1 | Plod3 | 5 |
| Q6ZWU9 | Rps27 | 2 | Q924C1 | Xpo5 | 6 |
| Q9JI48 | Plac8 | 1 | Q9QYB5 | Add3 | 3 |
| E9Q634 | Myo1e | 2 | Q64518 | Atp2a3 | 5 |
| P62751 | Rpl23a | 4 | A2AKX3 | Setx | 4 |
| P18527 | | 2 | P62315 | Snrpd1 | 3 |
| P61294;P35 | Rab6b;Rab6 | 1 | P51675 | Ccr1 | 4 |

| | | | | | |
|---|---|---|---|---|---|
| P62830 | Rpl23 | 6 | P14106 | C1qb | 4 |
| P60335 | Pcbp1 | 3 | P62855 | Rps26 | 3 |
| Q62095 | Ddx3y | 1 | P61620;Q9 | Sec61a1;Sec | 3 |
| P62827;Q6 | Ran;Rasl2-9 | 4 | Q9CQW9 | Ifitm3 | 3 |
| O54941 | Smarce1 | 4 | Q62425 | Ndufa4 | 4 |
| P03977 | | 1 | Q80UW3 | Rinl | 4 |
| Q8CH25 | Sltm | 2 | O09106 | Hdac1 | 3 |
| Q99M28 | Rnps1 | 2 | Q3THS6 | Mat2a | 5 |
| P84244;P8 | H3f3a;Hist | 3 | Q61207 | Psap | 6 |
| Q9CQJ6 | Denr | 3 | Q9CQ62 | Decr1 | 5 |
| P63158 | Hmgb1 | 1 | P47753 | Capza1 | 4 |
| Q8K010 | Oplah | 1 | Q9D1G1;P6 | Rab1b;Rab1 | 4 |
| Q9DBM1 | Gpatch1 | 1 | Q9D051 | Pdhb | 5 |
| Q9DCF9 | Ssr3 | 1 | P19783 | Cox4i1 | 5 |
| Q62186 | Ssr4 | 4 | Q8C0P5 | Coro2a | 5 |
| Q9CQY5 | Magt1 | 4 | Q8CH18 | Ccar1 | 5 |
| Q9QUI0;Q6 | Rhoa;Rhoc | 3 | Q6PDM2 | Srsf1 | 6 |
| Q921L3 | Tmco1 | 2 | Q8BK63 | Csnk1a1 | 3 |
| Q8BK63 | Csnk1a1 | 5 | O35691 | Pnn | 4 |
| P59764 | Dock4 | 3 | P62192 | Psmc1 | 4 |
| Q8BH95 | Echs1 | 2 | Q8K297 | Colgalt1 | 4 |
| P62843 | Rps15 | 2 | Q9Z183 | Padi4 | 6 |
| Q9QY73 | Tmem59 | 2 | Q6ZQ38 | Cand1 | 5 |
| P62892 | Rpl39 | 1 | B1AZI6 | Thoc2 | 7 |
| Q9CWZ3 | Rbm8a | 1 | Q3TBT3 | Tmem173 | 4 |
| Q3TWW8;Q | Srsf6;Srsf5 | 1 | P35486 | Pdha1 | 3 |
| Q6P5D3 | Dhx57 | 2 | Q8R1Q8 | Dync1li1 | 4 |
| Q9DCE5 | Pak1ip1 | 5 | Q9WUK4 | Rfc2 | 3 |
| P53986 | Slc16a1 | 2 | Q920E5 | Fdps | 3 |
| P62806 | Hist1h4a | 11 | Q62465 | Vat1 | 5 |
| Q8CJ40 | Crocc | 2 | Q8JZM7 | Cdc73 | 4 |
| Q9D0M5 | Dynll2 | 2 | Q8BMF4 | Dlat | 3 |
| Q7TSC1 | Prrc2a | 2 | Q64310 | Surf4 | 2 |
| P62849 | Rps24 | 4 | Q9DB27 | Mcts1 | 5 |
| P31996 | Cd68 | 3 | Q9CYG7 | Tomm34 | 4 |
| P35486 | Pdha1 | 4 | Q921G6 | Lrch4 | 4 |
| Q924C1 | Xpo5 | 4 | P62874 | Gnb1 | 2 |
| Q9R0P5 | Dstn | 1 | P05977;P09 | Myl1;Myl3 | 2 |
| Q4VA53 | Pds5b | 7 | Q9CZM2 | Rpl15 | 4 |
| P62309 | Snrpg | 2 | P62317 | Snrpd2 | 5 |
| Q80W93 | Hydin | 1 | P0CW03 | Ly6c2 | 4 |
| P17427 | Ap2a2 | 1 | Q9D898 | Arpc5l | 3 |
| Q6PGB6 | Naa50 | 1 | P43276 | Hist1h1b | 4 |
| P04187 | Gzmb | 6 | Q52KI8 | Srrm1 | 3 |
| Q91VR8 | Brk1 | 5 | Q61189 | Clns1a | 2 |

| | | | | | |
|---|---|---|---|---|---|
| Q9R0Q3 | Tmed2 | 1 | Q9JJU8 | Sh3bgrl | 4 |
| Q9CS42 | Prps2 | 1 | Q9QYJ0 | Dnaja2 | 6 |
| Q5SUF2 | Luc7l3 | 2 | Q8VEM8 | Slc25a3 | 3 |
| Q60973 | Rbbp7 | 3 | P19536 | Cox5b | 3 |
| P97450 | Atp5j | 2 | O35226 | Psmd4 | 4 |
| Q9WTX5 | Skp1 | 5 | O70251 | Eef1b | 4 |
| Q99LT0 | Dpy30 | 2 | Q7TSE6 | Stk38l | 3 |
| Q9D1M7 | Fkbp11 | 4 | O70503 | Hsd17b12 | 4 |
| Q61096 | Prtn3 | 2 | Q9DB20 | Atp5o | 6 |
| Q60692 | Psmb6 | 3 | Q9DCX2 | Atp5h | 4 |
| Q9CQQ7 | Atp5f1 | 3 | Q9D883 | U2af1 | 3 |
| Q9Z2X2 | Psmd10 | 1 | Q91VW3 | Sh3bgrl3 | 2 |
| P35278 | Rab5c | 5 | P84089 | Erh | 4 |
| P55258;Q9I | Rab8a;Rab1 | 1 | O55142 | Rpl35a | 3 |
| Q8K297 | Colgalt1 | 6 | O54879 | Hmgb3 | 3 |
| P52480 | Pkm | 5 | P62849 | Rps24 | 3 |
| Q9WUM5 | Suclg1 | 3 | Q9CPQ8 | Atp5l | 2 |
| O09117 | Sypl1 | 1 | Q9CPR4 | Rpl17 | 5 |
| Q8N7N5 | Dcaf8 | 6 | Q91WK2 | Eif3h | 6 |
| P18525 | | 1 | Q99JR1 | Sfxn1 | 4 |
| P62878 | Rbx1 | 1 | Q9DCJ5 | Ndufa8 | 4 |
| O08788 | Dctn1 | 6 | Q8BU14 | Sec62 | 3 |
| Q9CQN1 | Trap1 | 2 | P68368 | Tuba4a | 4 |
| Q9CPQ1 | Cox6c | 3 | O08583 | Alyref | 3 |
| Q8BKT7 | Thoc5 | 4 | P46978 | Stt3a | 3 |
| Q9D358 | Acp1 | 1 | Q9CQN1 | Trap1 | 4 |
| P15379 | Cd44 | 2 | Q60931 | Vdac3 | 3 |
| P33174 | Kif4 | 9 | B2RY56 | Rbm25 | 1 |
| Q9CRD2 | Emc2 | 3 | P61965 | Wdr5 | 5 |
| Q8R035 | Ict1 | 3 | Q9CX56 | Psmd8 | 3 |
| P24668 | M6pr | 1 | Q8K310 | Matr3 | 3 |
| P61222 | Abce1 | 1 | P01872 | Ighm | 5 |
| Q8VDT9 | Mrpl50 | 5 | P12382 | Pfkl | 5 |
| Q99M15 | Pstpip2 | 5 | Q61543 | Glg1 | 6 |
| Q61103 | Dpf2 | 3 | Q9DCH4 | Eif3f | 7 |
| Q5SFM8;Q( | Rbm27;Rbr | 2 | Q3TEA8 | Hp1bp3 | 2 |
| Q80Y86 | Mapk15 | 1 | Q99PL5 | Rrbp1 | 5 |
| Q8BGF7 | Pan2 | 2 | P00405 | Mtco2 | 3 |
| Q6R5N8 | Tlr13 | 1 | Q9Z0P5 | Twf2 | 3 |
| Q8BTV2 | Cpsf7 | 4 | Q64331 | Myo6 | 4 |
| Q9D287 | Bcas2 | 4 | O88487 | Dync1i2 | 3 |
| O88342 | Wdr1 | 8 | Q9EP72 | Emc7 | 4 |
| P01642 | Gm10881 | 2 | P26638 | Sars | 6 |
| Q08288 | Lyar | 2 | Q8CE96 | Trmt6 | 5 |
| Q6ZWV7 | Rpl35 | 6 | Q6GQT9 | Nomo1 | 5 |

| | | | | | |
|---|---|---|---|---|---|
| P01680 | | 1 | Q6PDG5 | Smarcc2 | 3 |
| P02088;CO | Hbb-b1;Hb | 2 | Q99KQ4 | Nampt | 5 |
| P11031 | Sub1 | 2 | Q9DC69 | Ndufa9 | 3 |
| O54988 | Slk | 5 | Q61881 | Mcm7 | 5 |
| Q9QYC0 | Add1 | 2 | Q8R323 | Rfc3 | 6 |
| Q8BH73 | Qpctl | 4 | O88796 | Rpp30 | 4 |
| O55142 | Rpl35a | 4 | P29416 | Hexa | 5 |
| Q00896;Po | Serpina1c;S | 2 | Q8BWZ3 | Naa25 | 3 |
| Q9JJU8 | Sh3bgrl | 4 | Q8BGZ4 | Cdc23 | 5 |
| Q60931 | Vdac3 | 6 | P52293 | Kpna2 | 4 |
| P51125 | Cast | 1 | Q9D0M5 | Dynll2 | 1 |
| P01660 | | 1 | Q02105 | C1qc | 3 |
| Q8VCW4 | Unc93b1 | 2 | Q64152 | Btf3 | 2 |
| Q9QX47 | Son | 3 | Q7TNV0 | Dek | 3 |
| Q8BFZ9 | Erlin2 | 5 | P84099 | Rpl19 | 4 |
| P10404;P11 | Fv4 | 5 | P56391 | Cox6b1 | 3 |
| Q8BP67 | Rpl24 | 3 | Q9WVJ2 | Psmd13 | 3 |
| Q6ZQ58 | Larp1 | 5 | P84091 | Ap2m1 | 3 |
| P01723;P01727 | | 2 | Q80SY5 | Prpf38b | 3 |
| Q8BFZ3 | Actbl2 | 1 | Q8BH43 | Wasf2 | 5 |
| Q80UK7 | Sass6 | 3 | P05532 | Kit | 2 |
| Q5SS90 | Gm11992 | 1 | O70551;Q9Z | Srpk1;Srpk3 | 3 |
| P97814 | Pstpip1 | 2 | Q06185 | Atp5i | 3 |
| Q99LX0 | Park7 | 1 | Q99M28 | Rnps1 | 3 |
| Q9QVN7 | Tcea2 | 1 | Q61096 | Prtn3 | 5 |
| Q8VHK9 | Dhx36 | 5 | Q91VR2 | Atp5c1 | 4 |
| P56135 | Atp5j2 | 2 | P12787 | Cox5a | 4 |
| P68369;P05 | Tuba1a;Tub | 1 | CON__P01966 | | 2 |
| P04944;P04941;P04946 | | 1 | Q9WTX5 | Skp1 | 3 |
| B2RY56 | Rbm25 | 2 | Q9CQJ6 | Denr | 5 |
| O55201 | Supt5h | 3 | Q99JI6;P62 | Rap1b;Rap1 | 3 |
| Q80SY5 | Prpf38b | 3 | Q9CYN2 | Spcs2 | 4 |
| Q9QWT9 | Kifc1 | 8 | Q9D0F3 | Lman1 | 4 |
| P83882 | Rpl36a | 1 | Q62351 | Tfrc | 3 |
| P01901 | H2-K1 | 4 | O55135 | Eif6 | 4 |
| Q9CXL3 | | 3 | P97450 | Atp5j | 2 |
| Q9Z0R9 | Fads2 | 3 | Q60668 | Hnrnpd | 3 |
| P48771 | Cox7a2 | 2 | Q9D0F6 | Rfc5 | 4 |
| Q9DA19 | Cir1 | 1 | Q91V41 | Rab14 | 2 |
| P47964 | Rpl36 | 3 | P48962 | Slc25a4 | 4 |
| P10639 | Txn | 2 | Q6PGB6 | Naa50 | 3 |
| Q8K396 | Mnd1 | 1 | O54941 | Smarce1 | 5 |
| Q99KC8 | Vwa5a | 5 | Q9JIX8 | Acin1 | 2 |
| Q8CDE2 | Ccin | 2 | Q6ZQA0 | Nbeal2 | 4 |
| O35723 | Dnajb3 | 1 | P70372 | Elavl1 | 3 |

| | | | | | |
|---|---|---|---|---|---|
| P01750 | | 1 | Q9QWT9 | Kifc1 | 6 |
| P49586;Q8 | Pcyt1a;Pcyt1 | 2 | P06330;P01757;P01756 | | 3 |
| A2ASS6 | Ttn | 3 | Q5SYD0 | Myo1d | 3 |
| Q8BTZ4 | Anapc5 | 3 | P56135 | Atp5j2 | 2 |
| CON__ENSEMBL:ENS | | 4 | Q78IK2 | Usmg5 | 3 |
| Q61464 | Znf638 | 2 | Q80UM3 | Naa15 | 4 |
| P01674 | | 1 | P37040 | Por | 5 |
| P84750 | | 3 | P60766 | Cdc42 | 2 |
| Q9CZ44 | Nsfl1c | 2 | Q9Z1D1 | Eif3g | 2 |
| Q8BH59 | Slc25a12 | 4 | P07901 | Hsp90aa1 | 6 |
| Q00724 | Rbp4 | 1 | P24668 | M6pr | 2 |
| P01844;P01 | Iglc2;Iglc3 | 3 | Q8BFY9 | Tnpo1 | 3 |
| P01634 | | 1 | A2AIV8 | Card9 | 3 |
| Q9CWX9 | Ddx47 | 5 | Q6PE01 | Snrnp40 | 4 |
| Q8BX10 | Pgam5 | 3 | Q3UJB9 | Edc4 | 3 |
| Q3THK3 | Gtf2f1 | 2 | P17182 | Eno1 | 3 |
| Q3TEA8 | Hp1bp3 | 2 | Q1HFZ0 | Nsun2 | 4 |
| Q8VE37 | Rcc1 | 5 | Q9EPL8 | Ipo7 | 2 |
| Q99J72 | Apobec3 | 4 | Q922J9 | Far1 | 3 |
| Q8VCH8 | Ubxn4 | 1 | Q3TDQ1 | Stt3b | 3 |
| A2BDX3 | Mocs3 | 2 | Q8BUR9 | Mzt1 | 2 |
| P06329 | | 1 | P59235 | Nup43 | 3 |
| Q9R0Q6 | Arpc1a | 1 | Q8VE80 | Thoc3 | 2 |
| Q6URW6 | Myh14 | 2 | P62492;P4 | Rab11a;Rab | 4 |
| B1AY13 | Usp24 | 2 | A2AJT4 | Pnisr | 2 |
| Q8VDD9 | Phip | 2 | A2AGT5 | Ckap5 | 6 |
| Q9DCD2 | Xab2 | 2 | Q9Z2V5 | Hdac6 | 2 |
| Q9DC48 | Cdc40 | 3 | P70333;O3 | Hnrnph2;H | 2 |
| Q8BSY0 | Asph | 9 | Q64378 | Fkbp5 | 2 |
| P61620;Q9 | Sec61a1;Sec | 3 | Q6ZPE2 | Sbf1 | 3 |
| Q9D6Z1 | Nop56 | 4 | Q8BKT7 | Thoc5 | 4 |
| Q8R0X7 | Sgpl1 | 4 | Q9DCT2 | Ndufs3 | 3 |
| Q8BHN1 | Txlng | 3 | Q9CZW5 | Tomm70a | 3 |
| P03930 | Mtatp8 | 1 | A2BH40 | Arid1a | 3 |
| Q9CPX7 | Mrps16 | 4 | A2BE28 | Las1l | 3 |
| P26369 | U2af2 | 5 | Q99M31 | Hspa14 | 4 |
| P62305 | Snrpe | 2 | P53994 | Rab2a | 4 |
| Q7TMY8 | Huwe1 | 3 | P97376 | Frg1 | 4 |
| Q923D4 | Sf3b5 | 3 | P97377;Q8 | Cdk2;Cdk3 | 2 |
| Q3TUH1 | Tamm41 | 3 | Q9D1M7 | Fkbp11 | 2 |
| Q61990;P57 | Pcbp2;Pcbp | 3 | P08508 | Fcgr3 | 2 |
| P25976 | Ubtf | 2 | Q9DBM1 | Gpatch1 | 2 |
| Q80X41 | Vrk1 | 8 | Q8N7N5 | Dcaf8 | 4 |
| O09167 | Rpl21 | 3 | Q9DC28;Q | Csnk1d;Csn | 3 |
| Q8VCM8 | Ncln | 4 | Q60875 | Arhgef2 | 4 |

| | | | | | |
|---|---|---|---|---|---|
| P51807 | Dynlt1 | 2 | Q9CY50 | Ssr1 | 3 |
| P60766 | Cdc42 | 4 | CON__Q3SX09;CON__ | | 3 |
| O08807 | Prdx4 | 2 | B9EJ86 | Osbpl8 | 2 |
| Q8BLD6 | Ankrd55 | 1 | Q9D7N9 | Apmap | 2 |
| Q9CQ75 | Ndufa2 | 2 | P49717 | Mcm4 | 2 |
| Q9ES97 | Rtn3 | 2 | Q61823 | Pdcd4 | 3 |
| P61514 | Rpl37a | 1 | O08756 | Hsd17b10 | 4 |
| Q61735 | Cd47 | 1 | CON__P35908 | | 3 |
| Q9CQL1;P6 | Magohb;Ma | 2 | Q9CR68 | Uqcrfs1 | 1 |
| P37913 | Lig1 | 3 | Q69ZR2 | Hectd1 | 3 |
| P50516 | Atp6v1a | 4 | P16254 | Srp14 | 4 |
| Q9ESD6 | Cmtm7 | 1 | Q8BVE3 | Atp6v1h | 1 |
| Q9DCA2 | Mrps11 | 3 | Q8BFQ8 | Pddc1 | 2 |
| Q9D0T1 | Nhp2l1 | 7 | Q9QZD8 | Slc25a10 | 1 |
| Q9R1C7 | Prpf40a | 5 | Q9CXY6 | Ilf2 | 6 |
| Q8VC03 | Eml3 | 3 | P62814 | Atp6v1b2 | 5 |
| P20491 | Fcer1g | 2 | P27870 | Vav1 | 4 |
| P01820;P01819 | | 1 | Q9JI48 | Plac8 | 1 |
| Q9R0Q4 | Morf4l2 | 1 | P62852 | Rps25 | 4 |
| D0QMC3 | Mndal | 3 | P15864 | Hist1h1c | 3 |
| P54116 | Stom | 7 | P43024 | Cox6a1 | 2 |
| Q8VHR5 | Gatad2b | 2 | Q05D44 | Eif5b | 4 |
| A2RTL5 | Rsrc2 | 1 | P62900 | Rpl31 | 1 |
| Q8K4B0 | Mta1 | 3 | P30681 | Hmgb2 | 5 |
| P19973 | Lsp1 | 4 | P31725 | S100a9 | 2 |
| Q8VE97 | Srsf4 | 1 | Q9D3D9 | Atp5d | 2 |
| Q9D0F3 | Lman1 | 3 | Q9CQS8 | Sec61b | 2 |
| Q8BVY0 | Rsl1d1 | 3 | P04187 | Gzmb | 3 |
| P28660 | Nckap1 | 2 | P61804 | Dad1 | 3 |
| P01899;P01 | H2-D1;H2-l | 1 | Q91VR8 | Brk1 | 5 |
| Q9CYL5 | Glipr2 | 1 | Q99JR8 | Smarcd2 | 3 |
| P14719 | Il1rl1 | 1 | P07091 | S100a4 | 1 |
| Q99PG2 | Ogfr | 1 | P97370 | Atp1b3 | 4 |
| Q61136 | Prpf4b | 8 | Q9R1P0 | Psma4 | 3 |
| P28076 | Psmb9 | 2 | Q9CQL1;P6 | Magohb;Ma | 2 |
| Q9JJA4 | Wdr12 | 5 | Q9ES97 | Rtn3 | 2 |
| O35598 | Adam10 | 4 | P35550 | Fbl | 3 |
| G5E870 | Trip12 | 5 | P52503 | Ndufs6 | 2 |
| Q8BM55 | Tmem214 | 5 | Q9QYC0 | Add1 | 2 |
| P01648;P01649 | | 1 | P29699 | Ahsg | 2 |
| P01679;P01678;P01677; | | 2 | Q62186 | Ssr4 | 3 |
| CON__ENSEMBL:ENS | | 5 | Q3UGC7;Q | Eif3j1;Eif3j | 2 |
| P01878 | | 9 | Q9CYI4 | Luc7l | 2 |
| P01666;P01665;P01668 | | 1 | P62806 | Hist1h4a | 4 |
| Q8VE22 | Mrps23 | 7 | Q9JMA1 | Usp14 | 2 |

| | | | | | |
|---|---|---|---|---|---|
| P01630 | | 3 | O55128 | Sap18 | 2 |
| Q9EP72 | Emc7 | 4 | Q9CZU3 | Skiv2l2 | 2 |
| P01747;P01746 | | 3 | Q9CS42 | Prps2 | 2 |
| CON__Streptavidin | | 2 | P28656 | Nap1l1 | 2 |
| P62869 | Tceb2 | 5 | A2BDX3 | Mocs3 | 1 |
| P83940 | Tceb1 | 4 | P40124 | Cap1 | 2 |
| Q9ER88 | Dap3 | 12 | Q9CQZ5 | Ndufa6 | 2 |
| P42669 | Pura | 4 | Q9CXL3 | | 3 |
| Q8K2T8 | Paf1 | 6 | P62743 | Ap2s1 | 2 |
| O35972 | Mrpl23 | 2 | Q8VE37 | Rcc1 | 4 |
| Q9CPP0 | Npm3 | 3 | Q9CWK8 | Snx2 | 2 |
| Q99N96 | Mrpl1 | 8 | P19467 | Muc13 | 3 |
| P61982 | Ywhag | 4 | P54071 | Idh2 | 2 |
| Q05CL8 | Larp7 | 6 | P29477 | Nos2 | 2 |
| Q9D1P0 | Mrpl13 | 5 | O89051 | Itm2b | 3 |
| P27048;P6 | Snrpb;Snrp | 6 | Q9QUI0;Q | Rhoa;Rhoc | 2 |
| P97760 | Polr2c | 7 | P62267 | Rps23 | 3 |
| Q9CQF0 | Mrpl11 | 7 | P68254 | Ywhaq | 2 |
| Q9D7N3 | Mrps9 | 6 | O70325;Q9 | Gpx4 | 2 |
| Q922U1 | Prpf3 | 9 | Q8C1B7 | 11-Sep | 2 |
| P12382 | Pfkl | 6 | Q99JY0 | Hadhb | 4 |
| Q921S7 | Mrpl37 | 7 | P63325 | Rps10 | 3 |
| P16254 | Srp14 | 5 | Q9DBH5 | Lman2 | 4 |
| Q9DBS1 | Tmem43 | 4 | O08917 | Flot1 | 2 |
| Q6ZWM4 | Lsm8 | 3 | P97311 | Mcm6 | 3 |
| Q8CFQ3 | Aqr | 6 | P25976 | Ubtf | 2 |
| Q99N94 | Mrpl9 | 4 | Q8R2Y8 | Ptrh2 | 2 |
| O70310;O7 | Nmt1;Nmt2 | 4 | Q923D4 | Sf3b5 | 3 |
| P83870 | Phf5a | 4 | Q5U4D9 | Thoc6 | 5 |
| Q9CY16 | Mrps28 | 5 | Q9ERI2 | Rab27a | 1 |
| P31266 | Rbpj | 4 | Q3TKT4 | Smarca4 | 3 |
| Q9CY52 | Thg1l | 4 | P06800 | Ptprc | 3 |
| Q8C052 | Map1s | 6 | Q80Y86 | Mapk15 | 1 |
| Q62383 | Supt6h | 11 | Q99JX4 | Eif3m | 2 |
| P97376 | Frg1 | 3 | A2AN08 | Ubr4 | 3 |
| Q14C51 | Ptcd3 | 6 | Q921N6 | Ddx27 | 5 |
| P56873 | Sssca1 | 3 | Q9CRB9 | Chchd3 | 2 |
| Q99JI6;P6 | Rap1b;Rap1 | 4 | P61750 | Arf4 | 2 |
| P01790;P01794;P01791; | | 3 | Q9WUM3 | Coro1b | 2 |
| Q99LD4 | Gps1 | 5 | P97371 | Psme1 | 2 |
| P51670 | Ccl9 | 3 | Q8CJ40 | Crocc | 1 |
| Q7TSI3 | Ppp6r1 | 4 | P14719 | Il1rl1 | 2 |
| Q9CY73 | Mrpl44 | 3 | Q99LX0 | Park7 | 1 |
| Q8K2M0 | Mrpl38 | 5 | Q8K2B3 | Sdha | 4 |
| A1L314 | Mpeg1 | 4 | Q8R1I1 | Uqcr10 | 1 |

| | | | | | |
|---|---|---|---|---|---|
| Q8VDF2 | Uhrf1 | 4 | P10639 | Txn | 1 |
| P49300 | Clec10a | 4 | Q8R5H1;P3 | Usp15;Usp4 | 2 |
| O35343 | Kpna4 | 4 | O35638 | Stag2 | 2 |
| Q8R1F9 | Rpp40 | 3 | Q9Z1Q9 | Vars | 2 |
| Q9Z321 | Top3b | 4 | P97452 | Bop1 | 3 |
| Q6ZQI3 | Mlec | 5 | P83870 | Phf5a | 5 |
| Q9D0M0 | Exosc7 | 4 | Q8C6G8 | Wdr26 | 2 |
| Q5SWD9 | Tsr1 | 7 | Q99M15 | Pstpip2 | 4 |
| Q921N6 | Ddx27 | 8 | CON__P06868 | | 4 |
| Q9CQC7 | Ndufb4 | 4 | Q9JKV1 | Adrm1 | 1 |
| P49722 | Psma2 | 4 | Q8BMA6 | Srp68 | 2 |
| O88545 | Cops6 | 4 | Q8R5F3 | Oard1 | 1 |
| Q9Z120 | Mettl1 | 4 | P63101 | Ywhaz | 2 |
| Q9D6J6 | Ndufv2 | 3 | Q9CPN8 | Igf2bp3 | 3 |
| Q80X85 | Mrps7 | 6 | O70310;O7( | Nmt1;Nmt2 | 4 |
| P97484 | Lilrb3 | 5 | A2A6A1 | Gpatch8 | 2 |
| P29351 | Ptpn6 | 7 | Q9D8V0 | Hm13 | 2 |
| Q8BP48 | Metap1 | 4 | P08003 | Pdia4 | 2 |
| O08585 | Clta | 3 | P61021 | Rab5b | 2 |
| P63085 | Mapk1 | 4 | Q922Q2 | Riok1 | 2 |
| Q9ER72 | Cars | 4 | Q8VHK9 | Dhx36 | 3 |
| Q64337 | Sqstm1 | 4 | Q8C147 | Dock8 | 5 |
| Q91W96 | Anapc4 | 4 | Q8BX70 | Vps13c | 3 |
| Q9WUQ2 | Preb | 4 | P70288 | Hdac2 | 2 |
| Q9QXK7 | Cpsf3 | 2 | O35295 | Purb | 3 |
| O35955 | Psmb10 | 4 | O35130 | Emg1 | 3 |
| P70318 | Tial1 | 1 | Q9Z0R9 | Fads2 | 3 |
| P59328 | Wdhd1 | 5 | Q3TZX8 | Nol9 | 2 |
| Q99LC3 | Ndufa10 | 6 | Q62318 | Trim28 | 2 |
| Q9WU56 | Pus1 | 4 | O09117 | Sypl1 | 1 |
| Q8CCP0 | Nemf | 5 | Q91X78 | Erlin1 | 2 |
| Q9DCL9 | Paics | 8 | Q5SW75 | Ssh2 | 2 |
| Q922D4 | Ppp6r3 | 4 | Q80VD1 | Fam98b | 3 |
| Q8K3W0 | Bre | 4 | Q9CPQ3 | Tomm22 | 2 |
| P35601 | Rfc1 | 6 | Q924T2 | Mrps2 | 3 |
| Q8JZM7 | Cdc73 | 5 | Q91V04 | Tram1 | 2 |
| O35654 | Pold2 | 4 | Q99J56 | Derl1 | 2 |
| Q8R3F9 | Tut1 | 5 | Q9QYA2 | Tomm40 | 1 |
| Q9CZ13 | Uqcrc1 | 6 | Q9DCE5 | Pak1ip1 | 5 |
| Q9JKC8 | Ap3m1 | 4 | Q9Z127 | Slc7a5 | 2 |
| P19467 | Muc13 | 4 | Q9ERN0 | Scamp2 | 1 |
| Q9JHI7 | Exosc9 | 4 | Q8K2Z4 | Ncapd2 | 3 |
| CON__ENSEMBL:ENS | | 6 | P13439 | Umps | 2 |
| P68254 | Ywhaq | 5 | Q8K1B8 | Fermt3 | 6 |
| Q9WVG6 | Carm1 | 3 | Q8JZN5 | Acad9 | 2 |

| | | | | | |
|---|---|---|---|---|---|
| Q9DBT5 | Ampd2 | 4 | Q61102 | Abcb7 | 2 |
| Q9D753 | Exosc8 | 4 | P19973 | Lsp1 | 2 |
| Q80X73 | Pelo | 5 | P53986 | Slc16a1 | 3 |
| Q8R480 | Nup85 | 8 | Q91YT0 | Ndufv1 | 2 |
| Q8VHL1 | Setd7 | 3 | Q3TIX9 | Usp39 | 4 |
| Q8K482 | Emilin2 | 4 | Q64261 | Cdk6 | 2 |
| Q8C2K5 | Rasal3 | 5 | Q6PDI5 | Ecm29 | 2 |
| Q7TQH0 | Atxn2l | 6 | Q99N94 | Mrpl9 | 3 |
| Q62419 | Sh3gl1 | 6 | Q80X73 | Pelo | 4 |
| Q8CGZ0 | Cherp | 4 | Q00896;Q0 | Serpina1c;S | 1 |
| Q8K2B3 | Sdha | 5 | Q8C7X2 | Emc1 | 5 |
| Q91XU0 | Wrnip1 | 5 | P62996 | Tra2b | 1 |
| Q69ZR2 | Hectd1 | 4 | Q921F4 | Hnrnpll | 2 |
| Q8BX70 | Vps13c | 6 | P09055 | Itgb1 | 2 |
| Q8CHW4 | Eif2b5 | 4 | Q09014 | Ncf1 | 2 |
| P07742 | Rrm1 | 3 | Q9JKF7 | Mrpl39 | 4 |
| O70378 | Emc8 | 4 | Q6P5F9 | Xpo1 | 2 |
| Q99JB2 | Stoml2 | 5 | Q9CQD1 | Rab5a | 1 |
| Q6PGG2 | Gmip | 4 | P52431 | Pold1 | 2 |
| Q571I9 | Aldh16a1 | 4 | Q9D1M4 | Eef1e1 | 3 |
| Q62351 | Tfrc | 5 | Q8BYZ1 | Abi3 | 1 |
| Q8BWU5 | Osgep | 3 | Q9ESB3 | Hrg | 2 |
| Q9D338 | Mrpl19 | 4 | P61027 | Rab10 | 1 |
| Q8VBV7 | Cops8 | 3 | Q3TDD9 | Ppp1r21 | 5 |
| A2BE28 | Las1l | 5 | Q6PD26 | Pigs | 1 |
| Q5SSH7 | Zzef1 | 5 | P99028 | Uqcrh | 2 |
| Q9DC69 | Ndufa9 | 4 | Q8K4F6 | Nsun5 | 1 |
| Q6ZQ03 | Fnbp4 | 4 | O70404 | Vamp8 | 1 |
| Q64310 | Surf4 | 2 | P61514 | Rpl37a | 1 |
| Q60520 | Sin3a | 4 | P67984 | Rpl22 | 1 |
| P97770 | Thumpd3 | 4 | P62843 | Rps15 | 3 |
| Q99KN2 | Ciao1 | 4 | Q9D1R9 | Rpl34 | 2 |
| Q8CCJ3 | Ufl1 | 4 | O09167 | Rpl21 | 2 |
| Q9QXE7 | Tbl1x | 2 | P63168 | Dynll1 | 1 |
| P54731 | Faf1 | 3 | P61294;P35 | Rab6b;Rab6 | 1 |
| Q9CPX6 | Atg3 | 3 | P47964 | Rpl36 | 2 |
| Q9JLV5 | Cul3 | 5 | Q9EP89 | Lactb | 2 |
| Q99JT2;Q9 | Stk26;Stk25 | 3 | Q8R010 | Aimp2 | 3 |
| D3YXK2 | Safb | 5 | Q9CWG9 | Bloc1s2 | 2 |
| P62814 | Atp6v1b2 | 3 | Q80W93 | Hydin | 2 |
| CON__P06868 | | 4 | O70405 | Ulk1 | 1 |
| Q60953 | Pml | 3 | P97379 | G3bp2 | 1 |
| P70670;Q6 | Naca | 3 | Q31125 | Slc39a7 | 1 |
| P54729 | Nub1 | 3 | Q62095 | Ddx3y | 1 |
| P18524 | | 0 | Q8C2Q3 | Rbm14 | 2 |

| | | | | | |
|---|---|---|---|---|---|
| P63154 | Crnkl1 | 2 | P97430 | Slpi | 3 |
| Q9D3R6;Q9 | Katnal2;Spa | 1 | O08807 | Prdx4 | 1 |
| P70700 | Polr1b | 4 | Q99J72 | Apobec3 | 3 |
| CON__ENSEMBL:ENS | | 2 | Q8K1I7 | Wipf1 | 2 |
| P04945 | | 1 | Q9JMG1 | Edf1 | 1 |
| Q9D1B9 | Mrpl28 | 3 | Q9R0Q3 | Tmed2 | 1 |
| Q9CWF2;Q | Tubb2b;Tul | 1 | Q3UKJ7 | Smu1 | 2 |
| Q8BMJ3;Q | Eif1ax;Eif1a | 3 | Q8BUE4 | Aifm2 | 1 |
| P47915 | Rpl29 | 3 | Q9CPQ1 | Cox6c | 1 |
| P01786 | | 1 | P16460 | Ass1 | 3 |
| Q9CZX2 | Cep89 | 1 | Q9DAS9 | Gng12 | 2 |
| P01810 | | 1 | Q9D0E1 | Hnrnpm | 3 |
| P62274 | Rps29 | 1 | Q3UX61;Q9 | Naa11;Naa1 | 2 |
| P18531 | Ighv3-6 | 1 | Q8VCW4 | Unc93b1 | 1 |
| P01629 | | 1 | P62827;Q6 | Ran;Rasl2-9 | 2 |
| Q99N95 | Mrpl3 | 2 | Q9DCF9 | Ssr3 | 1 |
| P62311 | Lsm3 | 2 | Q08943 | Ssrp1 | 2 |
| P01831 | Thy1 | 2 | Q921L3 | Tmco1 | 1 |
| P03976 | | 1 | Q9JKW0 | Arl6ip1 | 1 |
| Q99N89 | Mrpl43 | 2 | P56960 | Exosc10 | 1 |
| Q6ZWY3 | Rps27l | 2 | Q6ZQI3 | Mlec | 2 |
| P01647 | | 1 | Q8CHC4 | Synj1 | 1 |
| Q9DCU6 | Mrpl4 | 3 | P62307 | Snrpf | 2 |
| P62862 | Fau | 2 | Q01730 | Rsu1 | 2 |
| P01643 | | 1 | Q8C0L0 | Tmx4 | 1 |
| Q5SUE7 | Adad1 | 1 | Q64253 | Ly6e | 1 |
| Q8CBW3 | Abi1 | 5 | Q80XC2 | Trmt61a | 2 |
| P01749 | Ighv1-61 | 4 | Q9CY66 | Gar1 | 2 |
| Q8C6G8 | Wdr26 | 2 | P14428;P01 | H2-K1;H2-I | 1 |
| P62322 | Lsm5 | 2 | Q60973 | Rbbp7 | 1 |
| P01592 | Igj | 3 | P15379 | Cd44 | 1 |
| Q99N84 | Mrps18b | 3 | O54781 | Srpk2 | 2 |
| P59235 | Nup43 | 3 | Q9D958 | Spcs1 | 1 |
| Q9CQ06 | Mrpl24 | 4 | P62878 | Rbx1 | 1 |
| Q99LP6 | Grpel1 | 1 | Q9Z199;P6 | Supt4h1b;S | 3 |
| Q6A068 | Cdc5l | 3 | Q80U38 | Khnyn | 2 |
| Q9QWF0 | Chaf1a | 1 | Q9DCL9 | Paics | 2 |
| P26041;P26 | Msn;Rdx | 5 | Q9ESP1 | Sdf2l1 | 1 |
| Q9Z199;P6 | Supt4h1b;S | 3 | Q9JHE7 | Tssc4 | 1 |
| Q9R112 | Sqrdl | 2 | P84096 | Rhog | 2 |
| Q6WVG3 | Kctd12 | 2 | P01843 | | 3 |
| P62876 | Polr2l | 2 | P70245 | Ebp | 1 |
| Q9CQF8 | Mrpl57 | 1 | Q8BFZ3 | Actbl2 | 1 |
| Q9CQE6 | Asf1a | 3 | Q3UPF5 | Zc3hav1 | 2 |
| P06745 | Gpi | 1 | Q9R1J0 | Nsdhl | 2 |

| | | | | | |
|---|---|---|---|---|---|
| Q7TQE6 | Tmem57 | 1 | P53995 | Anapc1 | 3 |
| Q9CQQ8 | Lsm7 | 2 | P49300 | Clec10a | 3 |
| P01783;P18529 | | 1 | Q99P72 | Rtn4 | 2 |
| Q9DCG9 | Trmt112 | 2 | Q00651 | Itga4 | 1 |
| Q9CQE3 | Mrps17 | 2 | Q791V5 | Mtch2 | 1 |
| Q9D4H7 | Lonrf3 | 1 | P62075 | Timm13 | 1 |
| Q9ESP1 | Sdf2l1 | 2 | Q8VE97 | Srsf4 | 1 |
| Q64523;Q6 | Hist2h2ac;I | 1 | P35991 | Btk | 4 |
| Q9JIX0 | Eny2 | 2 | Q6A026 | Pds5a | 1 |
| Q923G2 | Polr2h | 2 | A1L314 | Mpeg1 | 3 |
| Q9D0D5 | Gtf2e1 | 3 | Q7TPR4 | Actn1 | 3 |
| P01656;P01654;P01655 | | 1 | Q61334 | Bcap29 | 1 |
| Q62189 | Snrpa | 1 | Q3V1L4 | Nt5c2 | 2 |
| Q924T2 | Mrps2 | 2 | Q9CWX9 | Ddx47 | 3 |
| Q9D1Q6 | Erp44 | 4 | P01942 | Hba | 1 |
| P70195 | Psmb7 | 2 | Q99LT0 | Dpy30 | 1 |
| Q9WVM3 | Anapc7 | 5 | Q8VE65 | Taf12 | 1 |
| Q9JL35 | Hmgn5 | 3 | Q8R0X7 | Sgpl1 | 2 |
| Q91V61 | Sfxn3 | 4 | Q9Z129 | Recql | 3 |
| Q62348 | Tsn | 3 | Q8BX90 | Fndc3a | 3 |
| Q60790 | Rasa3 | 6 | Q80ZS3 | Mrps26 | 2 |
| Q31125 | Slc39a7 | 3 | P70279 | Surf6 | 1 |
| Q8R1I1 | Uqcr10 | 1 | Q8BP67 | Rpl24 | 3 |
| Q91V41 | Rab14 | 3 | Q9WU03 | Spint2 | 1 |
| Q3U487 | Hectd3 | 3 | Q7TT50;Q8 | Cdc42bpb;A | 2 |
| P62313 | Lsm6 | 4 | Q64337 | Sqstm1 | 2 |
| Q9D773 | Mrpl2 | 4 | P50637 | Tspo | 1 |
| Q3TJZ6 | Fam98a | 2 | Q61768 | Kif5b | 1 |
| Q8BHD8 | Pcmtd2 | 4 | P01680 | | 1 |
| Q8R409 | Hexim1 | 2 | P43430 | Mcpt8 | 4 |
| Q99KI3 | Emc3 | 3 | Q7TMY4 | Thoc7 | 2 |
| Q9D1T0 | Lingo1 | 1 | Q9Z1N5;Q8 | Ddx39b;Dd | 4 |
| Q8R3N6 | Thoc1 | 3 | P28867;Q0 | Prkcd;Prkcq | 1 |
| O89051 | Itm2b | 2 | Q3UYV9 | Ncbp1 | 4 |
| Q9CQZ5 | Ndufa6 | 3 | Q8VBZ3 | Clptm1 | 1 |
| Q3UKJ7 | Smu1 | 3 | Q8BJ71 | Nup93 | 2 |
| Q9D0I8 | Mrto4 | 3 | P28660 | Nckap1 | 2 |
| O55028 | Bckdk | 4 | Q8CD15 | Mina | 1 |
| Q06185 | Atp5i | 3 | Q9WVM3 | Anapc7 | 2 |
| Q8JZX4 | Rbm17 | 3 | Q99JX7 | Nxf1 | 4 |
| Q99N92 | Mrpl27 | 2 | Q91YQ1;Q8 | Rab29;Rab3 | 1 |
| Q3TDD9 | Ppp1r21 | 4 | Q6DFW4 | Nop58 | 4 |
| Q91YK2 | Rrp1b | 1 | P01831 | Thy1 | 3 |
| P06327 | Gm5629 | 2 | Q8R3N6 | Thoc1 | 3 |
| P21550 | Eno3 | 1 | Q9ET30 | Tm9sf3 | 1 |

| | | | | | |
|---|---|---|---|---|---|
| Q9ERD7 | Tubb3 | 1 | Q9JKB3 | Ybx3 | 2 |
| P63073 | Eif4e | 1 | Q6P1F6 | Ppp2r2a | 3 |
| Q80WW9 | Ddrgk1 | 3 | P84102;O88 | Serf2;Serf1 | 2 |
| Q62241 | Snrpc | 2 | P10852 | Slc3a2 | 3 |
| Q8BYC6 | Taok3 | 4 | Q9EQH2 | Erap1 | 1 |
| P24547 | Impdh2 | 4 | Q9D071 | Mms19 | 1 |
| P16460 | Ass1 | 5 | Q9CPP0 | Npm3 | 1 |
| Q09014 | Ncf1 | 3 | Q8R3F5 | Mcat | 2 |
| P54818 | Galc | 3 | Q9WUQ2 | Preb | 2 |
| Q9D2G2 | Dlst | 2 | Q9R0Q4 | Morf4l2 | 1 |
| Q8K2C9 | Hacd3 | 2 | P69566 | Ranbp9 | 2 |
| Q99KQ4 | Nampt | 2 | Q9CR67 | Tmem33 | 1 |
| P52875 | Tmem165 | 2 | Q91ZW3 | Smarca5 | 3 |
| Q80XI3 | Eif4g3 | 3 | Q8BTV2 | Cpsf7 | 2 |
| Q9CQV5 | Mrps24 | 3 | O08585 | Clta | 1 |
| Q91YP3 | Dera | 3 | Q9D2E1 | Tssk3 | 1 |
| Q61510 | Trim25 | 3 | Q4VC33 | Maea | 1 |
| Q99JR1 | Sfxn1 | 6 | Q9QYJ3 | Dnajb1 | 2 |
| Q9D880 | Timm50 | 2 | Q504P2 | Clec12a | 1 |
| P49945;P29 | Ftl2;Ftl1 | 3 | Q9EQP2 | Ehd4 | 3 |
| Q9ERR7 | 15-Sep | 3 | Q4VA53 | Pds5b | 2 |
| P43430 | Mcpt8 | 4 | P11031 | Sub1 | 1 |
| Q80YD1 | Supv3l1 | 2 | P39053;P39 | Dnm1;Dnm | 1 |
| Q9JJZ2 | Tuba8 | 2 | Q62203 | Sf3a2 | 2 |
| Q6PE54 | Dhx40 | 2 | Q149F3;Q8 | Gspt2;Gspt | 1 |
| Q8VDP3 | Mical1 | 4 | Q62348 | Tsn | 2 |
| P0CW03;P0 | Ly6c2;Ly6c | 2 | P97742 | Cpt1a | 1 |
| P16879 | Fes | 2 | Q7TMY7 | Ipo8 | 1 |
| Q9EPE9 | Atp13a1 | 3 | Q9WUP7 | Uchl5 | 1 |
| Q9CY50 | Ssr1 | 3 | Q9D0I8 | Mrto4 | 5 |
| Q9CQN7 | Mrpl41 | 4 | CON__P35527 | | 3 |
| Q8R0A0 | Gtf2f2 | 2 | Q9CR62 | Slc25a11 | 3 |
| Q9R1Q6 | Tmem176b | 2 | Q60629;Q6 | Epha5;Epha | 1 |
| Q921F4 | Hnrnpll | 2 | P18337 | Sell | 1 |
| Q8BVE3 | Atp6v1h | 2 | Q9JJI8 | Rpl38 | 1 |
| Q9Z2A5 | Ate1 | 1 | Q8C547 | Heatr5b | 1 |
| A2A935 | Prdm16 | 1 | P29621;Q91 | Serpina3c;S | 1 |
| Q80UP3 | Dgkz | 5 | Q9JII5 | Dazap1 | 3 |
| Q80UW8 | Polr2e | 3 | P61211 | Arl1 | 1 |
| Q9DD18 | Dtd1 | 2 | Q9DB05 | Napa | 1 |
| Q8BYL4 | Yars2 | 2 | Q3V0C5 | Usp48 | 1 |
| O88622 | Parg | 2 | Q6PHQ8 | Naa35 | 3 |
| CON__P07477 | | 1 | Q9ERU9 | Ranbp2 | 3 |
| Q3UV17 | Krt76 | 1 | Q99PM9 | Uck2 | 1 |
| Q8BLR5 | Psd4 | 2 | Q9R0U0 | Srsf10 | 4 |

| | | | | | |
|---|---|---|---|---|---|
| Q9CQI7 | Snrpb2 | 1 | Q61735 | Cd47 | 1 |
| P61027 | Rab10 | 2 | P62309 | Snrpg | 2 |
| Q921X9 | Pdia5 | 3 | CON__P12763 | | 2 |
| Q8BU88 | Mrpl22 | 2 | Q9Z1T1 | Ap3b1 | 2 |
| Q91YT7 | Ythdf2 | 2 | Q8R349 | Cdc16 | 2 |
| Q99ME9 | Gtpbp4 | 2 | P04944;P04941;P0494 | | 1 |
| Q3TZX8 | Nol9 | 2 | Q91YI1 | Atg13 | 2 |
| O08784 | Tcof1 | 3 | Q8K209 | Gpr56 | 3 |
| Q9QZQ8 | H2afy | 4 | Q9D0M3 | Cyc1 | 4 |
| Q8BTI7 | Ankrd52 | 2 | P60060 | Sec61g | 2 |
| P63101 | Ywhaz | 1 | Q9QYS9 | Qki | 1 |
| Q99LL5 | Pwp1 | 2 | Q5SWD9 | Tsr1 | 3 |
| Q8BTW3 | Exosc6 | 2 | Q99M87 | Dnaja3 | 3 |
| Q8K2Y7 | Mrpl47 | 3 | P18527 | | 1 |
| P99028 | Uqcrh | 1 | Q8VCM8 | Ncln | 1 |
| Q6ZQB6 | Ppip5k2 | 3 | P59016 | Vps33b | 1 |
| O89098 | Cst7 | 2 | Q99PG2 | Ogfr | 1 |
| Q8R5F3 | Oard1 | 2 | Q61074 | Ppm1g | 2 |
| Q9CXU9;P | Eif1b;Eif1 | 2 | P34884 | Mif | 1 |
| Q9Z2D8 | Mbd3 | 1 | O35598 | Adam10 | 1 |
| Q8QZY9 | Sf3b4 | 3 | Q9Z1E4 | Gys1 | 1 |
| D3YZU1 | Shank1 | 1 | P51150 | Rab7a | 1 |
| Q80ZS3 | Mrps26 | 2 | O55131 | 7-Sep | 1 |
| Q3TDQ1 | Stt3b | 2 | Q9EQQ2 | Yipf5 | 1 |
| P51863 | Atp6v0d1 | 3 | P49586;Q8 | Pcyt1a;Pcyt1 | 1 |
| Q8CHP5 | Wibg | 2 | Q9CQB5 | Cisd2 | 1 |
| P23249 | Mov10 | 4 | Q9EPE9 | Atp13a1 | 2 |
| O08573 | Lgals9 | 3 | P47758 | Srprb | 2 |
| Q8BW41 | Pomgnt2 | 2 | Q9DCD2 | Xab2 | 2 |
| Q921M4 | Golga2 | 3 | Q571I4 | Sgk223 | 1 |
| A2AQ19 | Rtf1 | 3 | Q9CRA8 | Exosc5 | 1 |
| Q3UH60 | Dip2b | 2 | P70218 | Map4k1 | 2 |
| Q99JP4 | Cdc26 | 1 | Q61103 | Dpf2 | 1 |
| Q8C0L0 | Tmx4 | 3 | Q00612;RE | G6pdx;G6p | 2 |
| P52432 | Polr1c | 4 | Q8K2H6 | Anapc10 | 1 |
| O88532 | Zfr | 2 | D3Z6Q9 | Bin2 | 1 |
| Q3UHX2 | Pdap1 | 3 | Q8VDF2 | Uhrf1 | 2 |
| Q9WTX6 | Cul1 | 4 | P23249 | Mov10 | 2 |
| Q9QYJ3 | Dnajb1 | 4 | Q8VE22 | Mrps23 | 4 |
| O54774 | Ap3d1 | 3 | P01638;P01637 | | 1 |
| Q9DCS9 | Ndufb10 | 2 | Q9D023 | Mpc2 | 1 |
| Q9CWW6 | Pin4 | 2 | Q8CJ53 | Trip10 | 1 |
| Q9CQV1 | Pam16 | 2 | Q99J64 | Rbm43 | 1 |
| Q9QXK3 | Copg2 | 3 | Q8K2C9 | Hacd3 | 1 |
| Q9CPS7 | Pno1 | 2 | Q78XF5 | Ostc | 1 |

| | | | | | |
|---|---|---|---|---|---|
| Q9DA97;O5 | Sept14;Sept | 1 | Q91VR5 | Ddx1 | 2 |
| Q8BVU0 | Lrch3 | 2 | Q61168 | Laptm5 | 1 |
| P63166 | Sumo1 | 2 | Q8BJF9 | Chmp2b | 2 |
| P11835 | Itgb2 | 2 | Q9ERK4 | Cse1l | 1 |
| P58059 | Mrps21 | 1 | P80313 | Cct7 | 2 |
| Q9DCL2 | Fam96a | 2 | P00688;P0 | Amy2;Amy | 2 |
| P48193 | Epb41 | 1 | Q8BX57 | Pxk | 1 |
| Q8C1Z8 | Trmt10a | 1 | Q8K4L0 | Ddx54 | 4 |
| Q9D187 | Fam96b | 3 | Q8R480 | Nup85 | 2 |
| Q6R891 | Ppp1r9b | 2 | Q9Z315 | Sart1 | 2 |
| Q9JLZ3 | Auh | 2 | P51863 | Atp6v0d1 | 2 |
| Q80XN0 | Bdh1 | 2 | Q9DBS1 | Tmem43 | 4 |
| Q78XF5 | Ostc | 1 | Q9D0Q7 | Mrpl45 | 4 |
| Q8K003 | Tma7 | 1 | Q6A068 | Cdc5l | 1 |
| Q9DAA6 | Exosc1 | 2 | Q925I1 | Atad3 | 4 |
| Q9D1H8 | Mrpl53 | 2 | P28658 | Atxn10 | 2 |
| Q6P9Q6 | Fkbp15 | 2 | Q8BN21 | Vrk2 | 1 |
| Q69ZQ2 | Isy1 | 2 | P51807 | Dynlt1 | 2 |
| Q64737 | Gart | 5 | Q64213 | Sf1 | 3 |
| Q99PM9 | Uck2 | 2 | P12815 | Pdcd6 | 2 |
| Q3UKC1 | Tax1bp1 | 3 | Q8BP48 | Metap1 | 3 |
| Q91WN1 | Dnajc9 | 4 | Q8BG07 | Pld4 | 6 |
| Q8R5C5 | Actr1b | 2 | Q91VK4 | Itm2c | 1 |
| Q9D1N9 | Mrpl21 | 3 | P97760 | Polr2c | 5 |
| Q9D1G1 | Rab1b | 1 | P43247 | Msh2 | 3 |
| Q99N85 | Mrps18a | 2 | O88622 | Parg | 2 |
| Q9CY58 | Serbp1 | 2 | Q8VDD9 | Phip | 1 |
| Q9DBR1 | Xrn2 | 3 | Q60876 | Eif4ebp1 | 1 |
| Q9D8P4 | Mrpl17 | 2 | Q9CXY9 | Pigk | 1 |
| O35130 | Emg1 | 2 | Q9CXW3 | Cacybp | 1 |
| Q8R3F5 | Mcat | 2 | Q08288 | Lyar | 2 |
| Q99JH1 | Rpp25l | 3 | Q5SWU9 | Acaca | 2 |
| Q91WD1 | Polr3d | 2 | Q9D7A6 | Srp19 | 1 |
| Q9CZ04 | Cops7a | 2 | O08992 | Sdcbp | 1 |
| Q80TP3 | Ubr5 | 9 | Q61136 | Prpf4b | 4 |
| D3Z6Q9 | Bin2 | 3 | Q99JH1 | Rpp25l | 1 |
| Q8QZZ7 | Tprkb | 2 | Q62376 | Snrnp70 | 1 |
| Q6A0A9 | FAM120A | 3 | Q8K224 | Nat10 | 4 |
| O89103 | Cd93 | 3 | Q6P9N1 | Fam126a | 1 |
| Q8CJ53 | Trip10 | 2 | P97310 | Mcm2 | 3 |
| Q99P72 | Rtn4 | 2 | A2RSY6 | Trmt1l | 1 |
| P97353 | Sec1 | 1 | Q9D338 | Mrpl19 | 2 |
| P23198 | Cbx3 | 3 | P26369 | U2af2 | 3 |
| Q9DBB4 | Naa16 | 1 | Q8BFT2 | Haus4 | 1 |
| P39749 | Fen1 | 1 | Q9Z321 | Top3b | 2 |

| | | | | | |
|---|---|---|---|---|---|
| Q9CQL4 | Mrpl20 | 1 | P58742 | Aaas | 1 |
| Q9CXF4 | Tbc1d15 | 3 | Q3U487 | Hectd3 | 1 |
| Q99PW4 | Tp53rk | 2 | O54825 | Bysl | 2 |
| P40630 | Tfam | 2 | Q91VI7 | Rnh1 | 3 |
| Q64131;Q0 | Runx3;Run | 1 | P42208 | 2-Sep | 1 |
| P01628;P01 | Gm5153 | 1 | P46061 | Rangap1 | 2 |
| Q8C4Y3 | Nelfb | 2 | Q8VEE4 | Rpa1 | 2 |
| Q8R001 | Mapre2 | 1 | P50543 | S100a11 | 1 |
| P55096 | Abcd3 | 2 | Q60865 | Caprin1 | 1 |
| Q9Z2I0 | Letm1 | 2 | O88543 | Cops3 | 1 |
| Q8K273 | Mmgt1 | 1 | Q9CSN1 | Snw1 | 1 |
| P62488 | Polr2g | 3 | Q91V92 | Acly | 1 |
| Q5XJE5 | Leo1 | 2 | Q9EPU4 | Cpsf1 | 4 |
| Q91WD5 | Ndufs2 | 2 | P56399 | Usp5 | 1 |
| Q9CXE7 | Tmed5 | 1 | Q9EQI8 | Mrpl46 | 3 |
| P63242;Q8 | Eif5a;Eif5a | 2 | Q9D168 | Ints12 | 1 |
| Q8VHE0 | Sec63 | 3 | Q922Q4 | Pycr2 | 1 |
| P50431 | Shmt1 | 1 | Q8BH59 | Slc25a12 | 1 |
| Q9QY06 | Myo9b | 2 | E9PV87 | Talpid3 | 1 |
| P12815 | Pdcd6 | 2 | Q61696;P17 | Hspa1a;Hsp | 1 |
| P21619 | Lmnb2 | 1 | Q91WN1 | Dnajc9 | 2 |
| Q61584;P35 | Fxr1;Fmr1;F | 2 | Q62426 | Cstb | 1 |
| B2RY04 | Dock5 | 2 | Q99J39 | Mlycd | 1 |
| E9PVA8 | Gcn1l1 | 4 | Q99LE6 | Abcf2 | 2 |
| Q922H2 | Pdk3 | 1 | Q9D1C9 | Rrp7a | 2 |
| Q9CQ40 | Mrpl49 | 4 | Q9CQC7 | Ndufb4 | 2 |
| Q3UMY5 | Eml4 | 2 | Q8BYC6 | Taok3 | 3 |
| Q9CQW0 | Emc6 | 2 | Q61581 | Igfbp7 | 2 |
| Q7TMB8 | Cyfip1 | 3 | P01642 | Gm10881 | 2 |
| Q80ZK0 | Mrps10 | 2 | Q9D024 | Ccdc47 | 6 |
| P42125 | Eci1 | 1 | Q9WTS2 | Fut8 | 2 |
| Q9D2E2 | Toe1 | 3 | O08900 | Ikzf3 | 1 |
| Q8K4I3;Q9 | Arhgef6;Ar | 3 | P40630 | Tfam | 1 |
| Q8K1A6 | Cc2d1a | 1 | Q8C2E7 | Kiaa0196 | 2 |
| O35658 | C1qbp | 1 | P70315 | Was | 4 |
| Q9ERI2 | Rab27a | 3 | Q8VDP6 | Cdipt | 1 |
| Q3U2A8 | Vars2 | 2 | Q9JLV5 | Cul3 | 2 |
| Q8C5Q4 | Grsf1 | 1 | Q80UP3 | Dgkz | 3 |
| E9Q4Z2 | Acacb | 2 | Q9CQW0 | Emc6 | 1 |
| Q9ER69 | Wtap | 2 | Q689Z5 | Sbno1 | 1 |
| Q91XI1 | Dus3l | 3 | CON__Q3ZBS7 | | 1 |
| Q8CD15 | Mina | 1 | O35954 | Pitpnm1 | 1 |
| P58137 | Acot8 | 4 | Q62189 | Snrpa | 1 |
| Q9QZV9;Q | Nxt1;Nxt2 | 3 | Q8BVK9 | Sp110 | 1 |
| CON__ENSEMBL:ENS | | 2 | Q8CI33 | Cwf19l1 | 1 |

| | | | | | |
|---|---|---|---|---|---|
| Q80XC2 | Trmt61a | 2 | Q8VCG3 | Wdr74 | 1 |
| P84102;O8 | Serf2;Serf1 | 1 | Q91WC0 | Setd3 | 1 |
| P84096 | Rhog | 1 | Q91YK2 | Rrp1b | 1 |
| Q8BX09 | Rbbp5 | 2 | Q91YU8 | Ppan | 1 |
| Q3UI43 | Babam1 | 2 | Q99K95 | Rtfdc1 | 1 |
| Q3TUU5 | Tex30 | 3 | Q99KG3 | Rbm10 | 1 |
| Q80U72 | Scrib | 3 | Q9CQ40 | Mrpl49 | 1 |
| Q7TNP2 | Ppp2r1b | 1 | Q9CWS4 | Cpsf3l | 1 |
| Q9Z129 | Recql | 1 | Q9CYL5 | Glipr2 | 1 |
| Q9QZD8 | Slc25a10 | 2 | Q9DCS9 | Ndufb10 | 1 |
| Q9CR67 | Tmem33 | 1 | Q6WVG3 | Kctd12 | 1 |
| Q922F4 | Tubb6 | 3 | Q8CCJ3 | Ufl1 | 1 |
| Q99PU8 | Dhx30 | 3 | Q9Z1Q5 | Clic1 | 1 |
| O70325;Q9 | Gpx4 | 1 | Q8BY87 | Usp47 | 1 |
| Q9EQ61 | Pes1 | 3 | Q80T69 | Rsbn1 | 1 |
| P51912 | Slc1a5 | 2 | Q9DBE8 | Alg2 | 1 |
| Q9JKR6 | Hyou1 | 3 | CON__A2A | Krt15;Krt14 | 1 |
| P30285 | Cdk4 | 2 | Q9Z2I0 | Letm1 | 1 |
| Q9Z0G0 | Gipc1 | 3 | Q8R1J3 | Zcchc9 | 1 |
| Q62203 | Sf3a2 | 1 | P60898 | Polr2i | 1 |
| Q8C5L6 | Inpp5k | 2 | Q921I9 | Exosc4 | 1 |
| P97352 | S100a13 | 1 | Q9D198 | Syf2 | 1 |
| Q62465 | Vat1 | 3 | P63242 | Eif5a | 1 |
| Q99L47 | St13 | 2 | Q3THK3 | Gtf2f1 | 1 |
| P70460 | Vasp | 4 | Q9DBD5 | Pelp1 | 1 |
| P97742 | Cpt1a | 2 | Q5SSI6 | Utp18 | 1 |
| Q9CPQ3 | Tomm22 | 2 | Q9CQL4 | Mrpl20 | 1 |
| Q8BUN5;Q | Smad3;Sma | 2 | Q9D868 | Ppih | 1 |
| Q64261 | Cdk6 | 3 | Q9ERR7 | 15-Sep | 1 |
| E9Q5K9 | Ythdc1 | 1 | Q8R3F9 | Tut1 | 1 |
| Q8BH57 | Wdr48 | 2 | Q80TY0 | Fnbp1 | 1 |
| Q9JJG9 | Noa1 | 1 | Q8BLY2 | Tarsl2 | 1 |
| Q9DA08 | Ccdc101 | 1 | Q9CQS2 | Nop10 | 1 |
| Q9CZX9 | Emc4 | 1 | Q62432;Q8 | Smad2;Sma | 2 |
| O70152 | Dpm1 | 1 | Q9QZL0 | Ripk3 | 1 |
| Q9D903 | Ebna1bp2 | 1 | P01741 | | 1 |
| Q8C7D2 | Crbn | 1 | Q9EP82 | Wdr4 | 1 |
| P30999 | Ctnnd1 | 2 | O35405 | Pld3 | 1 |
| P47856;Q9 | Gfpt1;Gfpt2 | 2 | Q91YN0 | D6Wsu163e | 1 |
| P61211 | Arl1 | 2 | Q3UKC1 | Tax1bp1 | 1 |
| Q80ZF8 | Bai3 | 1 | P97298 | Serpinf1 | 1 |
| Q60631 | Grb2 | 2 | Q68ED3 | Papd5 | 1 |
| P01667 | | 1 | Q9JIW9 | Ralb | 1 |
| O88456 | Capns1 | 1 | P35922;Q9 | Fmr1;Fxr2; | 1 |
| Q9WUD1 | Stub1 | 2 | A2A6Q5 | Cdc27 | 1 |

| | | | | | |
|---|---|---|---|---|---|
| Q91W18 | Tdrd3 | 3 | Q920C4 | Ms4a3 | 1 |
| Q6ZPJ3 | Ube2o | 2 | Q9D7M8 | Polr2d | 1 |
| Q8VEA4 | Chchd4 | 1 | Q6NZB0 | Dnajc8 | 1 |
| P52503 | Ndufs6 | 1 | Q8BMQ2 | Gtf3c4 | 1 |
| P01728 | | 1 | E9PZJ8 | Ascc3 | 1 |
| Q3UPF5 | Zc3hav1 | 1 | P81269;Q01 | Atf1;Creb1 | 1 |
| Q78IK2 | Usmg5 | 1 | P97470 | Ppp4c | 2 |
| P51174 | Acadl | 1 | Q8R0S2 | Iqsec1 | 1 |
| Q3TDN2 | Faf2 | 1 | Q2VPQ9 | Meaf6 | 1 |
| Q80U93 | Nup214 | 2 | Q9DCT6 | Bap18 | 1 |
| Q9CQS5 | Riok2 | 1 | P01652 | | 1 |
| O70126 | Aurkb | 2 | Q5XJE5 | Leo1 | 1 |
| Q7TSQ8 | Pdpr | 2 | Q921E6 | Eed | 1 |
| P01804 | | 1 | O70472 | Tmem131 | 1 |
| Q921I9 | Exosc4 | 2 | Q9Z2U0;Q9 | Psma7;Psm | 1 |
| P70698 | Ctps1 | 2 | Q9CQ75 | Ndufa2 | 2 |
| Q9D7M1 | Gid8 | 2 | Q9CQ06 | Mrpl24 | 2 |
| Q6PFR5 | Tra2a | 1 | Q9EQ06 | Hsd17b11 | 1 |
| Q9WTX8 | Mad1l1 | 3 | P28574 | Max | 1 |
| Q6PAR5 | Gapvd1 | 2 | Q9DC71 | Mrps15 | 1 |
| P40124 | Cap1 | 1 | Q91YP3 | Dera | 1 |
| Q80UP5 | Ankrd13a | 2 | Q91VL9 | Zbtb1 | 1 |
| Q8VBV3 | Exosc2 | 2 | P63087;P62 | Ppp1cc;Ppp | 2 |
| Q6NZB0 | Dnajc8 | 1 | Q8CD10 | Micu2 | 1 |
| P06800 | Ptprc | 2 | Q6PAQ4 | Rexo4 | 1 |
| Q9CQH3 | Ndufb5 | 1 | Q03963 | Eif2ak2 | 1 |
| Q689Z5 | Sbno1 | 2 | Q9QXK7 | Cpsf3 | 1 |
| Q791N7 | Znrd1 | 1 | P63328;P48 | Ppp3ca;Ppp | 1 |
| Q9WUP7 | Uchl5 | 1 | P56581 | Tdg | 1 |
| Q9JLQ2;Q6 | Git2;Git1 | 2 | Q9CX86 | Hnrnpa0 | 2 |
| Q61687 | Atrx | 1 | Q9JJH7 | Trpm5 | 1 |
| P61750 | Arf4 | 2 | Q9Z2Z6 | Slc25a20 | 1 |
| Q8BGC0 | Htatsf1 | 1 | Q6P5B0 | Rrp12 | 2 |
| O09043 | Napsa | 1 | Q8CH25 | Sltm | 1 |
| P24788 | Cdk11b | 2 | Q8CI75 | Dis3l2 | 3 |
| Q4QRL3 | Ccdc88b | 3 | Q9WU56 | Pus1 | 2 |
| Q8BMQ2 | Gtf3c4 | 4 | Q91VH2 | Snx9 | 1 |
| Q8BGW1 | Fto | 1 | Q91W18 | Tdrd3 | 1 |
| Q9CR68 | Uqcrfs1 | 1 | Q63844 | Mapk3 | 1 |
| Q9QUM0 | Itga2b | 3 | Q9D2E2 | Toe1 | 1 |
| Q923T9;P11 | Camk2g;Ca | 2 | Q91V61 | Sfxn3 | 2 |
| Q6ZQ88 | Kdm1a | 2 | Q8BXV2 | Bri3bp | 1 |
| Q8BYB9 | Poglut1 | 3 | Q9ERF3 | Wdr61 | 1 |
| Q8K1C0 | Angel2 | 1 | P21995 | Emb | 1 |
| P21995 | Emb | 2 | Q9CPX7 | Mrps16 | 1 |

| | | | | | |
|---|---|---|---|---|---|
| Q5U4E2 | Repin1 | 3 | Q8BW10 | Nob1 | 1 |
| P58742 | Aaas | 2 | P61290 | Psme3 | 1 |
| Q99P91 | Gpnmb | 1 | E9Q5K9 | Ythdc1 | 1 |
| Q8C079 | Strip1 | 2 | Q923W1 | Tgs1 | 1 |
| Q99LY9 | Ndufs5 | 1 | P01803 | | 1 |
| Q9D103 | Ifitm1 | 1 | Q7TSI3 | Ppp6r1 | 1 |
| Q8JZN5 | Acad9 | 2 | Q8CCP0 | Nemf | 1 |
| Q60597 | Ogdh | 2 | O55013 | Trappc3 | 1 |
| P42225 | Stat1 | 3 | Q9CZD3 | Gars | 1 |
| Q922R5 | Smek2 | 2 | Q3U2A8 | Vars2 | 1 |
| Q8BUK6 | Hook3 | 3 | Q99LQ1 | Mbip | 1 |
| Q9DBG7 | Srpr | 5 | P70398 | Usp9x | 1 |
| Q60649 | Clpb | 1 | Q8BWJ3 | Phka2 | 1 |
| O35295 | Purb | 1 | Q80TR8 | Vprbp | 2 |
| Q8K209 | Gpr56 | 3 | Q8R3H7 | Hs2st1 | 1 |
| P61804 | Dad1 | 2 | Q9D6N5 | Drap1 | 1 |
| P53994;P59 | Rab2a;Rab2 | 2 | Q91Z92 | B3galt6 | 1 |
| O70404 | Vamp8 | 1 | Q64705 | Usf2 | 1 |
| O35368 | Ifi203 | 1 | P35601 | Rfc1 | 1 |
| Q9CWK3 | Cd2bp2 | 2 | Q922H9 | Znf330 | 1 |
| P03975 | Iap | 1 | Q3U4G3 | Xxylt1 | 2 |
| Q3UIR3 | Dtx3l | 2 | P62488 | Polr2g | 2 |
| Q80ZQ9 | Fam206a | 2 | P49290 | Epx | 1 |
| Q9Z2G6 | Sel1l | 1 | P26043 | Rdx | 2 |
| Q8K4L0 | Ddx54 | 2 | Q8BL74 | Gtf3c2 | 1 |
| Q8BZQ7 | Anapc2 | 2 | P48193 | Epb41 | 2 |
| Q07832 | Plk1 | 3 | O35218 | Cpsf2 | 2 |
| P50136 | Bckdha | 1 | Q8K4P0 | Wdr33 | 1 |
| Q8C156 | Ncaph | 3 | Q8BLR5 | Psd4 | 1 |
| P97452 | Bop1 | 2 | P01786;P01790;P01789; | | 1 |
| P27870 | Vav1 | 3 | Q80XI3 | Eif4g3 | 1 |
| G5E861 | Sclt1 | 1 | P01669;P01667;P01666 | | 1 |
| P97304 | Polr1d | 1 | P56959 | Fus | 1 |
| Q9D832 | Dnajb4 | 2 | Q60737 | Csnk2a1 | 2 |
| P27546 | Map4 | 1 | Q91XB0 | Trex1 | 3 |
| CON__ENSEMBL:ENS | | 2 | Q8VCK3;P8 | Tubg2;Tubg | 1 |
| Q8VE62 | Paip1 | 1 | O55098 | Stk10 | 2 |
| P01803 | | 1 | Q3UJD6 | Usp19 | 1 |
| Q922L6 | Nelfcd | 2 | Q9D103 | Ifitm1 | 1 |
| Q99JW4 | Lims1 | 1 | P58137 | Acot8 | 1 |
| P46737 | Brcc3 | 1 | P70700 | Polr1b | 2 |
| Q8CI94 | Pygb | 2 | P01673;P01671 | | 1 |
| Q91WC0 | Setd3 | 1 | Q9QYC1 | Pcnx | 1 |
| Q9D4H8 | Cul2 | 2 | Q4VBE8 | Wdr18 | 1 |
| Q9EQC5 | Scyl1 | 3 | Q8C5L6 | Inpp5k | 2 |

| | | | | | |
|---|---|---|---|---|---|
| Q9EP97 | Senp3 | 1 | Q64523;Q6 | Hist2h2ac;I | 1 |
| Q80YP0;P9 | Cdk3;Cdk2 | 2 | Q7TMF2 | Eri1 | 1 |
| Q9CRC0 | Vkorc1 | 1 | Q8BHL3 | Tbc1d10b | 1 |
| Q9D4J7 | Phf6 | 1 | P56183 | Rrp1 | 1 |
| Q6PHQ8 | Naa35 | 2 | P16330 | Cnp | 1 |
| Q9Z110 | Aldh18a1 | 2 | Q8CGN5 | Plin1 | 1 |
| P69566 | Ranbp9 | 2 | Q8CG46 | Smc5 | 1 |
| Q99LM2 | Cdk5rap3 | 3 | Q99KI3 | Emc3 | 1 |
| Q7TMF2 | Eri1 | 1 | Q5BLK4 | Zcchc6 | 2 |
| Q9DBY8 | Nvl | 3 | P20491 | Fcer1g | 1 |
| P09528 | Fth1 | 1 | P11680 | Cfp | 2 |
| Q9DBZ1 | Ikbip | 1 | Q9JKC8;Q8 | Ap3m1;Ap3 | 2 |
| Q9Z1R2 | Bag6 | 1 | P67871 | Csnk2b | 2 |
| Q91YE6 | Ipo9 | 3 | P07742 | Rrm1 | 2 |
| A2RSY6 | Trmt1l | 1 | Q8K273 | Mmgt1 | 1 |
| Q9CQS2 | Nop10 | 1 | Q6NZJ6 | Eif4g1 | 1 |
| Q8BMZ5 | Tsen34 | 3 | P01663;P01662 | | 1 |
| Q9D824 | Fip1l1 | 1 | P62311 | Lsm3 | 2 |
| Q9CPP6 | Ndufa5 | 2 | Q9D3R6;Q | Katnal2;Spa | 1 |
| O88351 | Ikbkb | 2 | Q9DBR1 | Xrn2 | 2 |
| Q9Z103 | Adnp | 1 | A6PWY4 | Wdr76 | 1 |
| Q810D6 | Grwd1 | 2 | Q8BHD8 | Pcmtd2 | 3 |
| O55013 | Trappc3 | 2 | CON__P01030;CON__ | | 1 |
| Q8VC85 | Lsm1 | 2 | P15089 | Cpa3 | 2 |
| Q921W4 | Cryzl1 | 2 | Q8BM55 | Tmem214 | 2 |
| Q9D7M8 | Polr2d | 2 | Q9CY73 | Mrpl44 | 1 |
| Q8R2U4 | Ntmt1 | 2 | O35972 | Mrpl23 | 2 |
| Q3U2P1 | Sec24a | 3 | Q8BSY0 | Asph | 1 |
| Q8K296 | Mtmr3 | 1 | Q8C5Q4 | Grsf1 | 1 |
| Q9Z1J3 | Nfs1 | 1 | O88532 | Zfr | 1 |
| Q9Z1Q5 | Clic1 | 2 | Q9CQ49 | Ncbp2 | 1 |
| Q9D2N9 | Vps33a | 2 | P52432 | Polr1c | 2 |
| Q9D8T7 | Slirp | 1 | Q8QZY9 | Sf3b4 | 1 |
| Q8R307 | Vps18 | 2 | Q8BH61 | F13a1 | 3 |
| Q9CYH6 | Rrs1 | 1 | Q8R5K4 | Nol6 | 2 |
| Q99K41 | Emilin1 | 2 | Q6Y7W8 | Gigyf2 | 1 |
| Q91ZW3;Q( | Smarca5;Sn | 2 | Q9D0D5 | Gtf2e1 | 1 |
| P62962 | Pfn1 | 1 | Q9WV02 | Rbmx | 1 |
| Q3UJD6 | Usp19 | 1 | Q9CY58 | Serbp1 | 2 |
| Q8BI72 | Cdkn2aip | 1 | Q61464 | Znf638 | 2 |
| Q80TH2 | Erbb2ip | 1 | Q9CZ57 | Nsun4 | 2 |
| B2RWS6 | Ep300 | 1 | Q9Z126 | Pf4 | 1 |
| Q8CD10 | Micu2 | 1 | Q7TSH2 | Phkb | 3 |
| Q8BWJ3 | Phka2 | 1 | Q9DCU6 | Mrpl4 | 2 |
| Q8CDM1 | Atad2 | 1 | Q3UMB9 | Kiaa1033 | 4 |

| | | | | | |
|---|---|---|---|---|---|
| O70145 | Ncf2 | 2 | Q60790 | Rasa3 | 4 |
| Q8BK64 | Ahsa1 | 1 | Q9DCI9 | Mrpl32 | 1 |
| O70194 | Eif3d | 1 | O70378 | Emc8 | 2 |
| Q8CEC0 | Nup88 | 2 | Q9D1N9 | Mrpl21 | 2 |
| O54825 | Bysl | 2 | Q9ESX5 | Dkc1 | 3 |
| Q9ET30 | Tm9sf3 | 2 | Q9CZX0 | Elp3 | 4 |
| Q9JHS4 | Clpx | 2 | D0QMC3 | Mndal | 1 |
| Q8BGS0 | Mak16 | 1 | Q3U2P1 | Sec24a | 4 |
| Q8CBY8 | Dctn4 | 3 | Q9D0G0 | Mrps30 | 3 |
| P70388 | Rad50 | 2 | P31996 | Cd68 | 1 |
| Q8C1D8 | Iws1 | 1 | P04945 | | 1 |
| Q8BHJ5 | Tbl1xr1 | 1 | Q8VHL1 | Setd7 | 2 |
| Q69ZS7 | Hbs1l | 1 | Q8R2U4 | Ntmt1 | 2 |
| Q9D0N7 | Chaf1b | 2 | Q9R1Q7 | Plp2 | 1 |
| Q80YV3 | Trrap | 2 | Q91XI1 | Dus3l | 3 |
| O89023 | Tpp1 | 1 | Q99N92 | Mrpl27 | 1 |
| Q3UPL0 | Sec31a | 2 | O55028 | Bckdk | 2 |
| Q8BYZ1 | Abi3 | 1 | Q8VC03 | Eml3 | 1 |
| Q91X20 | Ash2l | 1 | P62322 | Lsm5 | 1 |
| P05555 | Itgam | 2 | P83940 | Tceb1 | 2 |
| Q9D071 | Mms19 | 2 | Q99N85 | Mrps18a | 2 |
| Q9D8V0 | Hm13 | 1 | Q8BHN1 | Txlng | 2 |
| Q9Z0H3 | Smarcb1 | 3 | Q9Z0H1 | Wdr46 | 2 |
| Q9WTZ1 | Rnf7 | 1 | P35441 | Thbs1 | 3 |
| P15702 | Spn | 1 | Q9CRB2 | Nhp2 | 2 |
| Q8K1R3 | Pnpt1 | 1 | O08573 | Lgals9 | 2 |
| P97371 | Psme1 | 1 | CON__ENSEMBL:ENS | | 1 |
| Q9CZN8 | Qrsl1 | 2 | Q9D6Z1 | Nop56 | 2 |
| O88554 | Parp2 | 1 | Q63871 | Polr2k | 1 |
| Q6ZPR6 | Ibtk | 2 | Q8BFR5 | Tufm | 4 |
| Q9QZB7 | Actr10 | 1 | P01750 | | 1 |
| Q8BNV1 | Trmt2a | 1 | P58059 | Mrps21 | 2 |
| Q8BGA7 | | 1 | Q99PU8 | Dhx30 | 4 |
| P62141 | Ppp1cb | 1 | Q8K2M0 | Mrpl38 | 2 |
| Q2TBA3 | Malt1 | 1 | P28740 | Kif2a | 2 |
| Q99LC8 | Eif2b1 | 1 | Q9DCA2 | Mrps11 | 2 |
| Q9QYS9 | Qki | 1 | P62869 | Tceb2 | 1 |
| Q03347 | Runx1 | 1 | P01636 | | 2 |
| O08749 | Dld | 1 | P70280 | Vamp7 | 1 |
| P50427 | Sts | 1 | Q62036 | Cep131 | 2 |
| P63328;P48 | Ppp3ca;Ppp | 2 | Q5U458 | Dnajc11 | 3 |
| P01652 | | 1 | Q9CQV5 | Mrps24 | 3 |
| Q9D168 | Ints12 | 1 | Q7TSC1 | Prrc2a | 2 |
| Q8BN21 | Vrk2 | 1 | Q99J36 | Thumpd1 | 5 |
| Q5KU39 | Vps41 | 1 | Q9Z1K5 | Arih1 | 2 |

| | | | | | |
|---|---|---|---|---|---|
| Q91XU3 | Pip4k2c | 1 | Q8BU88 | Mrpl22 | 2 |
| P18242 | Ctsd | 1 | Q9D773 | Mrpl2 | 3 |
| Q9JJF3 | No66 | 1 | Q4VAA2 | Cdv3 | 1 |
| Q5U458 | Dnajc11 | 2 | Q6PFR5 | Tra2a | 1 |
| Q8R3Q0 | Saraf | 1 | P03976 | | 1 |
| Q9D706 | Rpap3 | 1 | Q9QZQ8 | H2afy | 3 |
| Q8BQZ5 | Cpsf4 | 1 | Q99JF8 | Psip1 | 2 |
| O35671 | Itgb1bp1 | 2 | Q921R4 | Dnajc14 | 1 |
| O35904 | Pik3cd | 1 | Q80XL6 | Acad11 | 3 |
| Q9D7H3 | RtcA | 2 | Q8BIJ7 | Rufy1 | 1 |
| P52479 | Usp10 | 3 | P08905;P17 | Lyz2;Lyz1 | 2 |
| Q08481 | Pecam1 | 1 | Q60960;O3 | Kpna1;Kpna | 4 |
| Q8CB87 | Rab44 | 1 | Q8BGC0 | Htatsf1 | 2 |
| Q8QZZ8 | Rab38 | 1 | Q6NVF4 | Helb | 4 |
| Q9DCA5 | Brix1 | 1 | Q9CQN7 | Mrpl41 | 3 |
| Q2EMV9 | Parp14 | 3 | Q14C51 | Ptcd3 | 5 |
| Q61263 | Soat1 | 1 | Q8BYK6;P5 | Ythdf3;Ythd | 2 |
| Q8C2A2 | Tsen54 | 1 | P14576 | Srp54 | 6 |
| Q3UYH7;Q | Adrbk2;Adr | 1 | P29391;P49 | Ftl1;Ftl2 | 3 |
| Q9Z2Q2 | Knop1 | 1 | Q8K2T8 | Paf1 | 3 |
| P01757;P01756 | | 1 | P33174 | Kif4 | 2 |
| Q9R0P6 | Sec11a | 1 | Q922J3 | Clip1 | 3 |
| A6PWY4 | Wdr76 | 2 | O08795 | Prkcsh | 3 |
| Q9WVF7 | Pole | 2 | Q922R5;Q6 | Smek2;Sme | 3 |
| E9Q735 | Ube4a | 1 | O35381 | Anp32a | 2 |
| Q8VD65 | Pik3r4 | 1 | Q6ZWM4 | Lsm8 | 2 |
| Q9CRT8 | Xpot | 2 | Q9D554 | Sf3a3 | 5 |
| O08992 | Sdcbp | 1 | Q99KK9 | Hars2 | 4 |
| Q9D125 | Mrps25 | 1 | Q99N89 | Mrpl43 | 2 |
| CON__Q1RMK2 | | 1 | P62320 | Snrpd3 | 2 |
| Q9Z2V5 | Hdac6 | 1 | P18653 | Rps6ka1 | 2 |
| Q14CH7 | Aars2 | 1 | Q9CY52 | Thg1l | 3 |
| Q9CQX2 | Cyb5b | 1 | Q8K2Y7 | Mrpl47 | 5 |
| Q9WTQ8 | Timm23 | 1 | Q9CQE6;Q | Asf1a;Asf1b | 2 |
| Q9DAS9 | Gng12 | 2 | Q9WVG6 | Carm1 | 4 |
| P97370 | Atp1b3 | 1 | Q9ER88 | Dap3 | 3 |
| Q8VDP6 | Cdipt | 1 | Q9CPR5 | Mrpl15 | 3 |
| Q8CI11 | Gnl3 | 1 | P01844;P01 | Iglc2;Iglc3 | 2 |
| O70439 | Stx7 | 1 | Q6PGG2 | Gmip | 4 |
| Q61171 | Prdx2 | 2 | Q8R035 | Ict1 | 2 |
| Q8BGS7 | Cept1 | 1 | P01630 | | 2 |
| P70302 | Stim1 | 1 | Q923G2 | Polr2h | 1 |
| Q8CHI8 | Ep400 | 2 | Q9D8B3 | Chmp4b | 2 |
| Q8K2X3 | Obfc1 | 1 | Q8BHN3 | Ganab | 5 |
| Q02053;P3 | Uba1;Uba1y | 2 | Q9CQF0 | Mrpl11 | 4 |

| | | | | | |
|---|---|---|---|---|---|
| Q9JL26 | Fmnl1 | 1 | O35286 | Dhx15 | 4 |
| Q9Z2Q5 | Mrpl40 | 1 | Q8BK72 | Mrps27 | 3 |
| Q9WTS2 | Fut8 | 1 | P49962 | Srp9 | 2 |
| P54310 | Lipe | 2 | P01592 | Igj | 2 |
| Q99N80 | Sytl1 | 1 | Q6ZWY3 | Rps27l | 1 |
| O70172 | Pip4k2a | 1 | Q99N84 | Mrps18b | 1 |
| Q9EQQ2 | Yipf5 | 1 | P83882 | Rpl36a | 2 |
| Q9CWL8 | Ctnnbl1 | 1 | Q91W50 | Csde1 | 2 |
| Q9CRB2 | Nhp2 | 3 | P01629 | | 1 |
| Q99LQ1 | Mbip | 1 | P01820;P01821;P01819 | | 2 |
| Q9DCT6 | Bap18 | 1 | P08071 | Ltf | 5 |
| Q9Z0V7 | Timm17b | 1 | O54988 | Slk | 5 |
| P11438 | Lamp1 | 1 | P18525 | | 1 |
| Q9DB41;Q9 | Slc25a18;Slc | 1 | P52875 | Tmem165 | 3 |
| P61166 | Tmem258 | 1 | P84244;P84 | H3f3a;Hist | 2 |
| Q8BGH2 | Samm50 | 1 | Q922U1 | Prpf3 | 4 |
| Q61699 | Hsph1 | 1 | Q6ZQ58 | Larp1 | 4 |
| Q07813 | Bax | 1 | Q99N96 | Mrpl1 | 5 |
| Q640N3 | Arhgap30 | 1 | Q8VDT9 | Mrpl50 | 4 |
| Q9JIW9 | Ralb | 1 | O08582 | Gtpbp1 | 4 |
| Q3U186 | Rars2 | 1 | P01634 | | 1 |
| Q8VDL4 | Adpgk | 1 | Q9CY16 | Mrps28 | 4 |
| Q63871 | Polr2k | 1 | Q9D1B9 | Mrpl28 | 3 |
| Q9DC70 | Ndufs7 | 1 | Q8BJZ4 | Mrps35 | 4 |
| Q8JZQ2 | Afg3l2 | 1 | P97822 | Anp32e | 3 |
| Q9CZT5 | Vasn | 1 | P01674 | | 1 |
| Q69ZX6 | Morc2a | 1 | P01635 | | 4 |
| Q9D868 | Ppih | 1 | P51670 | Ccl9 | 2 |
| Q8VE80 | Thoc3 | 1 | P01660 | | 1 |
| Q9Z315 | Sart1 | 1 | Q8BJW6 | Eif2a | 3 |
| Q8BU11 | Tox4 | 1 | CON__Q05B55 | | 4 |
| O35954 | Pitpnm1 | 1 | P01799;P01797;P01796; | | 1 |
| Q56A08 | Gpkow | 1 | Q6ZWU9 | Rps27 | 2 |
| Q99LJ7 | Rcbtb2 | 1 | P01648;P01649 | | 1 |
| Q8BIW9 | Chtf18 | 1 | Q5SUE7 | Adad1 | 1 |
| Q61768 | Kif5b | 1 | P01679;P01678;P01677; | | 2 |
| Q8VEM1 | Rnf130 | 1 | P01657;P01 | Igkv7-33 | 0 |
| Q9CQ92 | Fis1 | 2 | Q91WC9 | Daglb | 1 |
| Q8VCE2 | Gpn1 | 1 | A6H6E2 | Mmrn2 | 1 |
| Q9CQC6 | Bzw1 | 1 | P01633 | Igk-V19-17 | 1 |
| Q9CWH5 | Trmt11 | 2 | P47915 | Rpl29 | 1 |
| Q8K1E0 | Stx5 | 1 | P01806;P01811;P01810; | | 1 |
| Q9EQ28 | Pold3 | 1 | P18528 | | 1 |
| Q8CHY6 | Gatad2a | 1 | | | |
| Q9CYA6 | Zcchc8 | 1 | | | |

| | | |
|---|---|---|
| Q8CB77 | Tceb3 | 1 |
| Q8R1N0 | Znf830 | 1 |
| Q8C0G2 | Traf3ip3 | 1 |
| Q9D198 | Syf2 | 1 |
| Q91VH2 | Snx9 | 1 |
| Q91VM3 | Wdr45 | 1 |
| P36916 | Gnl1 | 1 |
| Q91VJ1 | Aim2 | 1 |
| Q8BMS9 | Rassf2 | 1 |
| Q9R1K9 | Cetn2 | 1 |
| Q8BHB4 | Wdr3 | 1 |
| P16045 | Lgals1 | 2 |
| Q9Z2Y8 | Prosc | 1 |
| Q8BU40 | Nlrp4a | 1 |
| Q02614 | Sap30bp | 1 |
| O35465 | Fkbp8 | 1 |
| Q8JZM0 | Tfb1m | 1 |
| Q6P9L6 | Kif15 | 1 |
| A2A5R2;G3 | Arfgef2;Arf | 1 |
| Q9CQK7 | Rwdd1 | 1 |
| O08900 | Ikzf3 | 1 |
| Q9Z127 | Slc7a5 | 1 |
| Q9CQ22 | Lamtor1 | 1 |
| Q9CXY9 | Pigk | 1 |
| P18572 | Bsg | 1 |
| P49446 | Ptpre | 1 |
| Q8CAS9 | Parp9 | 1 |
| P60898 | Polr2i | 1 |
| O08692 | Ngp | 1 |
| Q9QZ82 | Cyp11a1 | 1 |
| P63213 | Gng2 | 1 |
| Q9JLR9 | Higd1a | 1 |
| Q9CXW3 | Cacybp | 1 |
| Q9CQV4 | Fam134c | 1 |
| Q8K2A7 | Ints10 | 1 |
| Q8R3P6 | Vwa9 | 1 |
| Q6ZQ08 | Cnot1 | 2 |
| O55098 | Stk10 | 1 |
| Q8BRK9 | Man2a2 | 1 |
| Q76KJ5 | Cd3eap | 1 |
| O08915 | Aip | 1 |
| P11928 | Oas1a | 2 |
| Q9WUB0 | Rbck1 | 1 |
| Q6ZQM8 | Ugt1a7c | 1 |
| Q8BMP6 | Acbd3 | 1 |

| | | |
|---|---|---|
| P42230 | Stat5a | 1 |
| P41230 | Kdm5c | 1 |
| Q9Z0F8 | Adam17 | 1 |
| Q91YI4 | Arrb2 | 2 |
| A2A791 | Zmym4 | 1 |
| Q3TC46 | Patl1 | 2 |
| Q9CWY8 | Rnaseh2a | 1 |
| Q9CSN1 | Snw1 | 1 |
| Q3UVL4 | Vps51 | 1 |
| Q5SUQ9 | Ctc1 | 2 |
| Q8CI71 | Ccdc132 | 1 |
| Q3TBW2 | Mrpl10 | 1 |
| P0C7W3;Q8 | Tor2a | 1 |
| P36371 | Tap2 | 1 |
| Q91WA6 | Sharpin | 1 |
| Q68ED3 | Papd5 | 1 |
| O88668 | Creg1 | 1 |
| Q3UMU9 | Hdgfrp2 | 1 |
| O88379 | Baz1a | 2 |
| A2AIV2 | Kiaa1429 | 2 |
| Q91UZ5 | Impa2 | 2 |
| Q6PD26 | Pigs | 1 |
| Q8C4J7 | Tbl3 | 1 |
| Q6NZQ4 | Paxip1 | 1 |
| P60060 | Sec61g | 1 |
| Q9WV80 | Snx1 | 1 |
| Q9CZW4 | Acsl3 | 2 |
| Q3UMQ8 | Naf1 | 1 |
| Q3U2S8 | Hvcn1 | 1 |
| P49135 | Ercc3 | 1 |
| Q9QZH6 | Ecsit | 1 |
| Q8VI75 | Ipo4 | 1 |
| Q8R5K4 | Nol6 | 2 |
| Q5DTY9;Q5 | Kctd16;Kctd | 1 |
| Q61048 | Wbp4 | 1 |
| Q3UPH1 | Prrc1 | 1 |
| P56183 | Rrp1 | 1 |
| O08789 | Mnt | 1 |
| Q3UB74 | Tbrg1 | 1 |
| Q9DCR2 | Ap3s1 | 1 |
| Q811J3 | Ireb2 | 1 |
| Q9D174 | Ss18l2 | 1 |
| P61924 | Copz1 | 1 |
| Q80WQ2 | Vac14 | 1 |
| Q9WVL2 | Stat2 | 1 |

| | | |
|---|---|---|
| Q920I9 | Wdr7 | 1 |
| P35123 | Usp4 | 1 |
| Q9JLC8 | Sacs | 1 |
| Q9EQQ9 | Mgea5 | 1 |
| O35387 | Hax1 | 1 |
| Q8K4F6 | Nsun5 | 1 |
| B1AUH1 | Ptpru | 1 |
| O08848 | Trove2 | 1 |
| O70481 | Ubr1 | 1 |
| O88455 | Dhcr7 | 1 |
| P01798 | | 1 |
| P08249 | Mdh2 | 1 |
| P27612 | Plaa | 1 |
| P30355 | Alox5ap | 1 |
| P46467 | Vps4b | 1 |
| P46718 | Pdcd2 | 1 |
| P50518 | Atp6v1e1 | 1 |
| P52912 | Tia1 | 1 |
| P53612 | Rabggtb | 1 |
| P62046 | Lrch1 | 2 |
| Q149F1 | Rpusd2 | 1 |
| Q2VPQ9 | Meaf6 | 1 |
| Q497V5 | Srbd1 | 1 |
| Q6P9R1 | Ddx51 | 1 |
| Q8BJ71 | Nup93 | 2 |
| Q8BVK9 | Sp110 | 1 |
| Q8BXA1 | Golim4 | 1 |
| Q8BZ20 | Parp12 | 2 |
| Q8CGC6 | Rbm28 | 3 |
| Q8CGY8 | Ogt | 1 |
| Q8K3J1 | Ndufs8 | 1 |
| Q8R3H7 | Hs2st1 | 2 |
| Q91YY4 | Atpaf2 | 1 |
| Q922S8;Q8 | Kif2c;Kif2b | 2 |
| Q99J47 | Dhrs7b | 1 |
| Q99KG3 | Rbm10 | 1 |
| Q99N93 | Mrpl16 | 1 |
| Q99PT1 | Arhgdia | 1 |
| Q9CQB5 | Cisd2 | 1 |
| Q9CXJ1 | Ears2 | 1 |
| Q9D0D4 | Dimt1 | 2 |
| Q9D937 | | 1 |
| Q9DAM7 | Tmem263 | 1 |
| Q9JI90 | Rnf14 | 1 |
| Q9JKX6 | Nudt5 | 1 |

| | | |
|---|---|---|
| Q9R0N0 | Galk1 | 1 |
| Q9Z2Z6 | Slc25a20 | 2 |

**TABLE S5**

**CALM-AF10 RNA-seq targets + protein interactors**

**Gene Name**
1190002N15RIK
2900026A02RIK
4921529L05Rik
4933430I17RIK
6030419C18RIK
A2M
A430078G23RIK
A630033H20RIK
AA467197
ABCA1
ABCA4
ABCB1B
ABCB4
ACTR3
ADAM8
ADGRD1
ADGRG1
ADGRL4
AFP
AHSG
AKAP2
ALDH3A1
ALDOC
ALG11
AMOT
ANGPT1
APBB2
APLN
ARG1
ARGLU1
ARHGAP32
ARHGEF10L
ARHGEF18
ARMCX1
ARPC1B
ARPC2
ARPC3
ARPC4
ARPC5
ATP6V0D2

B3GLCT
B3GNT5
BAHCC1
BEND4
BHLHA15
BHLHE41
BLOC1S2
BMI1
BNIP3
BTLA
C030034L19Rik
C130046K22Rik
C1QA
C1QC
C1QTNF1
C2CD4A
C4B
C530008M17RIK
CALCRL
CAMK2A
CAPN5
CAR13
CAR3
CARNS1
CASC4
CASP12
CAST
CCDC112
CCDC27
CCDC88C
CCL2
CCL3
CCL4
CCL7
CCND1
CCR1
CD27
CD274
CD28
CD2AP
CD33
CD34
CD36
CD72
CD93

CD96
CDH2
CDK14
CDK17
CDKN2C
CFAP57
CGNL1
Chd3os
CIR1
CLNK
CLSTN3
CNKSR3
COL4A1
COL4A2
CORO1A
CORO1B
CPNE7
Creb5
CREBRF
CRISP3
CSF1
CSGALNACT1
CTLA2A
CTLA2B
CX3CR1
CXCL2
CYSLTR2
D8ERTD82E
DAB2
DCBLD2
DDX58
DHX58
DNAJC10
DOCK9
DOK2
DPY19L1
DSTN
DUSP3
DUSP4
ECHS1
EGLN3
EHD3
EIF2AK2
EIF2AK3
EMP1

ENDOD1
ENDOU
ENPP1
ERG
ETNK1
EYA1
F13A1
F2RL3
FADS3
FAM198B
FAM20A
FAM26F
FAM84B
FGF3
FLT3
FNBP1L
FRMD5
FSD1L
FSTL1
GAS2L3
Gbp11
GBP6
GCNT4
GDF15
GFI1B
GGACT
GIMAP5
GIMAP6
GJA5
Gm12250
Gm15915
GM5111
Gm6093
GM6377
GMFG
GPATCH1
GPD1
GPNMB
GPR84
GSN
GZMB
HBB-BT
HBEGF
HCAR2
HELZ2

HID1
HIVEP3
HLF
HMGA2
HMGN3
HOXA10
HOXA2
HOXA3
HOXA5
HOXA7
HOXA9
HP
HPGD
HRH2
HSH2D
HTRA3
ICOS
IFI203
IFI27
IFI47
IFIT1
IGF2BP2
IL18RAP
IL1R1
IL1RN
IL2RA
IL5RA
IL7R
IL9R
INSIG1
IPCEF1
IPO8
IRF7
IRF9
IRGM1
ISG15
ITIH5
JAG2
JAK1
JAK3
JUN
KBTBD11
KCNH2
KCTD5
KIF20B

KIF5A
KLHL24
LAT2
LEPR
LGALS3BP
LHFPL2
Lincred1
LPAR1
LPL
LRRC32
LRRFIP2
LUZP1
LY6A
MAFF
MAP7
MAP9
MAST4
ME1
MECOM
MEIS1
MGARP
MGAT4A
Mir155hg
Mira
MLLT10
MMP12
MMP13
MMP8
MOB2
Morc3
MOV10
MRVI1
MSI2
MTHFD1L
MUC13
Mx1
MYCN
MYCT1
MYH10
MYL1
MYL3
MYO10
MYO18A
MYO1E
MYO1G

NANOS1
NEURL1B
NKX2-3
NLRP1A
NR1D2
NRGN
NSF
NT5E
OAS1A
Oas1b
OAS1C
OAS2
OAS3
OASL1
OASL2
OXCT1
P4HA2
PARP12
PCP4L1
PDCD1LG2
PDE11A
PEA15A
PEAR1
PFKFB1
PGLYRP2
PHF11B
PIAS1
PICALM
Pim2
PIP5K1C
PIWIL2
PLA2G4C
PLBD2
PLEK
PLEKHA6
PLEKHA7
PLG
PLK2
PLOD2
PLS1
PLVAP
PMP22
PNRC1
PPIG
PPP1R7

PRDX1
PRDX4
PRKCQ
PROCR
PRRG4
PTGS2
Ptgs2os2
PTPRCAP
QPCTL
Raph1
Rasgef1b
RASGRP3
RASL11A
RB1CC1
RCOR2
RFTN1
RGL1
RGS1
RGS11
RHOJ
RINL
RSAD2
RTP4
SAMD9L
SAMSN1
SAP18
SASH1
SCIMP
SCIN
SCML4
SCN3A
SCYL2
SDC1
SDC3
SEL1L3
SELENBP1
SEMA7A
Serpina3g
SERPINE2
SETX
SGK1
SGSH
SHANK3
SIX1
SKIDA1

SLC13A2
SLC14A1
SLC17A8
SLC18A2
SLC22A23
SLC22A3
SLC26A9
SLC29A3
SLC30A4
SLC35F2
SLC39A10
SLC4A8
SLFN5
SLFN8
SOX6
SPP1
SPRED1
SPSB4
SSBP2
ST3GAL1
ST8SIA4
STXBP6
SULF2
TBXA2R
TCEA2
TCF4
TCTEX1D1
TFR2
TFRC
TGM2
TGTP1
TIE1
TIMP3
TIPARP
TLE4
TLR12
TLR13
TLR7
TMEM215
TMEM41B
TMOD3
TNFAIP3
TNIP3
TNS1
TOX

TRF
TRIM30A
TRIM35
TRPC6
TSC22D1
TSPAN13
TTLL7
TYMS
UNC93B1
USP18
VASH2
VLDLR
WDFY1
WDR1
WFDC17
WNT9B
XAF1
ZBTB6
ZFP760
ZFP882
